# Identifying phage proteins that activate the bacterial innate immune system

**DOI:** 10.1101/2025.07.02.662641

**Authors:** Toni A. Nagy, Gina W. Gersabeck, Amy N. Conte, Aaron T. Whiteley

## Abstract

Bacteria have evolved sophisticated antiphage systems that halt phage replication upon detecting specific phage triggers. Identifying phage triggers is crucial to our understanding of immune signaling, however, they are challenging to predict. Here we used an expansive plasmid library that expressed 400 phage protein-coding genes from 6 different phages to identify novel triggers of known and undiscovered antiphage systems. We transformed our library into 72 diverse strains of *E. coli*. Each strain natively harbors a different suite of antiphage systems whose activation typically inhibits growth. By tracking plasmids that were selectively depleted, we identified over 100 candidate phage trigger-*E. coli* pairs. Two phage trigger proteins were investigated in detail, revealing a novel antiphage system that detects multiple phage tail fiber proteins and identifying major capsid protein as the activating ligand of the antiphage system Avs8. These experiments provide a unique dataset for continued definition of the molecular details of the bacterial immune system.

## Introduction

Bacteria and bacteriophages (phages) have been coevolving for over a billion years and are often in direct competition with each other^1,2^. Phages hijack host machinery to repurpose these proteins for their benefit and, in turn, bacteria counter phage using antiphage systems that halt infection. Antiphage systems are suites of genes, often encoded in an operon or a single large gene, that sense phage infection and limit virion production. Antiphage systems were previously thought to be limited to restriction modification and CRISPR-Cas systems, however, we now appreciate there are over 100 distinct antiphage systems encoded by bacteria that quickly sense and respond to infection^3–7^. Cataloging these discoveries has enabled accurate predictions of defense systems from genome sequence^8–10^. On average, each bacterial strain only encodes 5.8 antiphage systems, however the identity of the antiphage systems within each strain varies from strain-to-strain^4^. In addition, antiphage systems are typically encoded in mobile genetic elements, which can be rapidly exchanged with other bacteria. In this way, bacteria distribute phage defense systems in the pangenome^1^ and maintain the upper hand against phages that must adapt to each new strain’s resident antiphage systems.

Each phage defense system conforms to a general signaling scheme: a sensor detects a phage trigger, a signal amplifier/transducer relays the signal and sets a threshold for signaling, and an effector stops virion production^11^. Conserved protein domains identified within each component help generate hypotheses for the signaling and effector mechanism. However, predicting the phage stimuli that activate sensor domains is rarely possible based on sequence information alone because the protein folds in these domains are uninformative, for example, tandem repeat domains such as leucine-rich repeats, non-conserved regions, or domains of unknown function. Further, classic genetic techniques such as selecting for phage escaper mutants that evade detection are complicated by phage stimuli that are essential for replication or abundant phage-encoded protein inhibitors of antiphage systems, making the question of how antiphage systems sense infection elusive.

Previous studies show that antiphage systems can detect proteins and their functions, nucleic acids and their modifications, and even changes in the cell metabolome^12^. The coevolution between phages and bacteria has provided a particularly rich opportunity to uncover novel protein-protein interactions, as these interactions are high affinity and sometimes mediated by the protein fold (as opposed to amino acid sequence)^13^. Here we sought to map the landscape of phage proteins that activate the bacterial immune system by flipping the paradigm of how antiphage systems are investigated. Rather than focusing on a single system and asking how it is activated, we generated a library of candidate phage triggers from phage open reading frames (ORFs) and surveyed the pangenome for immune activation by transforming our ORF library into diverse strains of bacteria, each with different resident defense systems. Signature tagged mutagenesis then identified the specific phage defense system responsible for immune activation. Our analysis led to discovery of a new defense system, the activator of a previously discovered system, and a wealth of new phage triggers for future investigations.

## Results

### Construction of an extensive phage ORF library

We began by constructing a library of plasmids expressing every open reading frame (ORF) from phages T2, T7, λ, MS2, ϕX174, and M13 using a medium-copy vector with an IPTG-inducible promoter (**Fig. 1a**). These phages were selected because they represent well-studied dsDNA (T2, T7, λ), ssDNA (ϕX174, M13), and ssRNA (MS2) genomes. Plasmids were cloned *en mass* and verified by next generation sequencing, see Methods section for full details. We successfully constructed vectors expressing 406 of the 414 identified ORFs; the remaining eight could not be constructed, likely because these ORFs were too toxic for vector construction even under non-inducing conditions (**Table S1**).

**Figure 1.**
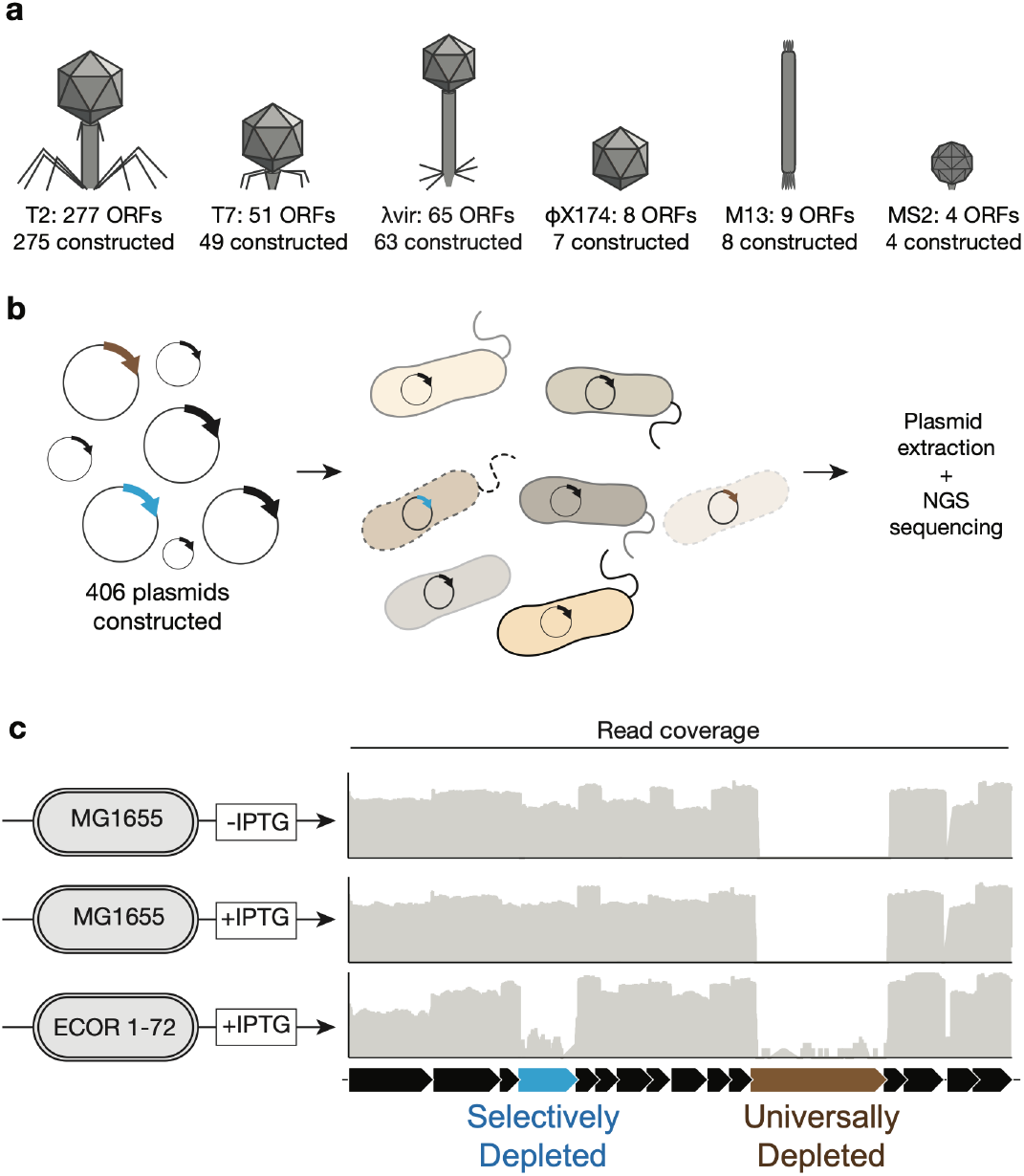
A genetic screen to identify candidate phage triggers. **(a)** Plasmids expressing every phage open reading frame (ORF) predicted to encode a protein were constructed. Phage ORFs from the genomes of dsDNA phages (T2, T7, λ), ssDNA phages (φX174, M13), and a ssRNA phage (MS2) were expressed under the control of an IPTG-inducible promoter. Plasmids for 406 out of a total of 414 ORFs could be constructed. **(b)** Individual plasmids were pooled and transformed into bacterial strains of interest. By transforming K-12 *E. coli* (strain MG1655), proteins that are generally growth inhibitory could be identified. By transforming diverse *E. coli* from the ECOR collection, proteins that are growth inhibitory to specific strains could be identified. **(c)** Transformants were recovered on solid agar with or without 50 µM IPTG, plasmids were extracted from the pooled population, sequenced, and reads mapped to phage genomes. Representative data shows read depth on a log scale for reads =aligned to a portion of the phage λ genome.

We pooled our plasmid library expressing 406 phage ORFs along with a version of the vector that expressed a neutral protein, GFP, and transformed the pool into K-12 *E. coli* strain MG1655 (**Fig. 1b**). Bacteria were plated on solid agar with 50 µM IPTG to induce phage ORF expression or without inducer. Plasmids were extracted from surviving bacteria, deep sequenced, and the reads were mapped to phage genomes to quantify each plasmid (**Fig. 1b**). We first assessed the raw read count across phage genomes and found that 41 phage ORFs were poorly recoverable when transformed into MG1655 under non-inducing conditions, defined as having less than 10 reads covering the ORF (**Table S2A, B**). These ORFs likely inhibited growth even at low, uninduced levels, but were originally able to be cloned likely because the strain of *E. coli* used to construct the library expressed additional copies of the lac repressor.

To compare plasmid coverage across different data sets, we normalized read counts using transcripts per million (TPM) scores for each ORF (**Table S2C**). Using a threshold of a 10-fold decrease in TPM compared to TPM in non-inducing conditions, an additional 28 phage ORFs were depleted when MG1655 transformants were cultivated under inducing conditions (**Table S2D, E**). ORFs depleted from MG1655 under any condition were excluded from further analysis as these appeared to generally inhibit growth of *E. coli*, leaving 337 phage ORFs that could be tested as potential triggers of the bacterial immune system (83% of total, **Table S2F**).

### A large-scale screen for phage triggers

We next determined genetic interactions between specific phage ORFs and diverse *E. coli* strains by transforming our pooled plasmid library into 72 *E. coli* strains from the *E. coli* reference (ECOR) collection^14^ and monitoring plasmid abundance. The majority of antiphage systems stop virion production by “abortive infection”, a general term for when a phage defense system stops virion production but does not allow the infected bacterium to survive^2,11^. Many abortive infection antiphage systems cause damage to the host cell (e.g., nucleases that destroy phage and host genomes or NADases that depleting essential metabolites)^15^. Therefore, activation of an antiphage system by a phage ORF is likely to inhibit growth of the bacterium and plasmids expressing that ORF would be depleted from the population.

Each ECOR strain is “wild” compared to the domesticated K-12 lab strains that have lost many mobile genetic elements. Profiling how phage ORFs impact growth of ECOR strains therefore provides a method to survey otherwise unavailable segments of the *E. coli* pangenome. Genome analysis confirmed that each ECOR strain encodes a different arsenal of ∼5–20 known antiphage systems and the 72 ECOR strains represent a total of >60 distinct systems (**Table S3**, see Methods).

Our screen revealed greater than 100 trigger-ECOR pairs: candidate triggers of antiphage systems that selectively inhibited growth of specific ECOR strains (**Figure 2 and Table S2F-H**). Unexpectantly, 22 ECOR strains were poorly transformable and we were only able to recover a few colonies and an additional 11 produced poor-quality sequencing for unknown reasons; these strains were not investigated further (**Table S4**).

**Figure 2.**
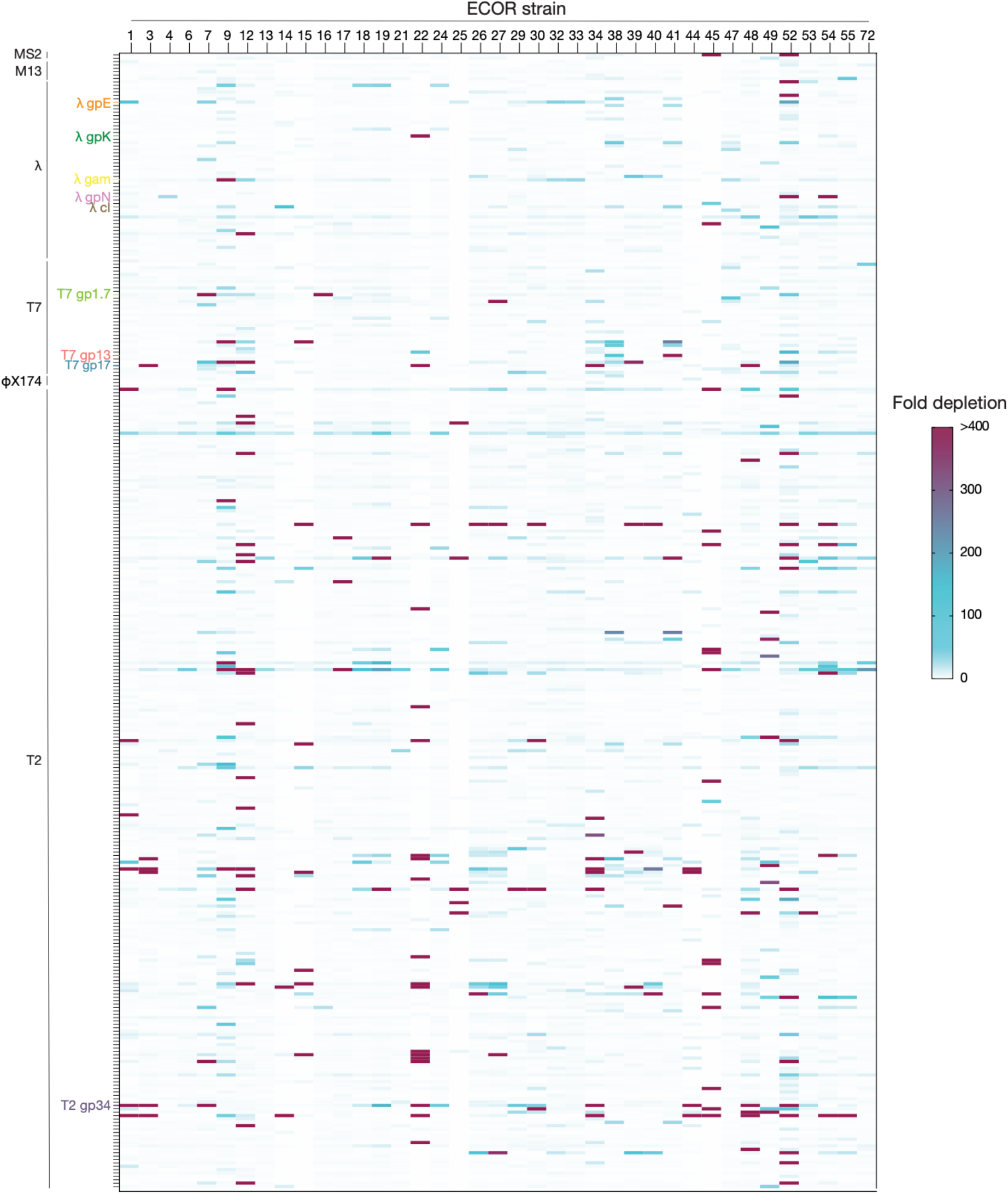
Phage ORFs selectively depleted in ECOR strains. Our expansive library of plasmids expressing phage ORFs was transformed into diverse E. coli and the frequency with which each plasmid could be recovered was calculated by deep sequencing. Read coverage of each ORF was compared as transcripts per million (TPM) and expressed as a fold depletion compared to uninduced MG1655 (see Methods for full details). The y-axis is each phage ORF analyzed and grouped for each phage, as labeled. Specific phage ORFs analyzed in Fig. 3 are labeled. The x-axis is each ECOR strain analyzed and labeled for each ECOR strain number.

We validated a subset of our findings by comparing the number of colonies recovered when MG1655 or an ECOR strain was transformed with a single plasmid expressing the candidate phage trigger (**Fig. 3, ED Fig.1**). We focused on ORFs that we hypothesized were ideal targets of an innate immune system: (*i*) structural proteins such as major capsid proteins (e.g., λ gpE) and tail fibers (e.g., T7 gp17, T2 gp34, λ gpK) that exhibit structural conservation, (*ii*) proteins that are required for the successful replication or lysogeny of phage (e.g., λ cI, λ gpN, T7 gp1.7), (*iii*) an internal virion protein of T7 (T7 gp13), which is the first protein that enters the bacterium upon infection^16,17^, or (*iv*) known inhibitors of phage defense systems (e.g., λ gam).

**Figure 3.**
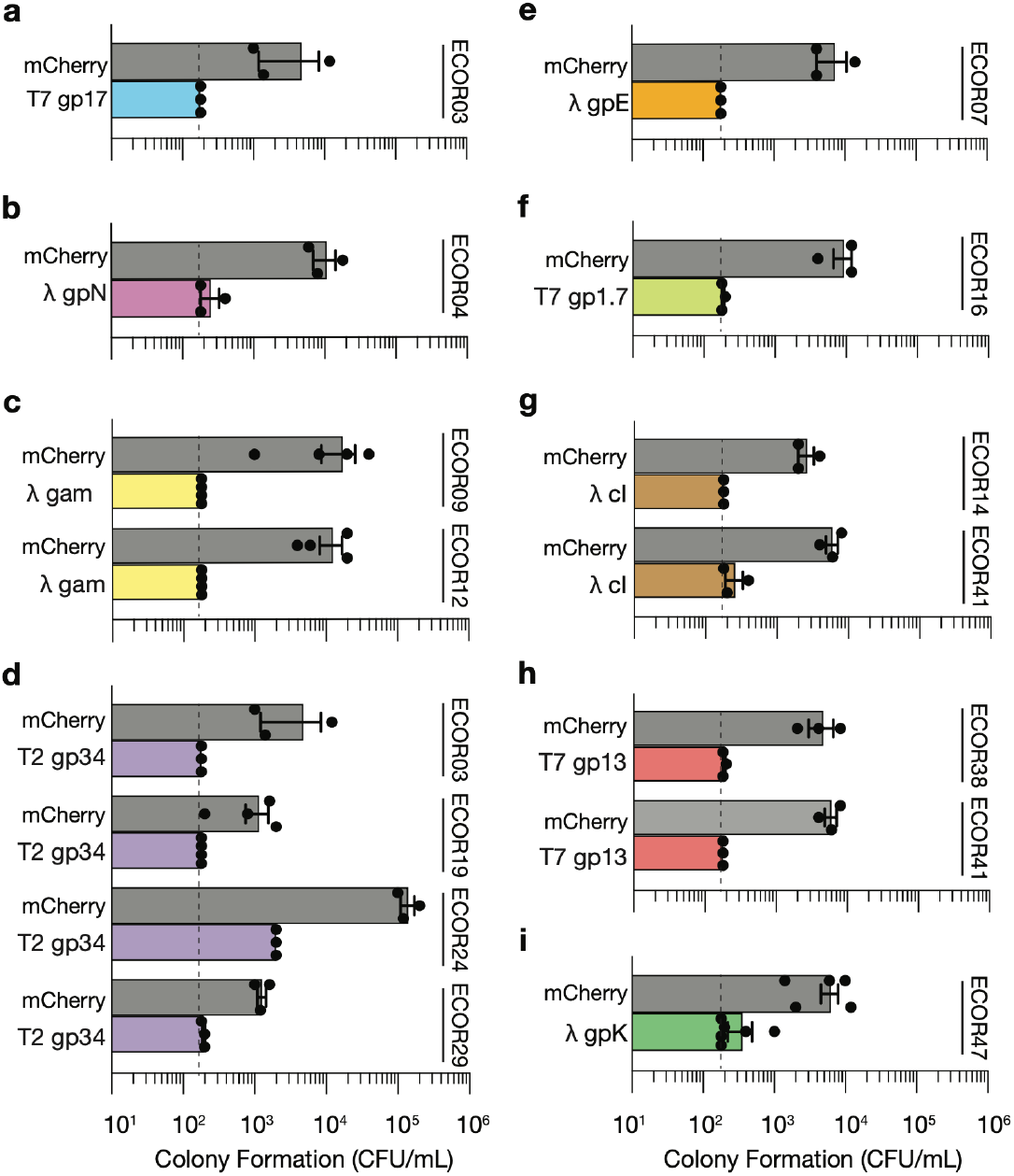
Diverse phage proteins inhibit growth of specific wild strains of *E. coli* but not MG1655. Number of colonies of MG1655 or the indicated ECOR strain recovered when transformed with a plasmid expressing the indicated phage ORF and grown in medium containing 50 µM IPTG. The maximum recovered bacteria reflects the electroporation efficiency/bacterial competence for that strain, which can be compared using the mCherry neutral protein control. Data are the mean ± standard error of the mean (SEM) of n≥3 biological replicates.

We have used conventional gene and gene product names to describe our results; however, genome annotations can produce conflicting nomenclature. See the Methods section for accession numbers of molecules appearing in this study and detailed locus tags in Table S1 for unambiguous descriptions. Transformation efficiency experiments validated results from the screen and demonstrated robust selective depletion of phage ORFs in ECOR strains compared to in MG1655 (**Fig. 3, ED Fig.1**).

### A novel defense system is activated by multiple phage proteins

We first analyzed the genetic interaction between ECOR03 and T7 gp17, an essential phage protein that trimerizes to form each individual tail fiber on the T7 virion^18^. We searched for mutations in ECOR03 that disrupted T7 gp17-mediated growth inhibition. A transposon mutant library was constructed in ECOR03, these strains were transformed with T7 gp17, and surviving transformants were recovered from inducing medium. The majority of ECOR03 mutants are not expected to alter T7 gp17-mediated growth inhibition and therefore do not grow. Sequencing the transposon insertion site of mutants that did grow revealed 28 out of 48 mutants contained transposons that disrupted a two-gene operon of unknown function (**Table S5**). We gave the genes in this operon the placeholder names of “*gene A*” and “*gene B*” and their encoded proteins can be found in genomes throughout the Enterobacteriaceae (**Fig. 4a**). Sequence analysis of these proteins was largely uninformative: A has a single predicted transmembrane domain with no similarity/homology to described proteins, B also has no similarity/homology to described proteins (**Fig. 4a, ED Fig. 2**). Structure predictions further failed to detect protein interactions between Genes A, B, and T7 gp17 (ipTM scores of each pair or all three ≤0.5, **Supplementary File 1**).

**Figure 4.**
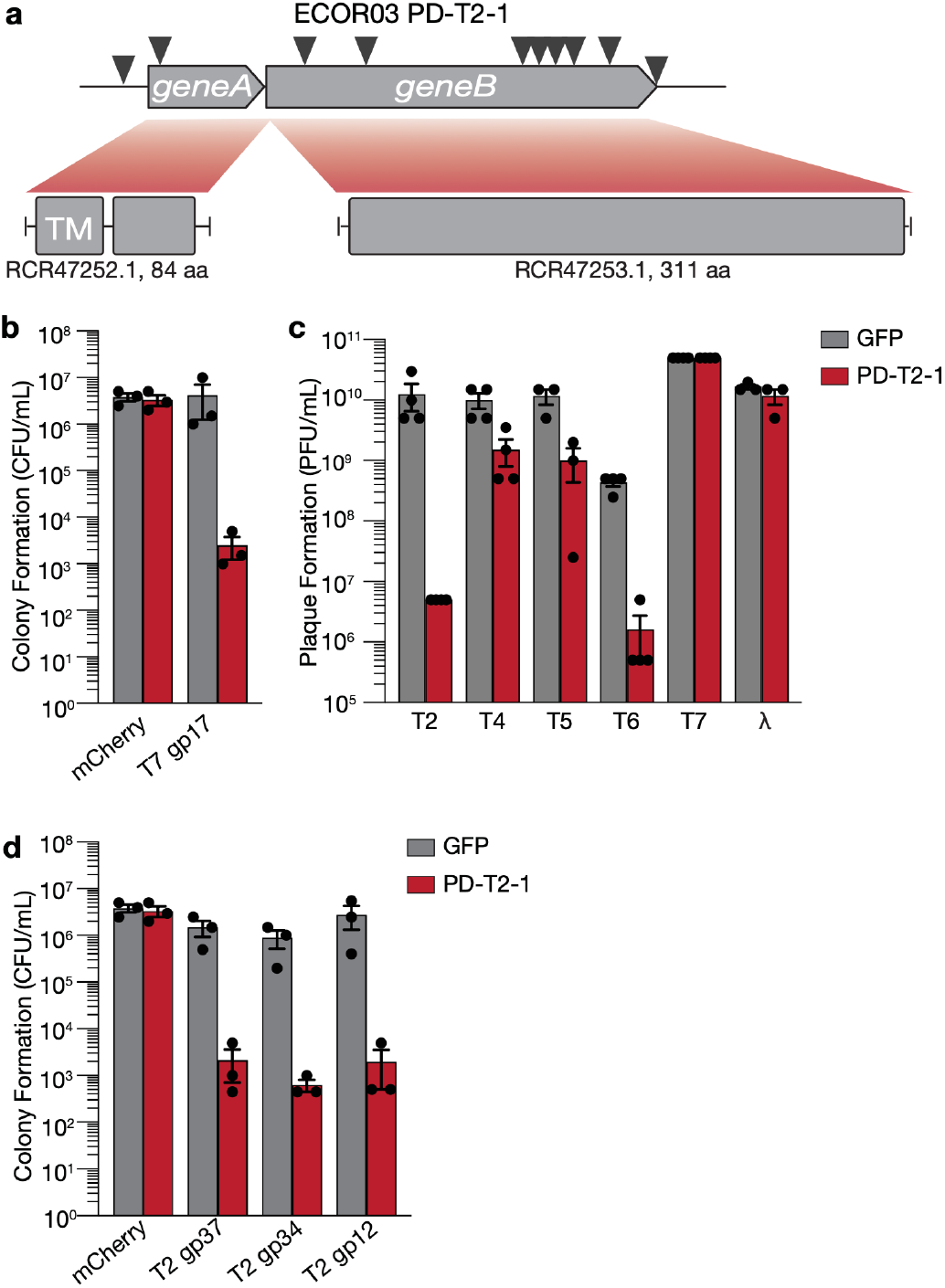
PD-T2-1 is an antiphage system that is activated by multiple phage ORFs. **(a)** Operon structure of <u>p</u>hage defense against <u>T2</u> system 1 (PD-T2-1). Transposon insertions identified in mutant ECOR03 expressing T7 gp17 are indicated with triangles. A transmembrane domain (TM) is predicted for PD-T2-1A and the length of each gene in amino acids (aa) is shown. (**b**) Transformation efficiency of MG1655 expressing GFP or PD-T2-1 transformed with mCherry or select phage ORFs and grown in medium containing 50 µM IPTG. (**c**) Efficiency of plating for the indicated phages infecting MG1655 expressing GFP or PD-T2-1. For b–d, data are the mean ± SEM of n≥3 biological replicates.

We investigated *geneAB* by expressing the two gene operon under the control of its native promoter in MG1655. This approach helps to limit effects of other identified and unidentified antiphage systems also resident in ECOR03 that could confound our analysis. We first confirmed *geneAB* could respond to T7 gp17 (**Fig. 4b**) and found T7 gp17 inhibited colony formation only when transformed into bacteria expressing *geneAB*. Next, we tested whether *geneAB* was indeed a bona fide antiphage system by challenging bacteria expressing *geneAB* with diverse phages (**Fig. 4c** and **ED Fig. 1**). *geneAB* conferred >100-fold protection against phages T2 and T6, and about 10-fold protection against phages T4 and T5 (**Fig. 4c, ED Fig. 1**). Intriguingly, we saw no protection against phage T7, despite robust growth inhibition when T7 gp17 was co-expressed with *geneAB*. We speculate that phage T7 may encode an inhibiting protein, however, we have not investigated this idea further. These findings led us to rename this operon <u>p</u>hage <u>d</u>efense against <u>T2</u> system <u>1</u> (PD-T2-1), adopting the provisional nomenclature introduced by Vassallo *et al*^7^. We invite the renaming of these genes and system upon determination of their defense mechanism.

We hypothesized that PD-T2-1 must recognize additional phage triggers that are produced by phage T2 and returned to our original dataset of proteins that selectively inhibited ECOR03 growth. Genes encoding T2 gp37 (large distal tail fiber subunit), T2 gp34 (proximal tail fiber subunit), and T2 gp12 (short tail fiber) were also depleted and share gross similarity to T7 gp17. Transformation assays revealed that PD-T2-1 also selectively inhibited bacterial growth when T2 gp37, T2 gp34, and T2 gp12 were co-expressed (**Fig. 4d**). These results suggest that PD-T2-1 may recognize a shared feature of these proteins or perhaps these ORFs are indirectly sensed through a shared host intermediate.

### Avs8 is activated by major capsid proteins

We next characterized the genetic interaction between ECOR07 and λ gpE, which encodes the phage major capsid protein^19^. Transposon mutagenesis of ECOR07 followed by transformation with a plasmid expressing λ gpE selected for mutants able to survive gpE expression. Transposon insertion site sequencing revealed that 10 out of 24 mutants had a transposon in the previously identified antiphage system gene PD-λ-4A^7^ (**Table S6, Fig. 5a**). Sequence analysis shows that PD-λ-4A is a STAND NTPase and a member of the antiviral ATPases/NTPases of the STAND superfamily (AVAST) system^5,13^ (Alex Gao, personal communication). However, this protein is distinct from previously identified Avs proteins and we therefore renamed PD-λ-4 as Avs8. The Avs8A protein is highly conserved and homologs sharing >80% identity are abundant across Gammaproteobacteria. Vassallo *et al*. defined this Avs8 antiphage system as a two-gene operon made up Avs8A and Avs8B. Avs8A contains a Mrr nuclease domain and a P-loop NTPase^7^. Avs8A is encoded in an operon with a gene encoding Avs8B, which had no similarity/homology to described proteins in our analysis (**Fig. 5a, ED Fig. 3**). AlphaFold3 structure modeling predicted Avs8A forms a tetramer (ipTM = 0.64), with lower confidence predictions for trimer (ipTM = 0.34) or dimer (ipTM = 0.20) formation (**ED Fig. 3, Supplementary File 1**). These results are similar to the observed active forms of antiviral ATPase/NTPase of the STAND superfamily (AVAST) system members Avs3 and Avs4, which form homotetramers and defend against phage^13^. However, structure modeling is not sufficient to determine if Avs8A is a stable tetramer in the inactive form or requires activation for oligomerization.

**Figure 5.**
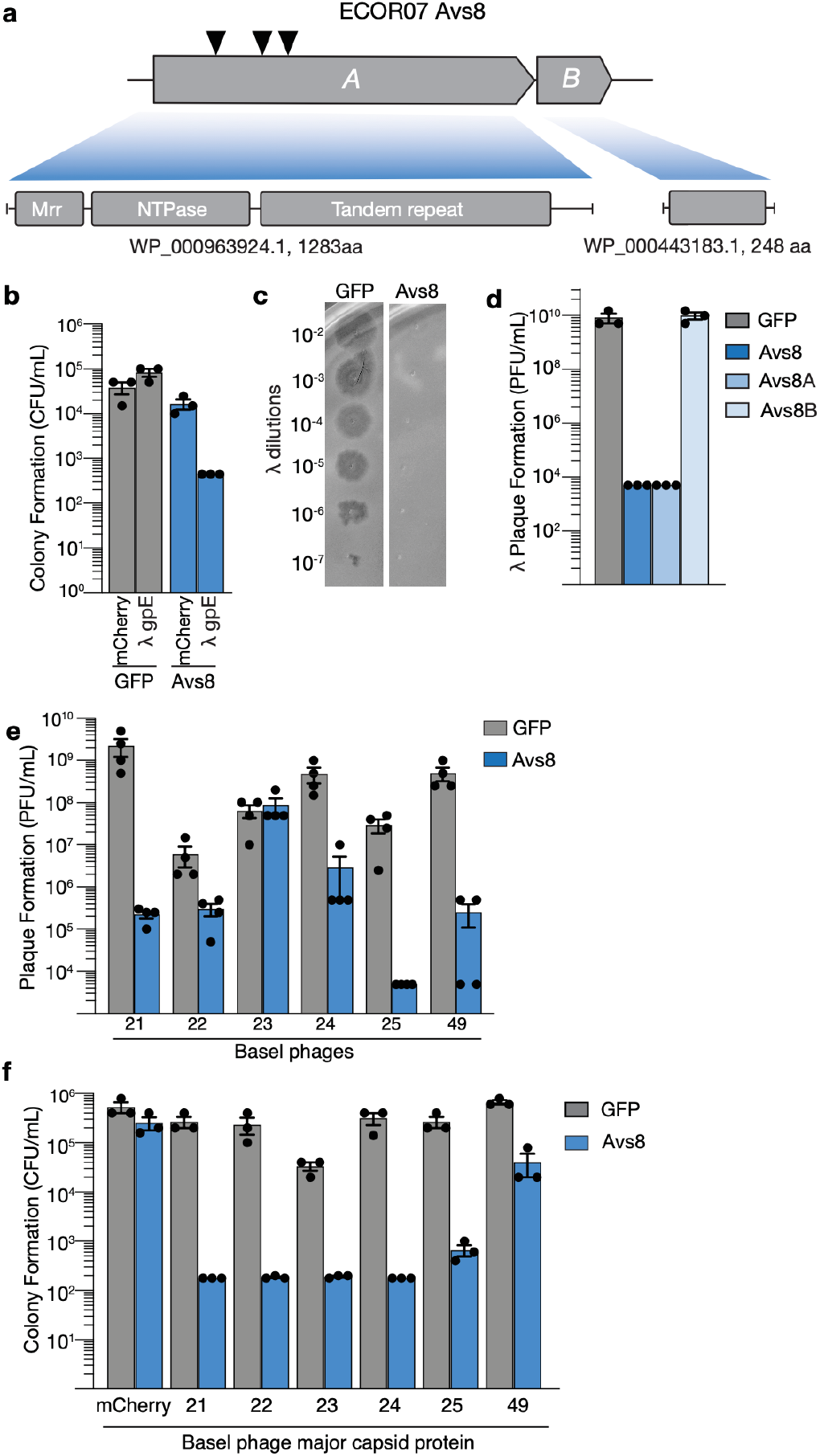
Avs8 is activated by λ gpE and other HK97 fold proteins. **(a)** Operon architecture of Avs8. Transposon insertions are indicated with arrow heads. Mrr nuclease, and P-loop NTPase domains are indicated. **(b)** Quantification of recovered transformants of MG1655 expressing GFP or Avs8 transformed with mCherry or λ gpE and plated on 50 µM IPTG. **(c-d)** Efficiency of plating of phage λ*vir* infecting MG1655 expressing GFP or the indicated genotypes of Avs8. (**e**) Efficiency of plating of the indicated Basel phages infecting MG1655 expressing GFP or Avs8. **(f)** Quantification of recovered transformants of MG1655 expressing GFP or Avs8 transformed with mCherry or major capsid protein from indicated Basel phages and plated on 50 µM IPTG. For b, d, e, and f, data are the mean ± SEM of n≥3 biological replicates. For c, data are representative of n=3 biological replicates.

We investigated Avs8 by expressing the full operon under the control of its native promoter in MG1655. Strains expressing Avs8 could not be transformed with a plasmid expressing gpE, however, gpE-expressing plasmid was readily transformed into bacteria with an empty vector (**Fig. 5b**). Consistent with previous analysis, Avs8 conferred robust protection against phage λ (**Fig. 5c**). These data show that gpE is a trigger for Avs8 and suggest that λ defense is mediated by detection of the major capsid protein.

We measured the range of phage defense provided by Avs8 by analyzing defense against the BASEL collection^20^ of phages. Avs8 also provided defense against Bas21-25, Bas46, and Bas49 (**Fig. 5c, ED Fig. 1, Fig. 5e**). The major capsid proteins encoded by these phages share low sequence identity (**Table S7**) but all adopt an HK97 protein fold. Transformation efficiency assays revealed that plasmids expressing the major capsid protein from Bas21-25 phages could not be expressed in bacteria expressing Avs8 (**Fig. 5f**), similar to λ gpE.

### Avs8A recognizes diverse HK97 fold proteins

The mechanism of Avs8-mediated phage protection was investigated in detail by first determining which of the components of the two-gene operon were required. Avs8A provided equivalent phage λ defense to the complete Avs8 system but Avs8B was dispensable and did not provide defense in the absence of Avs8A (**Fig. 5d**). These results are consistent with our transposon mutagenesis, which only recovered mutations in Avs8A, not Avs8B (**Fig. 5d and Table S6**).

We hypothesized that Avs8A directly binds to major capsid proteins and subsequently activates the N-terminal effector domain to halt virion production. We tested this by first measuring the affinity of Avs8A for major capsid proteins. Recombinant Avs8A bound to λ gpE with an apparent *K*_*D*_ of 127± 23 nM and bound to Bas21 major capsid protein with an apparent *K*_*D*_ of 14 ± 6 nM (**Fig. 6 a, b and ED Fig. 4**). AlphaFold 3 protein structure predictions further supported these interactions, predicting complex formation between Avs8A and gpE with an ipTM = 0.63 and complex formation between Avs8A and Bas21 major capsid protein with an ipTM = 0.79 (**ED Fig. 4**). Additional Alphafold predictions of HK97 fold-containing major capsid proteins from diverse phages retrieves ipTM scores > 0.6, supporting that Avs8A may also recognize more proteins (**Fig. 6g**). This is in stark contrast to ipTM scores of < 0.3 when modeling Avs8A interacting with major capsid/coat protein monomers with non-HK97 folds: MS2 and Qβ^21–23^; φX174 jelly-roll fold^24^; and M13^25^.

**Figure 6.**
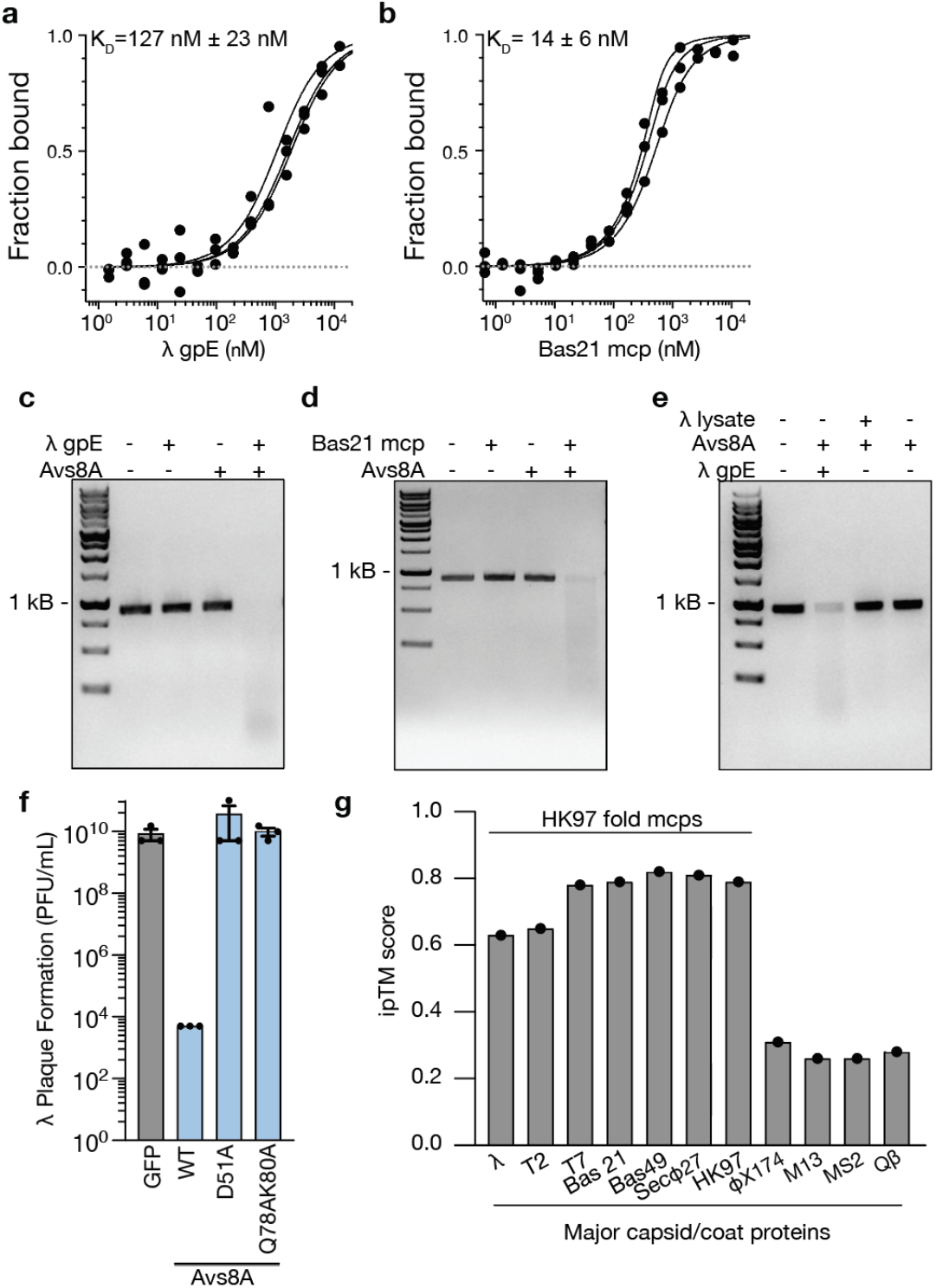
Major capsid proteins bind to Avs8-A to activate DNase activity. (**a–b**) Microscale thermophoresis measurement of interactions between purified his-tagged Avs8A and λ gpE or Avs8A and Bas21 major capsid protein (mcp). Data are the best fit lines for each of three biological replicates and the K_D_ is the average ± SEM for those replicates. (**c–e**) Agarose gel electrophoresis of linear dsDNA substrate incubated with Avs8A, λ gpE, Bas21 mcp, or λ virions as indicated. Data are representative of n=3 biological replicates. (**f**) Efficiency of plating of phage λ*vir* infecting MG1655 expressing GFP or the indicated genotypes of Avs8A. Data are the mean ± SEM of n=3 biological replicates. (**g**) Alphafold3 ipTM scores of HK97 fold and non-HK97 fold major capsid proteins predicted with Avs8A.

The N-terminus of Avs8A contains a predicted Mrr-family nuclease domain which is part of the larger PD-(D/E)xK nuclease-like superfamily of domains. In related antiphage systems, these domains degrade phage and bacterial DNA upon activation^13,26,27^. To test whether binding major capsid protein induces Avs8A nuclease activity, we incubated purified Avs8A and λ gpE with a linear dsDNA substrate and monitored DNA integrity by agarose electrophoresis. Avs8A potently degraded the dsDNA substrate in response to gpE or Bas21 major capsid protein but showed no DNase activity in response to maltose binding protein, which served as a negative control (**Fig. 6 c, d and ED Fig. 3**). Nuclease activity could be visualized over time and required Mg^2+^ and ATP (**ED Fig. 3**). Intriguingly, Avs8A also appeared to require ATP-hydrolysis for nuclease activation because no activity was observed when ATP was substituted for the non-hydrolyzable analog AMP-PNP (**ED Fig. 3**). These results are surprising as the related STAND NTPase Avs4 and NACHT NTPase bNACHT11 do not require ATP hydrolysis for function, but do require ATP for oligomerization, likely through stabilizing a conformational change. It is important to note that we have not determined if Avs8A can bind AMP-PNP similarly to ATP and thus our observations for ATP hydrolysis are not yet conclusive.

These data support a model in which Avs8A senses phage infection through binding major capsid proteins and is activated to destroy phage and host dsDNA. In support of this model, mutation of the active site residues of Avs8A nuclease domain resulted in a complete loss of phage defense (**Fig. 6f**). Recombinant λ gpE is expected to be a monomer because oligomerization requires accessory phage proteins. During infection, 415 monomers irreversibly assemble into the capsid^28^. We tested whether Avs8A was able to sense fully formed phage capsids by incubating Avs8A with phage λ lysate but observed no nuclease activity (**Fig. 6e**).

## Discussion

In this work, we defined how bacteria sense phage infection by searching an expansive library of phage ORFs for proteins that trigger immune signaling. The immune system of bacteria is distributed in the pangenome and therefore our analysis required investigating diverse strains of *E. coli* to survey many phage defense systems. What emerges is an inventory of over 100 phage ORF-ECOR strain pairs that each may represent novel phage stimuli for different phage defense systems. We demonstrate the potential of this dataset by characterizing two of these pairs in detail, which discovered the antiphage system PD-T2-1 and uncovered the activator of Avs8.

Investigating the selective growth inhibition observed for the T7 gp17-ECOR03 pair revealed PD-T2-1, a novel phage defense system that is activated by phage tail fiber proteins T7 gp17, T2 gp37, T2 gp34, and T2 gp12. All of these proteins are structural and required for tail fiber formation in their respective phage, however, the biochemical feature that unites all four of these proteins is only vague. All four proteins form trimeric β-helixes^29–33^. The T2 proteins all appear to have a shared domain found in the T4 proximal long tail fiber protein gp34 (PDB: 5NXF)^34^. In the related phage T4, there is sequence homology shared between gp12 and gp34^34^. It is unclear if T7 gp17 adopts a similar T4 gp34 structure and structure-based comparisons are challenging for these proteins because the three protomers of the trimer interweave to adopt the overall structure. One peculiar feature of the highly similar T4 gp12, T4 gp36, and T4 gp37 is that all three require the phage chaperone gp57A for proper assembly^35^. Other unrelated phage defense systems require host protein chaperones to sense phage infection^36^. These details may provide a clue for the molecular mechanism of PD-T2-1 activation.

Characterization of the gpE-ECOR07 genetic interaction led to identifying that gpE bound to Avs8A at high affinity, which activated DNase activity in the defense system. Avs8A showed an impressive ability to sense major capsid proteins from phages λ, Bas21, and Bas49, which each share <15% sequencing identity. These data suggest that the Avs8A recognizes the conserved HK97 fold protein structure in a way that is buffered against changes in amino acid sequence. This is an ideal feature for an innate immune system, which must maintain the upper hand against a virus that can mutate rapidly to escape detection. Avs8A joins other related P-loop NTPase members of the AVAST system that recognize structures of proteins that are crucial to the viral lifecycle. An interesting feature of all of these proteins is the use of an apparent tandem-repeat like C-terminal domain for ligand recognition. We speculate that biophysical features of these tandem repeats, and perhaps many redundant interactions between receptor and trigger, cage the mutational landscape available to the phage that cannot be escaped without compromising trigger protein function. Another conspicuous feature is that the ligands that active Avs8A and Avs1–4 all must oligomerize for function but only activate their receptors when bound as monomers. We hypothesize that by recognizing ligand interfaces required for oligomerization, the immune receptor further limits the mutational landscape the virus can explore. These defense systems are further complicated because as least one member, KpAvs2, can bind structurally unrelated ligands^37^.

We anticipate the remaining candidate phage trigger-ECOR strain pairs will be valuable dataset for determining the molecular mechanisms for how additional antiphage systems are activated. In addition, our analysis provides a list of phage genes that universally inhibit *E. coli* growth. While some of those ORFs may be expressed by phages to initiate lysis and complete the viral lifecycle, we expect others may be important to reprogram and remodel central processes within the cell. The growth inhibition phenotypes observed here are a starting point for characterizing the function of these proteins and better understanding the viral lifecycle.

Our approach is conceptually similar to Silas et al.^38^, who used a similar plasmid-based screen of phage accessory genes that revealed both inhibitors of phage defense systems and phage inhibitors that themselves activate defense systems. Together with many other studies, we find that phage defense systems often fall into two broad categories that reflect conserved strategies of innate immune signaling^11^. Defense systems either bind phage components directly or indirectly sense phage activities through an intermediate. Avs8A is an example of direct sensing whereas PD-T2-1 may be an example of indirect sensing. By continuing to investigate and understand molecular mechanisms of how antiphage systems are activated, we hope to expand and elaborate these paradigms.

## Supporting information

Alphafold of PD-T2-1

Alphafold of Phage PAMPs

Alphafold of Avs8 and nonHK97 capsids

Alphafold of Avs8

Alphafold of Avs8 and HK97 capsids

## Acknowledgements

The authors would like to thank L. Aravind for discussions and insights into PD-T2-1 and Avs8; Annette Erbse and the Shared Instruments Pool (RRID: SCR_018986) of the Department of Biochemistry at the University of Colorado Boulder for providing access to the Avanti JXN 26 Super Speed centrifuges and rotors, which are funded by NIH Grant R24OD033699-01, as well as the ITC 200, funded by the NIH Shared Instrumentation Grant S10RR026516; Aedan Monahan for preparation of linear DNA substrate for nuclease assays and mini-preps of vectors; and all members of the Whiteley lab for their advice and helpful discussion. This work was funded by the National Institutes of Health through the NIH Director’s New Innovator award DP2AT012346 (A.T.W.); the PEW Charitable Trust Biomedical Scholars Award (A.T.W.); the Boettcher Foundation Webb-Waring Biomedical Research Award (A.T.W.); and the Burroughs Wellcome Fund PATH Award (A.T.W.).

## Author contributions

Conceptualization, T.A.N. and A.T.W.; Methodology, T.A.N. and A.T.W; Investigation, T.A.N., G.W.G, and A.T.W.; Resources, A.N.C.; Writing – Original Draft, T.A.N. and A.T.W.; Writing – Review & Editing, T.A.N., A.N.C., and A.T.W; Visualization, T.A.N.; Supervision, A.T.W.; Funding Acquisition, A.T.W.

## Declaration of interests

The authors declare no competing interests.

## Methods

### Bacterial strains and culture conditions

*E. coli* strains used in this study are listed in **Table S8A**. Bacteria were grown in LB medium (1% tryptone, 0.5% yeast extract, and 0.5% NaCl) at 37 °C with shaking at 220 rpm in 2 mL of media in 14 mL culture tubes, unless otherwise indicated. Carbenicillin (100 μg/mL) or chloramphenicol (20 μg/mL) were added for plasmid selection when appropriate. All strains were frozen for storage in LB plus 30% glycerol at ™70 °C. *E. coli* OmniPir was used for construction and propagation of all plasmids. *E. coli* MG1655 (CGSC6300) was used to collect all experimental data. *E. coli* BL21 (DE3) (NEB cat#: C2527H) was used to express proteins for purification.

For phage and colony formation assays, bacteria were grown in MMCG medium (47.8 mM Na_2_HPO_4_, 22 mM KH_2_PO_4_, 18.7 mM NH_4_Cl, 8.6 mM NaCl, 22.2 Mm Glucose, 2 mM MgSO_4_, 100 μM CaCl_2_, 3 μM Thiamine, Trace Metals at 0.1× (Trace Metals Mixture T1001, Teknova, final concentration: 8.3 μM Ferric chloride, 2.7 μM Calcium chloride, 1.4 μM Manganese chloride, 1.8 μM Zinc Sulfate, 370 nM Cobalt chloride, 250 nM Cupric chloride, 350 nM Nickel chloride, 240 nM Sodium molybdate, 200 nM Sodium selenite, 200 nM Boric acid)) with appropriate antibiotics.

When growing strains that required induction, 50 μM IPTG or 0.2% arabinose was used to induce, as appropriate.

### Identification of antiphage systems in ECOR strains

The Prokaryotic Antiviral Defense Locator (PADLOC)^8^ webserver was used to catalog previously-identified defense genes in sequenced ECOR strains appearing in **Table S3**^39^. Results were obtained as precomputed data generated running PADLOC v2.0.0 with PADLOC DB v2.0.0 over the ECOR RefSeq draft genome sequences.

### Phage amplification and storage

The phages used in this study are listed in **Table S8B**. Phage were liquid amplified by infecting a 5 mL mid-log cultures of *E. coli* MG1655 in LB plus 10 mM MgCl_2_, 10 mM CaCl_2_, and 100 μM MnCl_2_ were infected with phage at an MOI of 0.1 and grown, shaking, for 2–16 hours. The supernatant was harvested and filtered through a 0.2 μm spin filter to remove bacterial contamination.

### Plasmid construction

The plasmids used in this study are listed in **Table S8A**. Genes of interest were amplified from using Q5 Hot Start High Fidelity Master Mix (NEB, M0494L). Phage genomic DNA, or reverse-transcribed cDNA for MS2, served as the template for phage genes and defense system coding sequences and endogenous regulatory regions were amplified from the genomic DNA of *E. coli* strains from the ECOR collection^14^ Avs8 point mutations were generated by amplifying out the gene of interest in two parts from a plasmid template, with the desired mutation occurring in the overlapping region between the two amplicons. PCR products were ligated into restriction-digested, linearized vectors by modified Gibson Assembly^40^. Gibson reactions were transformed via electroporation into competent OmniPir and plated onto appropriate antibiotic selection. Unless otherwise indicated, all enzymes were purchased from New England Biolabs.

For all vectors using the pLOCO2 backbone, pAW1382 was amplified and purified from OmniPir. Purified plasmid was then linearized using SbfI-HF and NotI-HF or FseI-HF. For all vectors using the pTACxc backbone, pAW1608 was amplified and purified from OmniPir. Purified plasmid was then linearized using BamHI-HF and NotI-HF. For all vectors using the pBAD30x backbone, pAW1367 was amplified and purified from OmniPir. Purified plasmid was then linearized using EcoRI-HF and HindIII-HF. For all vectors using the pETSUMO2 backbone, pAW1123 was amplified and purified from OmniPir. Purified plasmid was then linearized using BmtI-HF and NotI-HF. Sanger sequencing (Genewiz or Quintara) was used to validate the correct sequence within the multiple cloning site.

### Orfeome generation and screening

Geneious Prime was used to extract a total of 414 ORFs predicted to encode proteins across T2, T7, λ*vir*, <X174, M13, and MS2 genomes. Primers were automatically designed for each ORF to individually PCR amplify sequences, and each amplicon was cloned into linearized pAW1608, and transformed into OmniPir and selected on antibiotic media. Three colonies from each transformation were grown in LB + antibiotic for 18hours and clones were pooled, plasmids extracted, and DNA sequenced using 200Mbp Illumina Sequencing (SeqCenter). For both types of sequencing, reads were mapped to the predicted plasmid sequence using the *Map to Reference* feature of Geneious Prime (default settings).

Plasmids containing phage ORFs were pooled and 180ng was transformed into *E. coli* MG1655 or ECOR strains 1-72, in duplicate. After one hour of recovery in S.O.C. at 37 °C and 220 rpm, cells were plated onto MMCG agar containing 50 μM IPTG and 20 μg/mL chloramphenicol, and MMCG agar containing 20 μg/mL chloramphenicol for *E. coli* MG1655. After 18 hours of growth at 37 °C, surviving colonies with isolated by miniprep. 20 μL of miniprep was sequenced using 200Mbp Illumina Sequencing (SeqCenter). Reads were mapped to phage reference sequences using the *Map to Reference* feature of Geneious Prime (default settings). We employed the Transcripts Per million (TPM) feature in Geneious to compare relative read abundance between ORFs in different ECOR strains. Average TPM across replicate samples was used to compare ORF abundance between uninduced MG1655 and MG1655 or ECOR strains 1-72 under IPTG induction. ORFs that had less than 10 reads under non-inducing conditions in MG1655 were considered to be basally growth inhibitory, and those with 10-fold decrease in reads under IPTG induction were considered as growth inhibitory under induction. ORFs that inhibited growth of MG1655 were not included in our analysis of potential phage triggers.

### Transformation efficiency assay

Recipient strains were made electrocompetent and 40 ng of control or phage ORF-containing plasmid were electroporated in 50 μL of competent cells. Cells were then incubated in 1 mL of S.O.C. for 1 hour at 37 °C with shaking at 220 rpm, then serially-diluted and plated on LB agar supplemented with 20 μg/mL chloramphenicol, with or without 50 μM IPTG. Transformation efficiency was assessed by counting colony forming units after overnight incubation at 37 °C.

### Transposon mutagenesis and screening

*E. coli* MFD^41^ containing pSC189^42^, which contains a himar-based transposon, was grown overnight in LB + 300 μM DAP (2,6-Diaminopimelic acid) and 100 μg/mL carbenicillin. Simultaneously, ECOR03 or ECOR07 were grown in LB overnight. The following day, 1mL of each strain was pelleted via centrifugation, washed in 1mL of LB plus 300 μM DAP, and pelleted again. Pellets of MFD + pCS189 and ECOR03 or ECOR07 were combined by resuspending one strain and then the other in the same 50 μL of LB+ 300 μM DAP. The combined strains were spotted onto a mating filter (Pall Corporation), which was placed on and LB plus 300 μM DAP agar plate. Spots were allowed to dry under a flame for approximately 10 minutes prior to incubation at 37 °C for 1 hour. After mating, the filter was placed into a 15 mL conical tube contain 2mL of LB and vortexted for 1 minute to remove bacteria from filters. Conjugates containing transposon insertions were selected for by plating 300 μL onto 3x 15cm^2^ LB agar plates containing 25 μg/mL kanamycin. 300 μL of a 10^−1^ dilution were plated onto 3x 15cm^2^ LB agar plates containing 25 μg/mL kanamycin to ensure single colonies.

After incubation overnight at 37 °C, all mutants were collected using a cell scraper, washed 2 times with ice cold water, and resuspended in 1 mL of 10% glycerol, creating a transposon library. 100 μL of electrocompetent ECOR03 was transformed with 100 ng of T7 gp17 and 100 μL of electrocompetent ECOR07 was transformed with 100 ng of λ p08. Cells were then incubated in 1 mL of S.O.C. for 1 hour at 37 °C with shaking at 220 rpm, then 300 μL plated on 3×15cm^2^ MMCG agar plates supplemented with 50 μg/mL kanamycin and 20 μg/mL chloramphenicol, with 50 μM IPTG. Plates were grown overnight at 37°C.

Individual mutants were gridded into a 96-well plate containing 200 μL of LB with 50 μg/mL kanamycin and 20 μg/mL chloramphenicol, grown overnight at 37°C with shaking at 220 rpm, and 100 μL of 50% glycerol was added to each well prior to storage at -70 °C.

### Arbitrarily primed PCR transposon mapping

Transposon mutants in ECOR strains that survived induction of the phage trigger were mapped using arbitrary PCR as previously described^42,43^. A scraping of frozen transposon mutants was used as a template in the first round of PCR using primers: pSC189-PCR1 GGCTGACCGCTTCCTCGTGCTTTAC and Arb1 GGCCACGCGTCGACTAGTACNNNNNNNNNNCTTCT. 1 μL of the product of this reaction was used as a template for the second round of PCR using pSC189-PCR2 AATGATACGGCGACCACCGAGATCTACACTCTTTCGGGGACTTATCAGCCAACCTG and Arb2 GGCCACGCGTCGACTAGTAC. The reaction was purified using Exo-CIP (NEB) treatment as recommended by the manufactures. PCR products were Sanger Sequenced (Azenta and Quintara) using CTTTCGGGGACTTATCAGCCAACCTGTTA.

The resulting sequences were then aligned to the genome of ECOR03 or ECOR07 to identify the TA site at which the transposon was inserted (**Table S5** and **S6**). Blast and HHpred^44^ (Geneious prime software 2023) were used to determine gene identity, predicted domains, and predicted function. The Identical Protein Group data, found in NCBI, was used as a proxy to determine abundance and taxonomic distribution for PD-T2-1 and Avs8 genes.

### Phage resistance analysis

A modified double agar overlay was used to measure the efficiency of plating (EOP) of phages^45,46^. Overnight cultures, defined as bacterial cultures grown 16– 20 hours post-inoculation from a glycerol stock, of *E. coli* MG1655 expressing the indicated plasmids cultured in MMCG plus appropriate antibiotics were diluted 1:10 into the same media and cultivated for an additional two hours to reach OD_600_ 0.1–0.4. 400 μL of the mid-log culture was mixed with 3.5 mL MMCG (0.35% agar) with an additional 5 mM MgCl_2_ and 100 μM MnCl_2_. After three inversions, the mixture was poured onto an MMCG (1.6% agar) plate and allowed to cool. 2 μL of a phage dilution series in SM buffer was spotted onto the overlay, dried for 10 minutes under a flame, plates incubated overnight at 37 °C, and plaque formation enumerated the following day. When no plaque formation or hazy zone of clearance occurred, 0.9 plaques at the least dilute spot were used as the limit of detection.

### Colony formation assays

*E. coli* expressing indicated plasmids were cultivated overnight in MMCG with appropriate antibiotics. Cultures were diluted in a 10-fold series into MMCG and 5 μL of each dilution was spotted onto MMCG agar plates containing appropriate antibiotics with or without 0.2% arabinose. Spots of bacteria were dried for 10 minutes under a flame, incubated overing at 37°C and growth was enumerated the next day by counting colony forming units (CFU/mL) of each strain. In instances when no individual colonies could be counted, the lowest bacterial concentration at which growth was observed was counted as 10 CFU. If no colonies could be counted at the highest bacterial concentration, 0.9 CFU was used as the limit of detection.

### Protein purification

Avs8A, λ capsid (λ gpE), Basel 21 capsid, T7 gp17, and MBP were expressed as 6x-His-SUMO fusion proteins in *E. coli* BL21 (DE3) cells. Plasmids were transformed into BL21 (DE3) using electroporation and plated on LB-agar supplemented with carbenicillin 100 µg/mL plus 1% glucose and grown overnight at 37°C. A single colony was then used to inoculate a 50 ml starter culture of LB supplemented with 100 µg/ml carbenicillin and 1% glucose, which was grown overnight at 37°C. 10 mL of the overnight culture was used to inoculate 500 mL-1 L of LB media in 2.5 L Thompson flasks (500 mL / flask). Cultures were grown at 37°C with shaking at 220 rpm until reaching an OD_600_ of 0.600-0.800, and 500µM IPTG was added prior to an additional 18 hours of growth at 20°C. Cells were harvested by centrifugation at 4000 x *g* and resuspended in Buffer 1: 20 mM HEPES pH 7.5, 400 mM NaCl, 30 mM imidazole, 10% glycerol, and 1 mM DTT. Cells were sonicated on ice at an amplitude of 70, for 30 seconds on / off for 20 minutes total. Sonicated lysates were centrifuged at 14,000 x *g* for 1 hour at 4°C to pellet debris, and the supernatant was run over 1 ml of Ni-NTA resin (Thermo Fisher Scientific) equilibrated with Buffer 1. The resin was washed with 50 mL of Buffer 2: 20 mM HEPES pH 7.5, 1M NaCl, 30 mM imidazole, 10% glycerol, and 1 mM DTT to remove nonspecific components bound to the resin. Proteins were eluted using Buffer 3: 20 mM HEPES pH 7.5, 400 mM NaCl, 300 mM imidazole, 10% glycerol, and 1 mM DTT. For all constructs except Avs8A, the 6xhis-SUMO tag was cleaved using 6xhis-ULP1 (produced in-house) via dialysis overnight at 4°C against 2L of Buffer 4: 20 mM HEPES pH 7.5, 250 mM KCl, and 1 mM DTT using a 10 kDa MWCO dialysis membrane. The cleaved sample was run over 2 ml Ni-NTA to capture cleaved 6xhis-SUMO tag and uncleaved protein and the flowthrough containing the recombinant tag-free protein was collected. All proteins were concentrated via 3 kDa, or 100kDa for Avs8A, ultracentrifugation filter at 4000 x *g* at 4°C and stored at -80°C until needed. Protein purity was assessed via SDS-PAGE followed by Colloidal Coomassie staining.

### Microscale thermophoresis

Protein binding affinities were measured using microscale thermophoresis^47^. Avs8-A fused to 6xHis-SUMO was labeled with Monolith His-Tag Labeling Kit RED-tris-NTA 2nd Generation (Nanotemper: Cat# MO-L018) following manufacturer instructions. Samples were equilibrated for 30 minutes at room temperature before MST measurement. Experiments were performed using independently labeled proteins and ligand titrations in MST Buffer (20 mM HEPES pH 7.5, 250 mM KCl, 5mM MgCl_2_, 0.05% v/v Tween-20, 0.1 mM DTT). Measurements were performed using 60%-80% laser excitation, medium MST power, and a chamber temperature of 25 °C on a Nano-BLUE/RED Monolith NT.115 (NanoTemper). All data was analyzed using a hot time of 9–10 seconds. Fraction bound values were calculated by Mo.AffinityAnalysis software (NanoTemper). Binding data from three independent experiments were fit using the quadratic binding equation^48^.

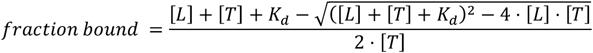

In this equation [L] is the concentration of ligand, [T] is the concentration of labeled target (50 nM for all experiments reported here). The dissociation constants reported here represent the average of the dissociation constants calculated for each biological replicate ± standard error of the mean.

### Nuclease assays

Purified Avs8A (100 nM) was incubated with λ gpE, Basel 21 mcp, or MBP (1μM) and 100ng nucleic acid substrate in 20 μL reactions containing 50 mM HEPES pH 8.0, 15 mM NaCl, 1 mM DTT. 4 mM ATP or AMP-PNP and 10 mMMgCl_2_ was added where indicated^49,50^. Reactions were incubated at room temperature for the indicated time points, 5 μL of 6x DNA loading dye was added (final concentration 3.3 mM Tris-HCl pH 8, 2.5% Ficoll-400, 10 mM EDTA, 0.05% Orange G) to stop reactions, and run for 30 minutes at 160V on a 1% agarose gel containing ethidium bromide (1% agarose, 40 mM Tris, 20 mM acetic acid, 1 mM EDTA, SYBR Safe DNA stain). Gels were imaged using an Azure Biosystems Azure 200 Bioanalytical Imaging System.

### Accession numbers of molecules appearing in this study

T2 phage genome: NC_054931

T7 phage genome: NC_001604

λ*vir* phage genome: NC_001416

M13 phage genome: JX412914

ϕX174 phage genome: NC_001422

MS2 phage genome: NC_001417

ECOR03 PD-T2-1A: RCR47252.1; WP_075861856.1

ECOR03 PD-T2-1B: RCR47253.1; WP_08789477.1

ECOR07 Avs8A: WP_000963924.1

ECOR07 Avs8B: WP_000443183.1

T7 gp17, locus tag: T7 p52; protein accession: NP_042005.1

T7 gp1.7, locus tag: T7 p14; protein accession: NP_041967.1

T7 gp13, locus tag: T7 p48; protein accession: NP_042001.1

T2 gp37*, locus tag: KMC23_gp164, protein accession: YP_010073897.1

T2 gp34*, locus tag: KMC23_gp167: protein accession: YP_010073894.1

T2 gp12*, locus tag: KMC23_gp252: protein accession: YP_010073809.1

λ*vir* gpE, locus tag: λ p08; protein accession: NP_040587.1

λ*vir* gpK, locus tag: λ p19; protein accession: NP_040598.1

λ*vir* gam, locus tag: λ p42; protein accession: NP_040618.1

λ*vir* gpN, locus tag: λ p49; protein accession: NP_040625.1

λ*vir* cI, locus tag: λ p88; protein accession: NP_040628.1

Basel 21 major capsid protein: QXV79517.1

Basel 22 major capsid protein: QXV82347.1

Basel 23 major capsid protein: QXV85745.1

Basel 24 major capsid protein: QXV84465.1

Basel 25 major capsid protein: QXV85427.1

Basel 49 major capsid protein: QXV77910.1

*Phage T4 gene names were used to ease comparisons to the literature. T2 and T4 loci are highly similar, however, there are multiple genomes available for T2 phage that differ in annotations.

### Statistics and reproducibility

Each experiment presented was performed at two to three independent times using cultures grown on separate days. Data was plotted using Graphpad Prism 9. Figures were created using Adobe Illustrator CC 2024 v28.6.0.

## Data availability

All data supporting the findings of this study are available within the paper and its Supplementary Information. Supplementary File 1 containing all Alphafold predictions has been split into unique folders for phage PAMPs, PD-T2-1, Avs8, Avs8 co-folded with HK97 capsids, and Avs8 co-folded with non-HK97 capsids.

## Extended Data Figures

**Extended Data Figure 1.**
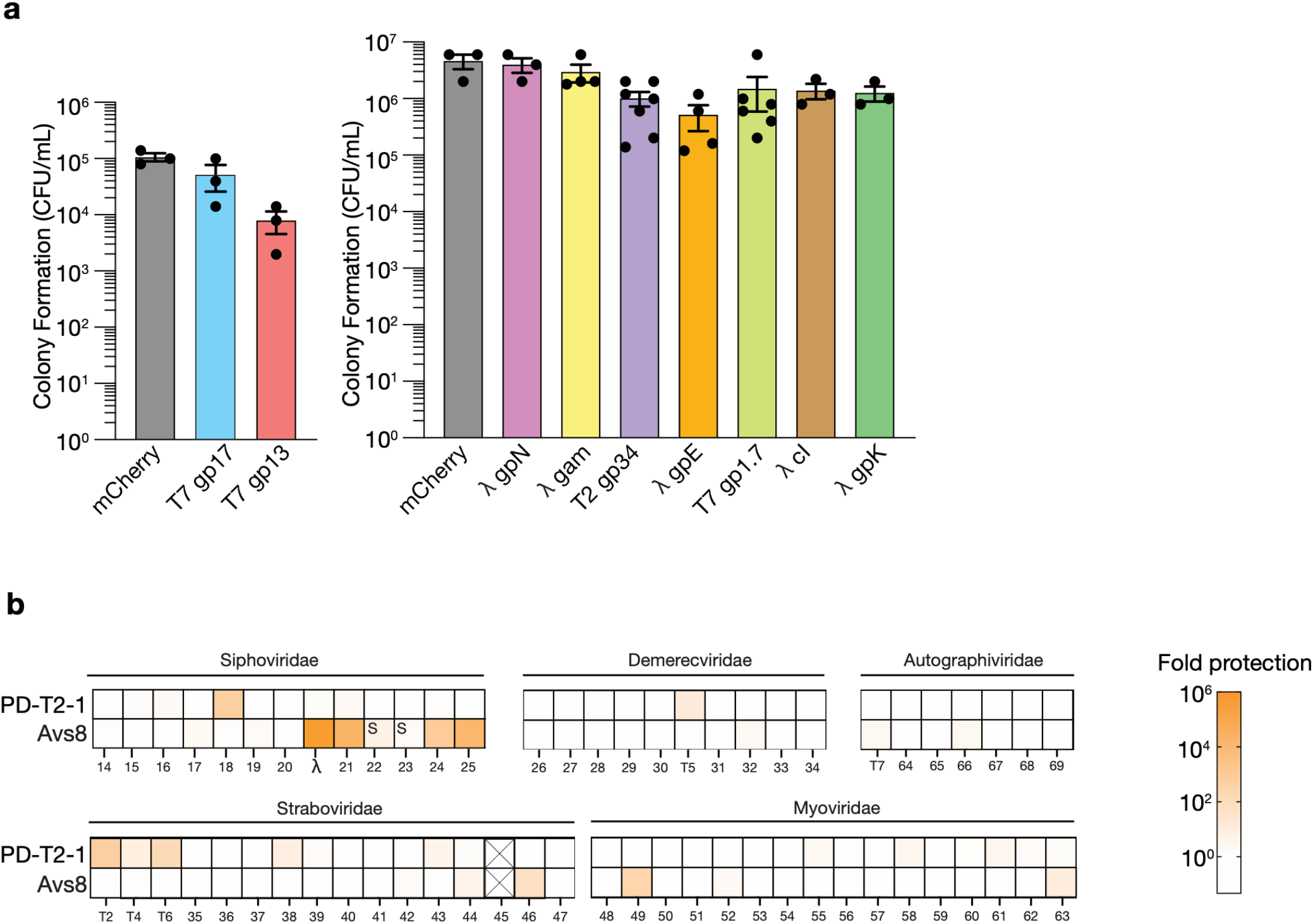
PD-T2-1 and Avs8 protect against diverse phages. **(a)** Number of colonies of MG1655 recovered when transformed with a plasmid expressing mCherry or the indicated phage ORF and grown in medium containing 50 µM IPTG. Data are the mean ± standard error of the mean (SEM) of n≥3 biological replicates. **(b)** Plaque formation (PFU/mL) was enumerated for each phage infecting MG1655 expressing either GFP, PD-T2-1 or Avs8. Fold protection was calculated by dividing the efficiency of plating on GFP by the efficiency of plating on defense system-expressing bacteria and depicted as a heatmap on a log scale. An “X” represents uncountable plaques in any condition. The number on the x-axis refers to the Basel phage used. Phage are grouped taxonomically by family^51^.

**Extended Data Figure 2.**
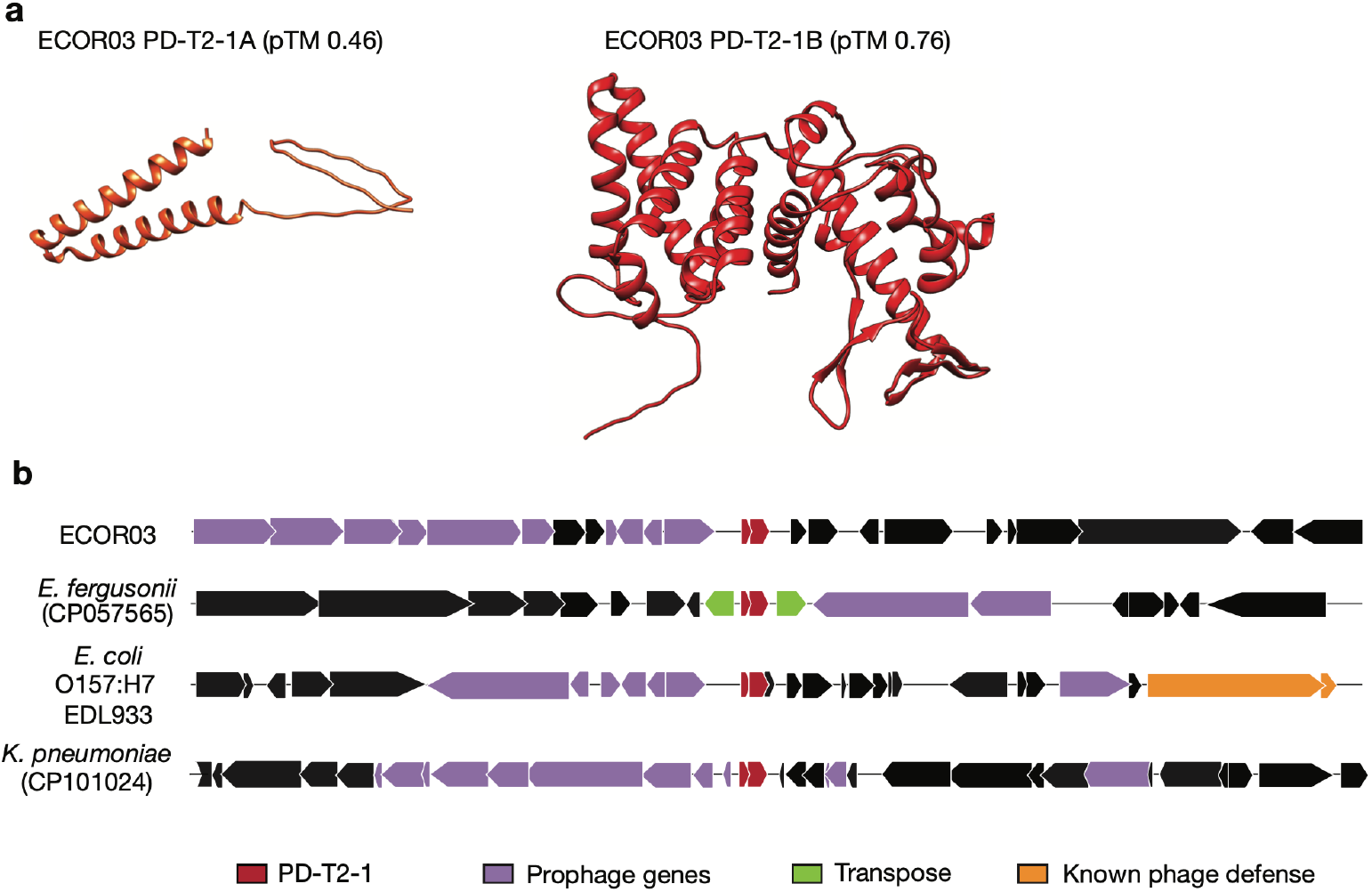
PD-T2-1. **(a)** AlphaFold3 predicted structures of PD-T2-1A and PD-T2-1B. See **Supplementary File 1** for models and statistics. (**b**) Gene neighborhoods ± 10 kb of PD-T2-1 in the indicated Gammaproteobacteria.

**Extended Data Figure 3.**
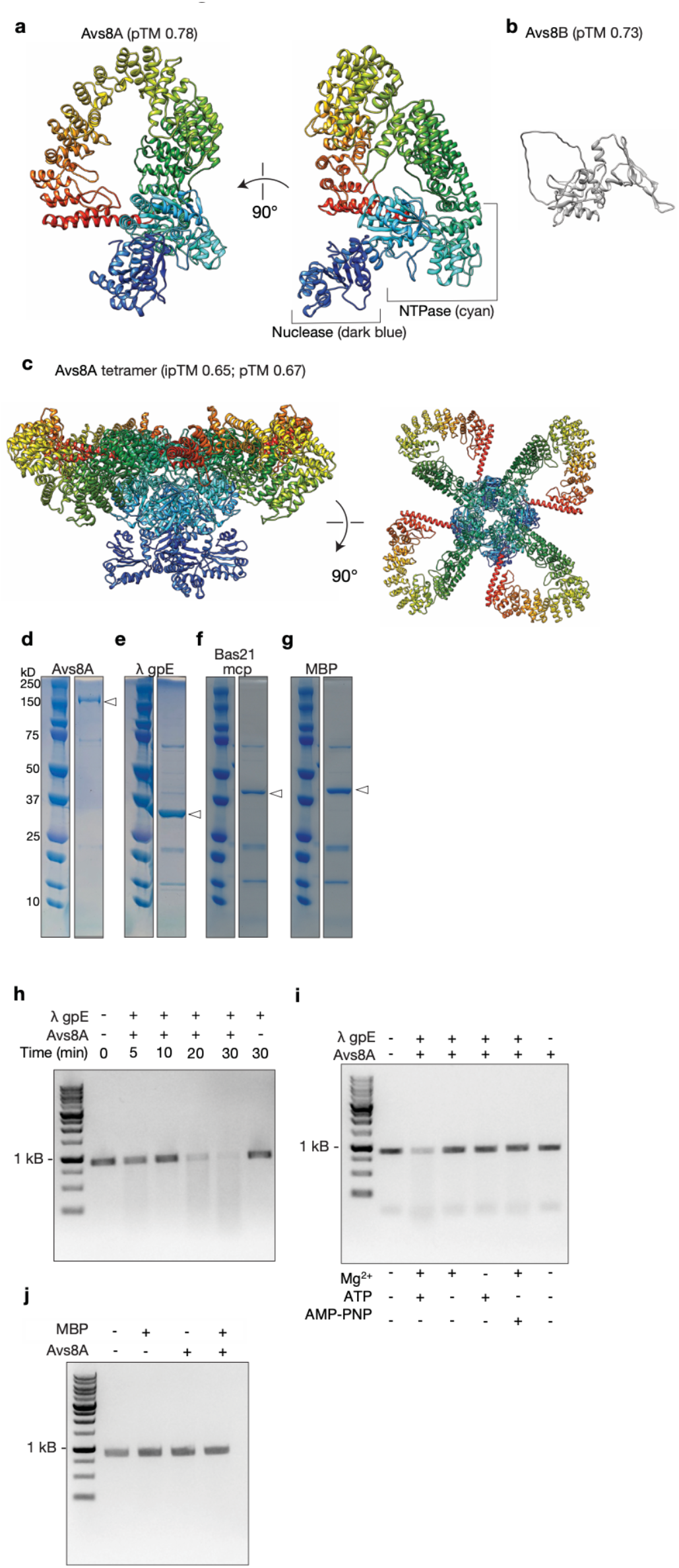
Avs8-A. (**a**) AlphaFold3 predicted structures of Avs8-A and (**b**) Avs8-B. (**c**) Predicted tetramer formation by Avs8-A. (**d–g**) Coomassie-stained SDS-PAGE gels of the indicated purified protein, an arrow shows the expected recombinant protein. Expected molecular weights: 6x-His-hSUMO-Avs8-A: 155.8 kDa, λ gpE (expected MW: 38.2 kDa), Bas21 major capsid protein (mcp, expected MW: 38.5 kDa), and maltose binding protein (MBP, expected MW: 42 kDa). (**h–i**) Agarose gel electrophoresis of linear dsDNA substrate incubated with the indicated proteins or in the indicated conditions. Data are representative of n=3 biological replicates. For a–c, see **Supplementary File 1** for models and statistics.

**Extended Data Figure 4.**
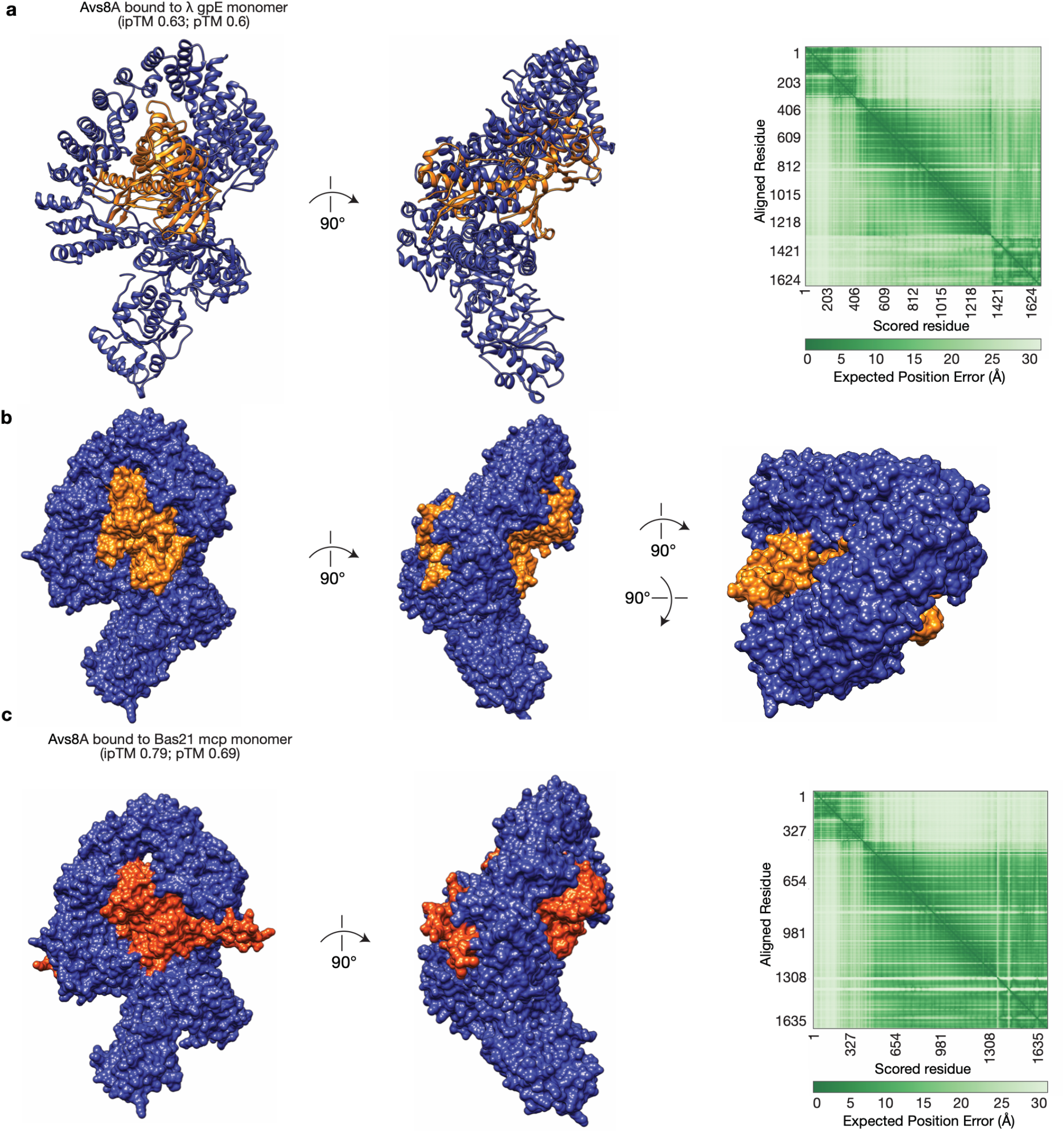
Structure predictions of Avs8-A interacting with major capsid proteins. AlphaFold3 predicted structures of (**a**,**b**) Avs8A and λ gpE and Avs8A and Bas21 mcp. See **Supplementary File 1** for models and statistics.

**Extended Data Figure 5.**
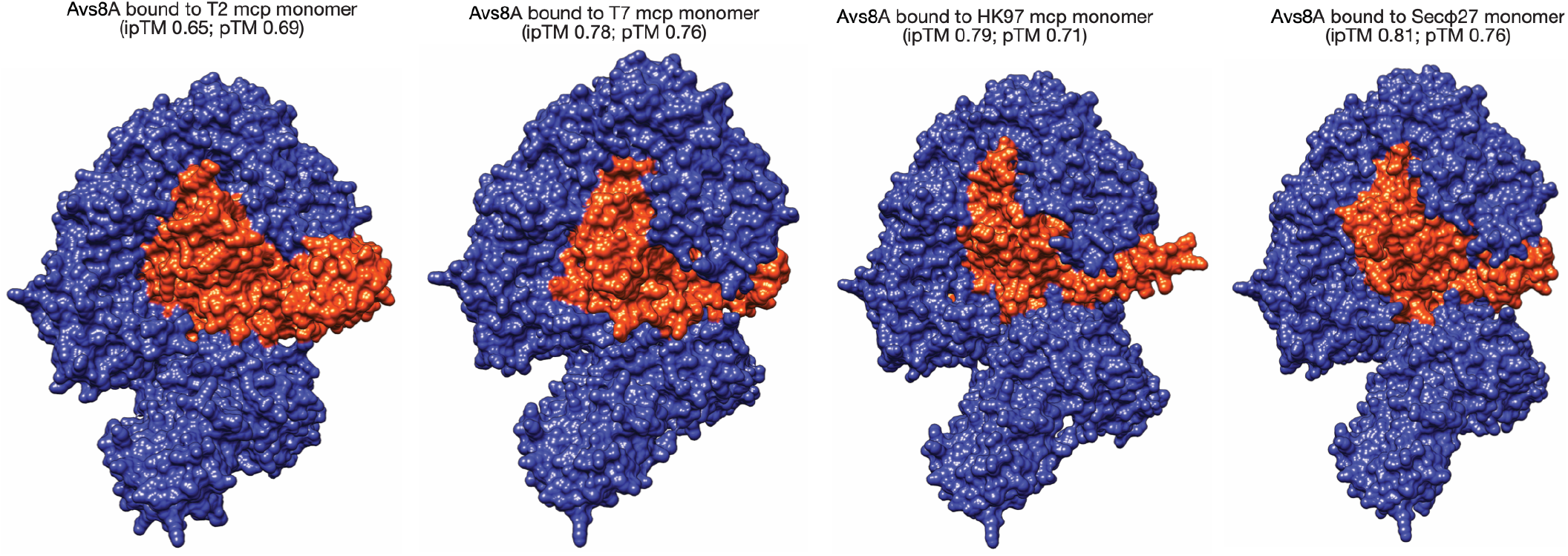
Structure predictions of Avs8A interacting with HK97 fold major capsid proteins. AlphaFold3 predicted structures of Avs8A with the indicated major capsid protein (mcp). See **Supplementary File 1** for models and statistics.

## Notes

### Competing Interest Statement

The authors have declared no competing interest.

