## Supplementary figures and images for "Identifying phage proteins that activate the bacterial innate immune system"

### blueorange90degrees.png

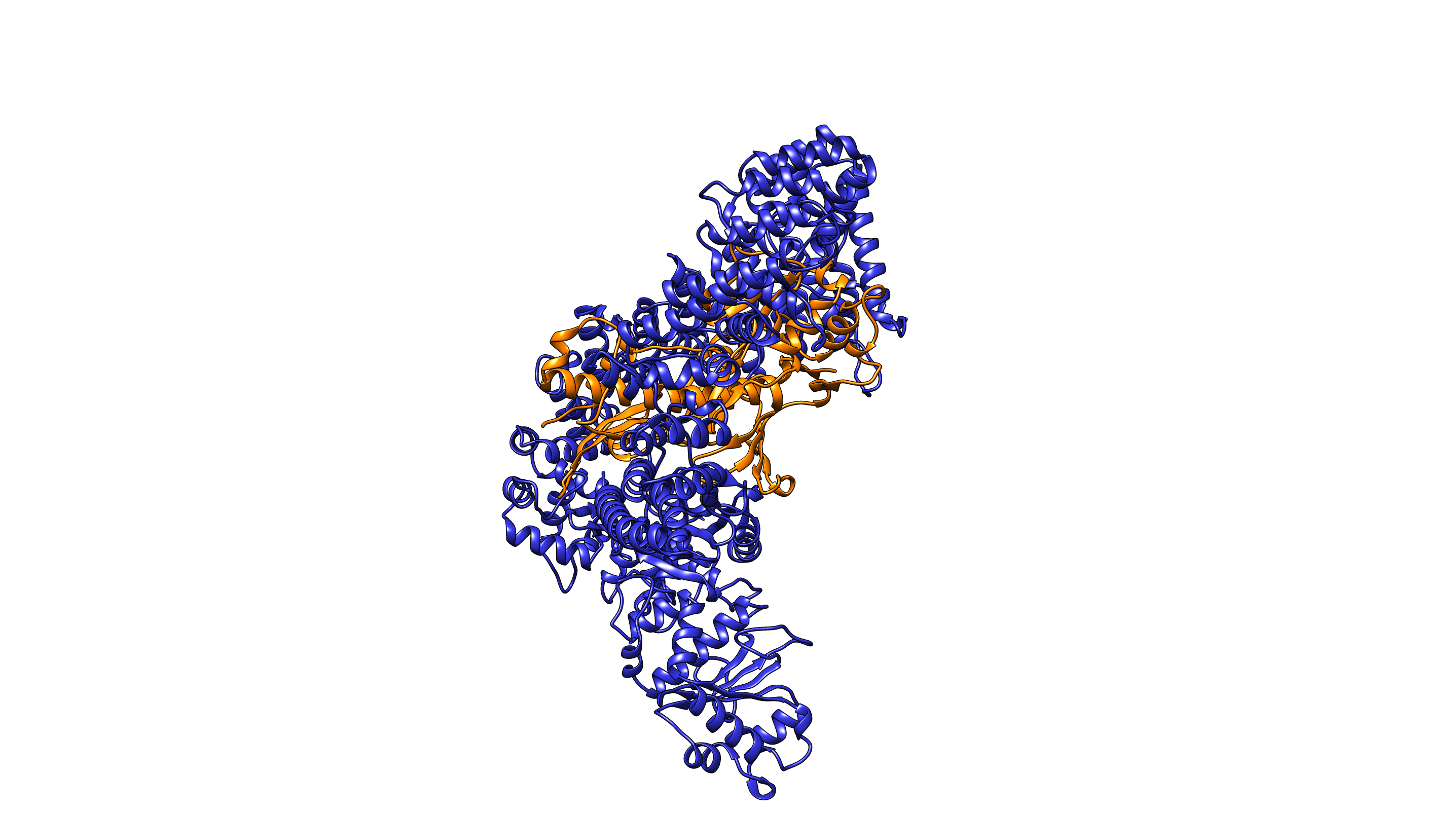

### blueorange.png

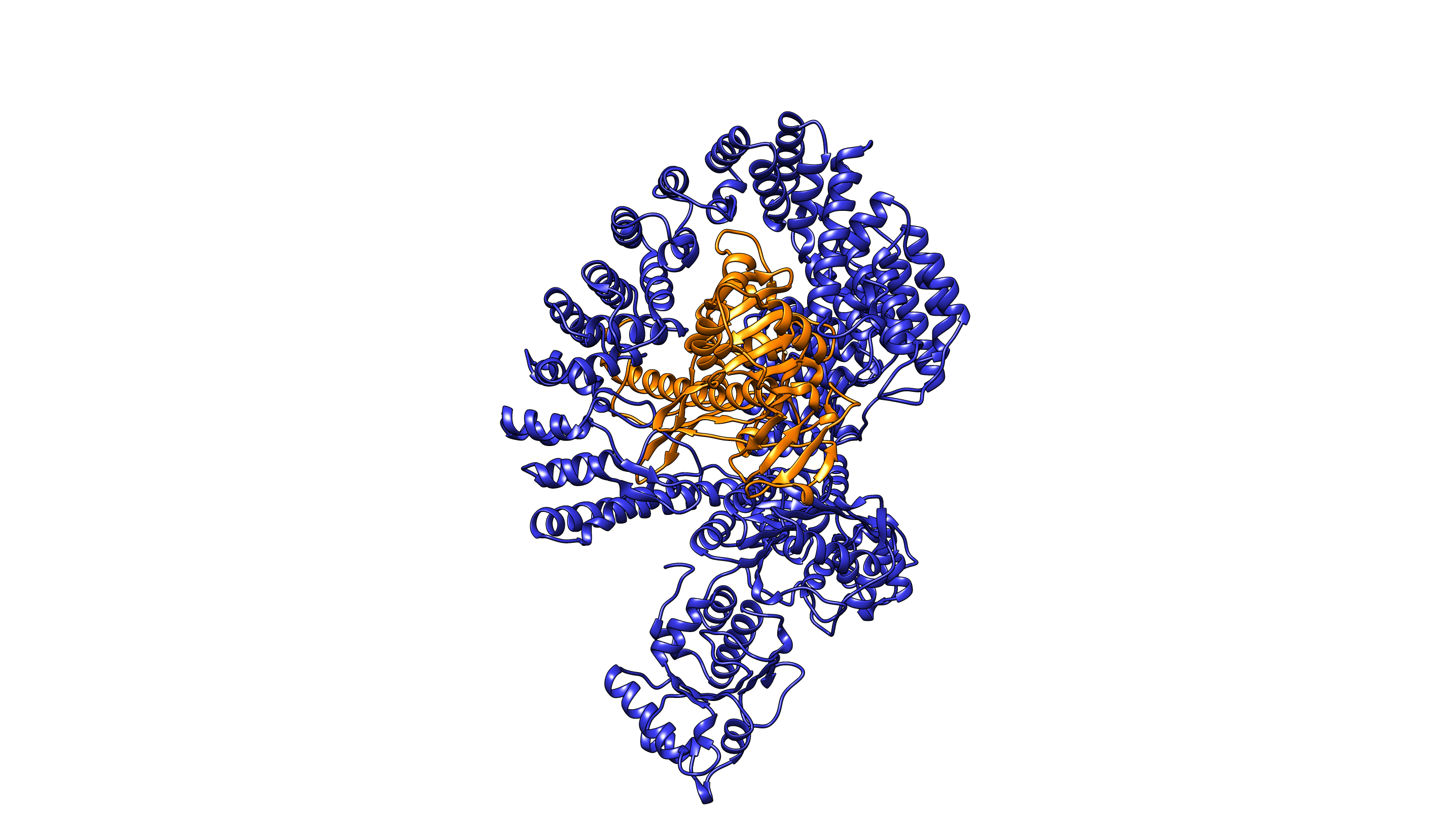

### blueorangesolid90degrees.png

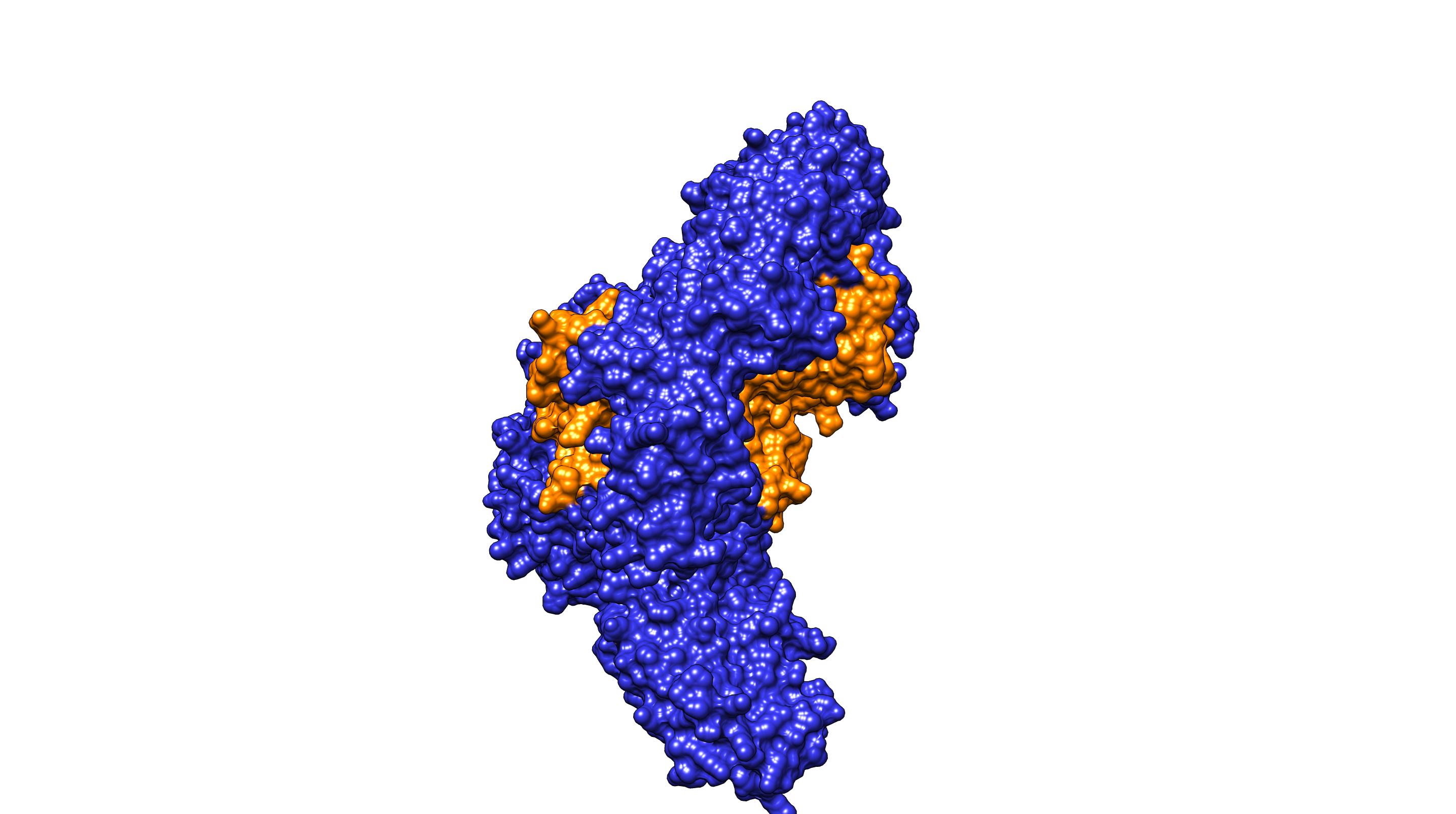

### blueorangesolid90degreestimes2.png

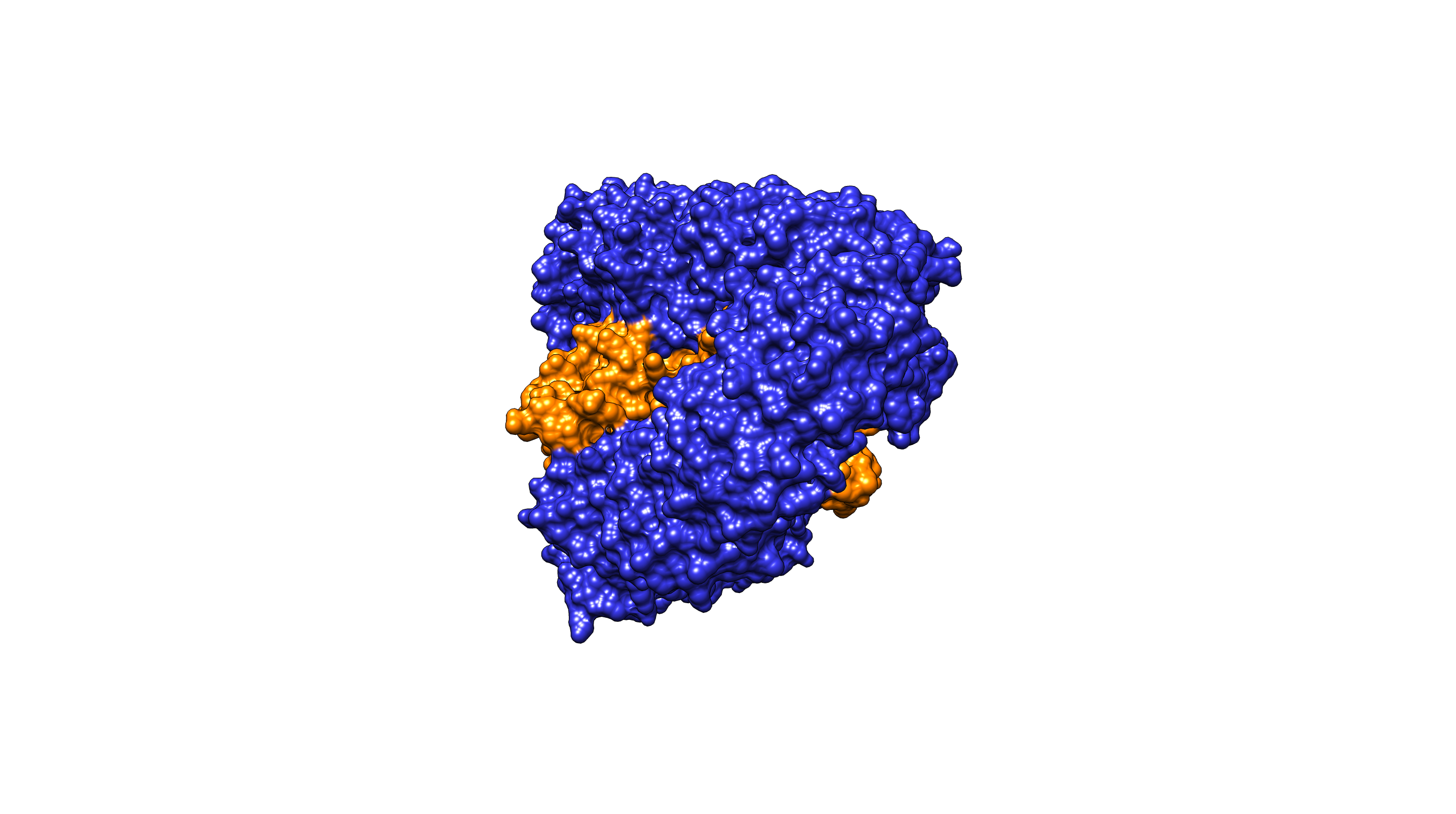

### blueorangesolid.png

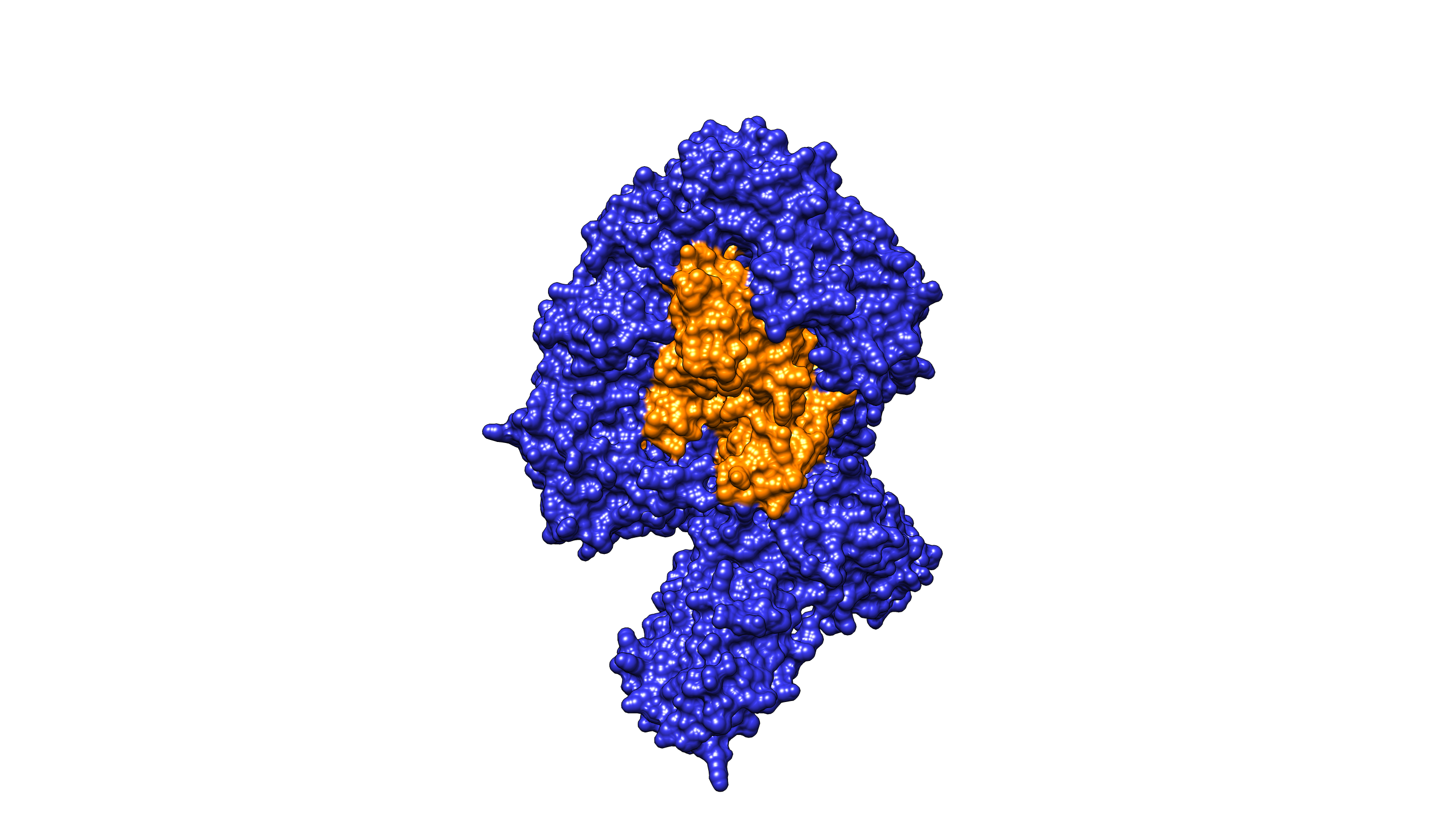

### dark blue.png

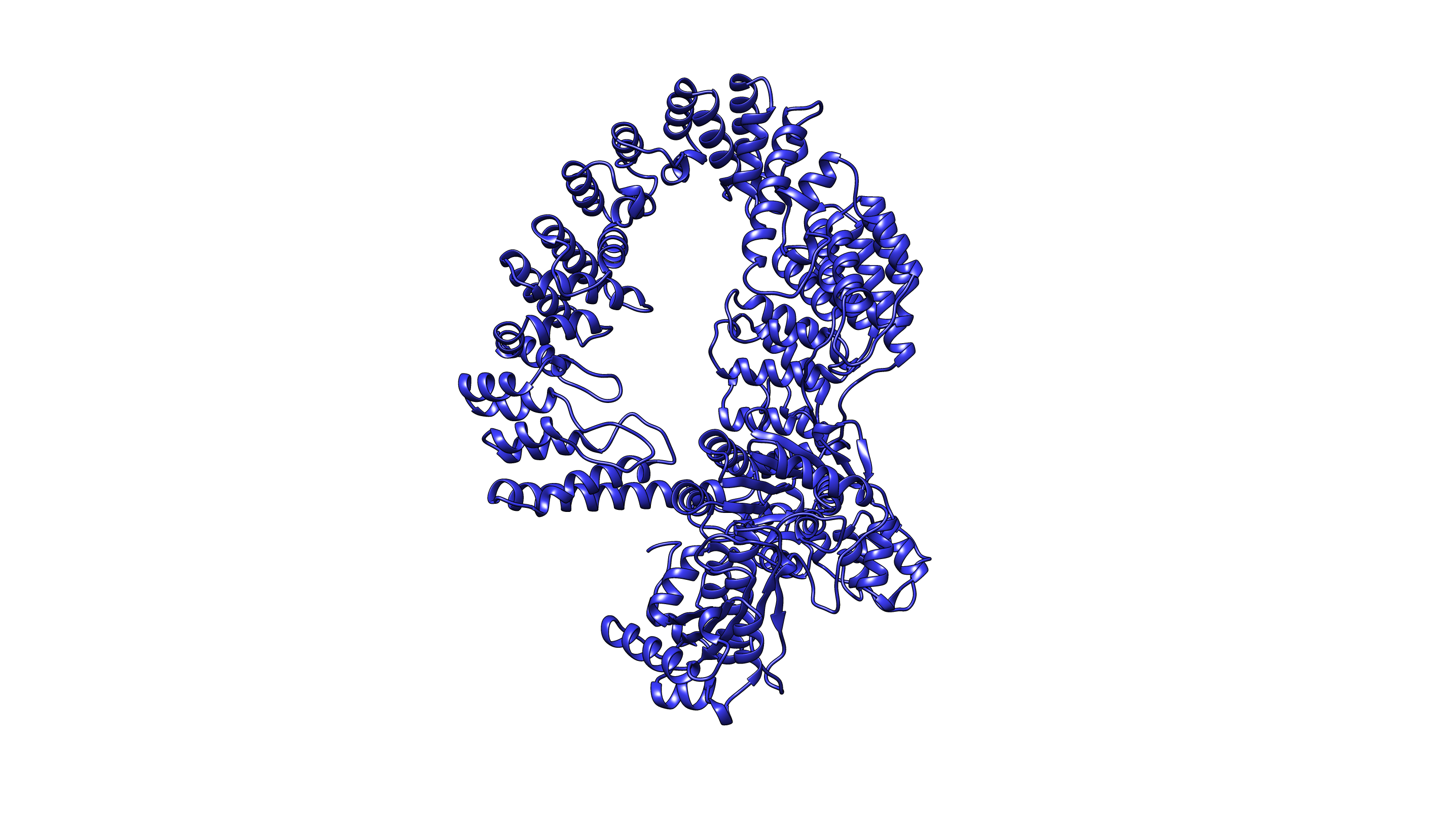

### darkblue orange90degrees.png

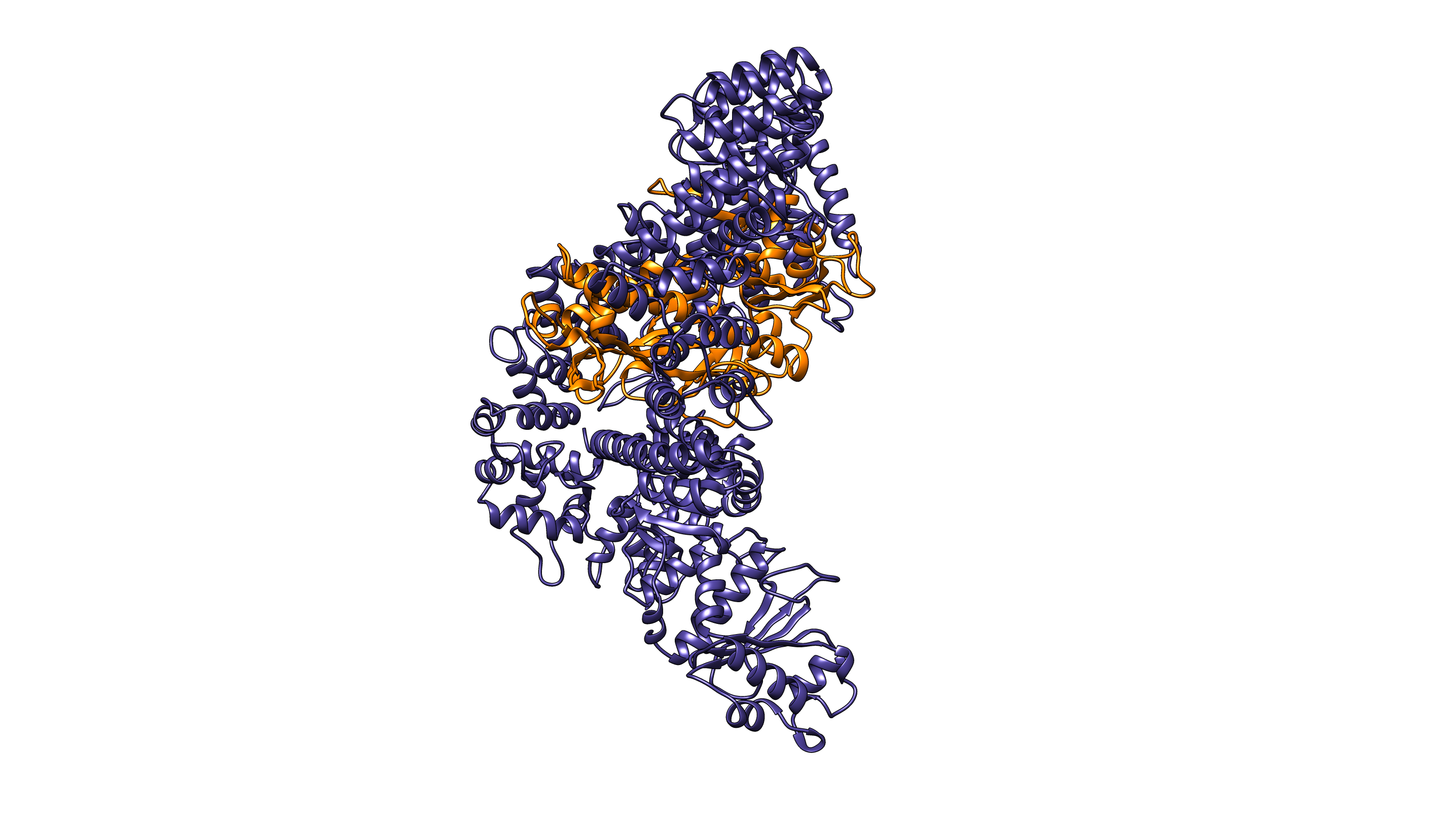

### darkblue orange 90 degrees.png

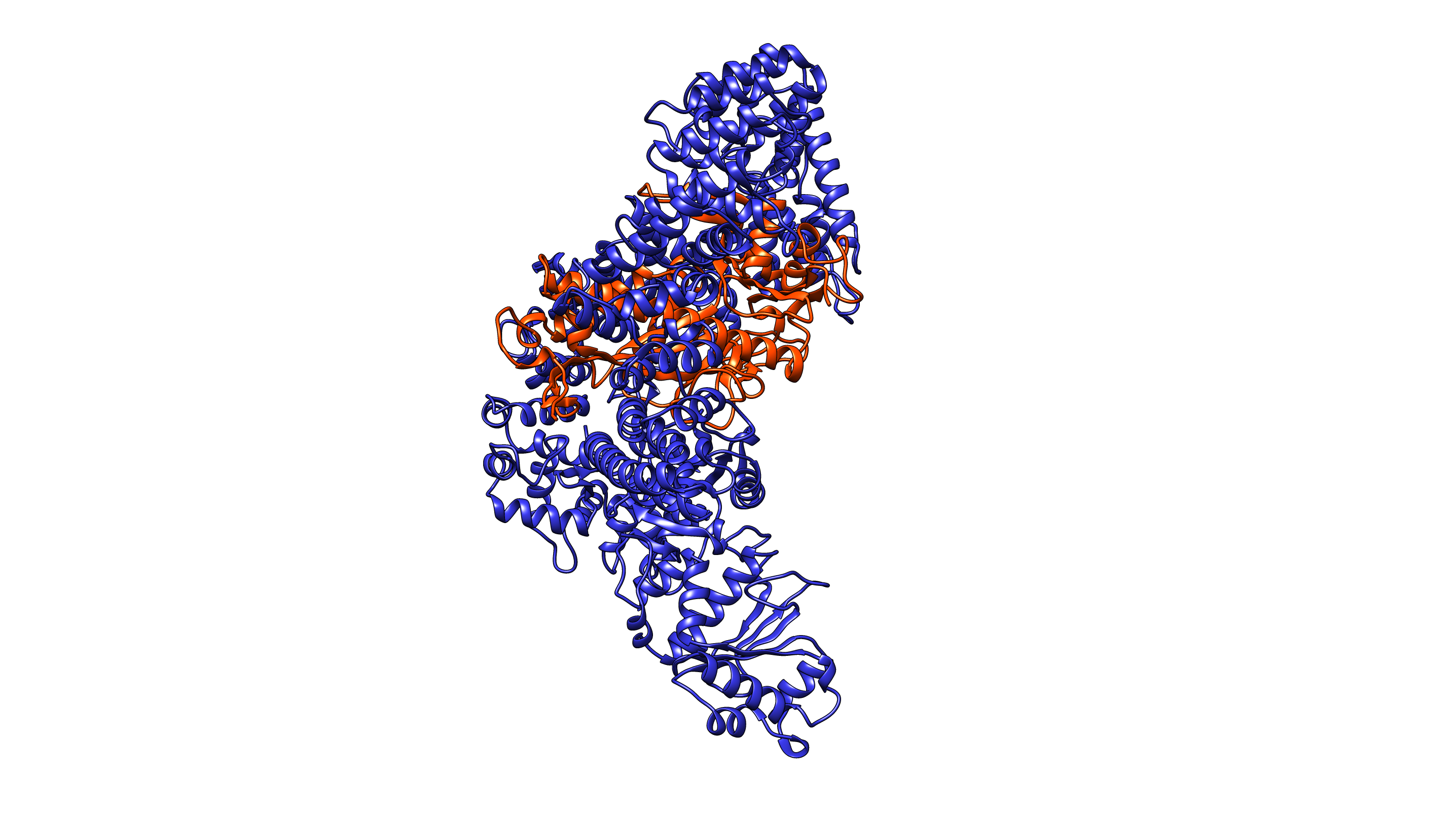

### darkblue orange surface 90 degrees.png

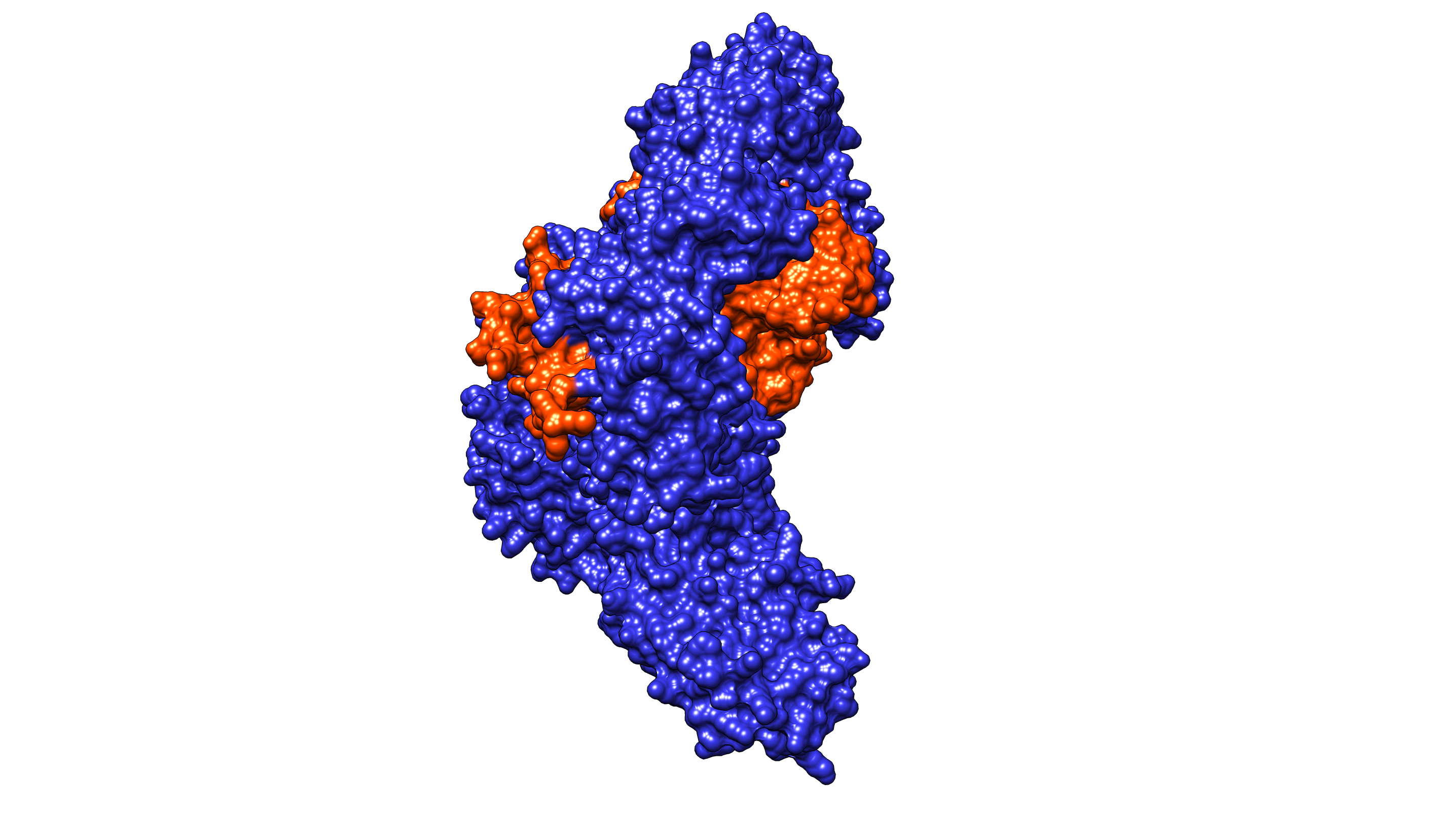

### darkblue orange surface.png

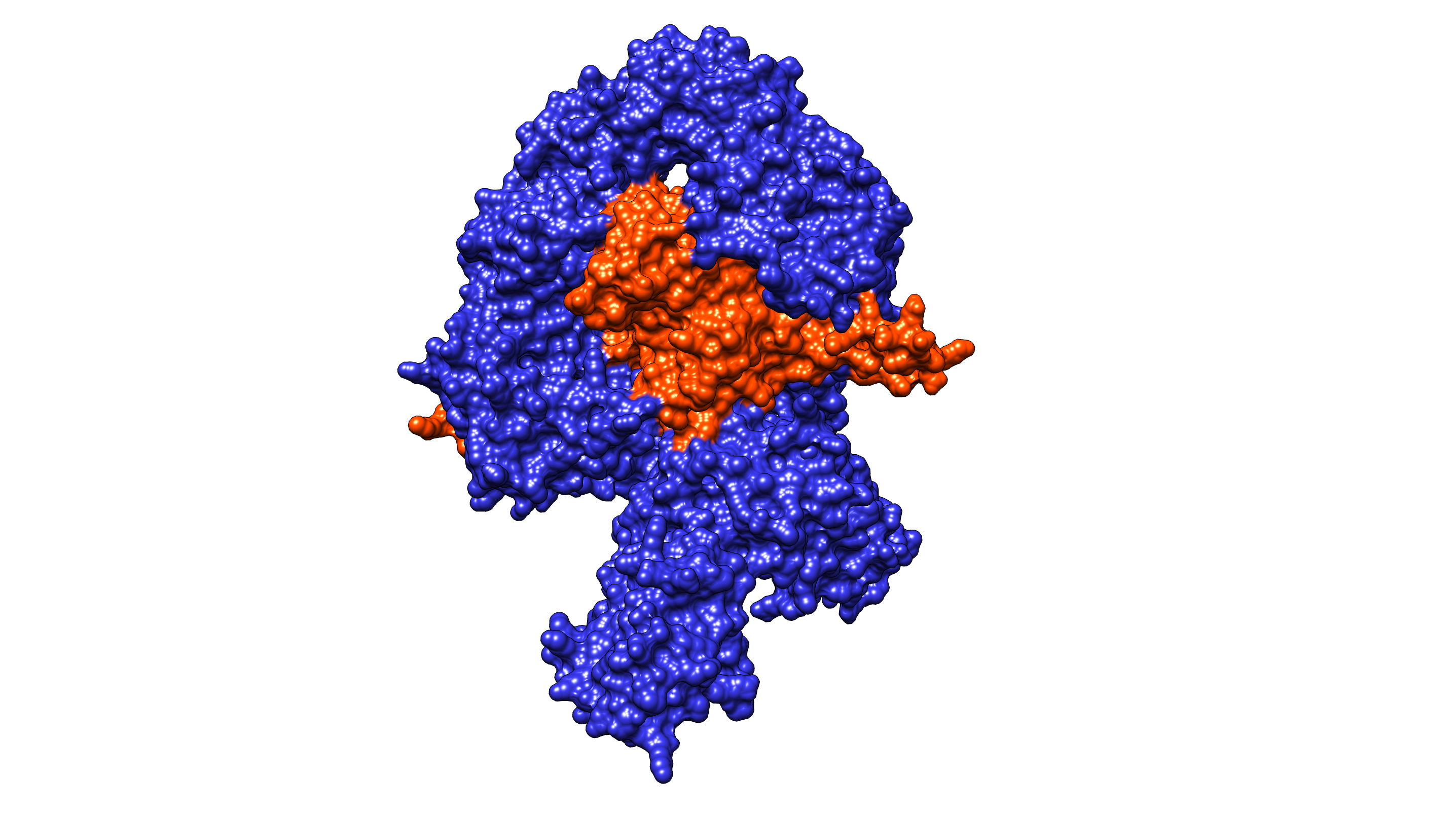

### darkblue orange.png

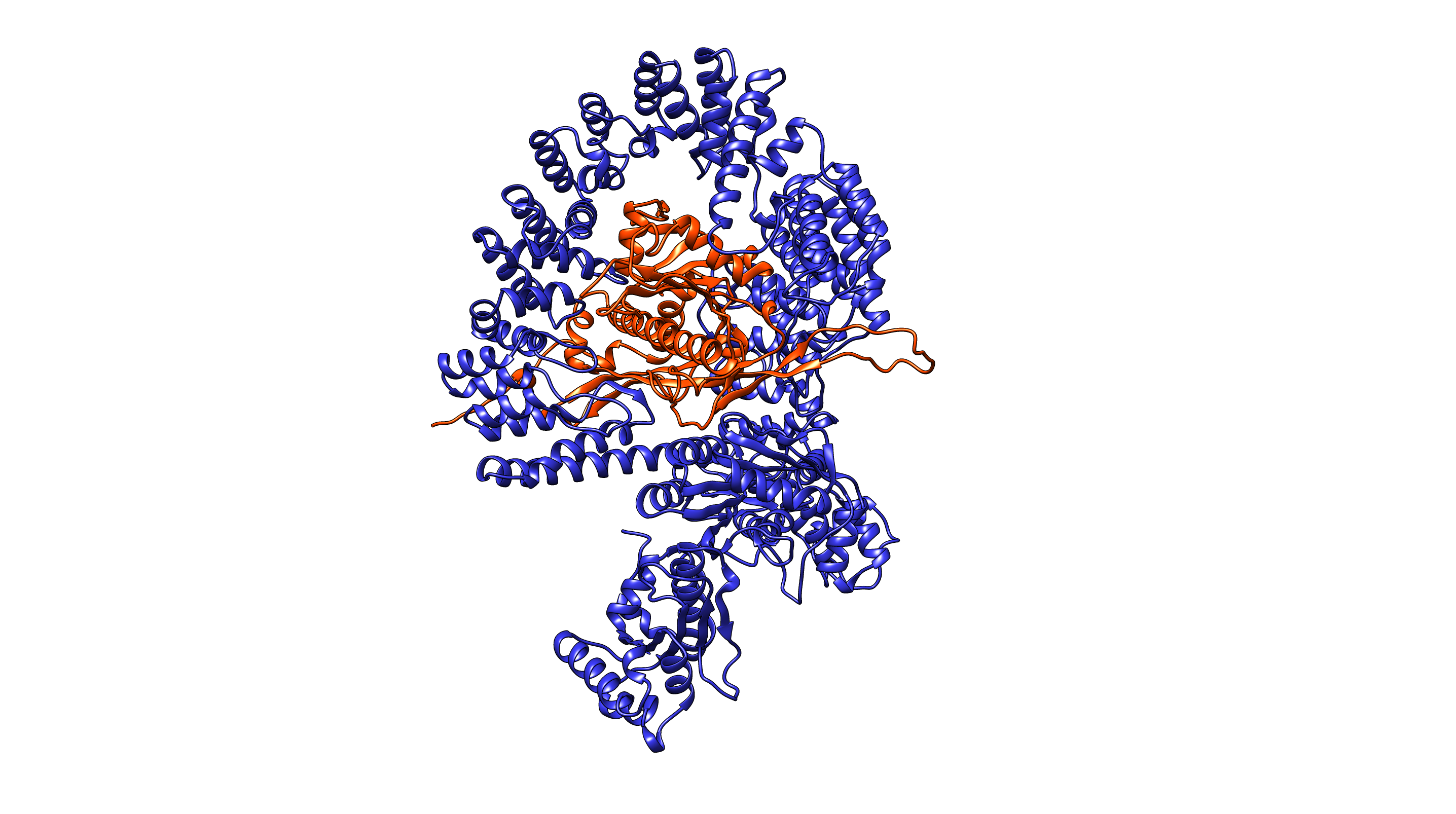

### image.png

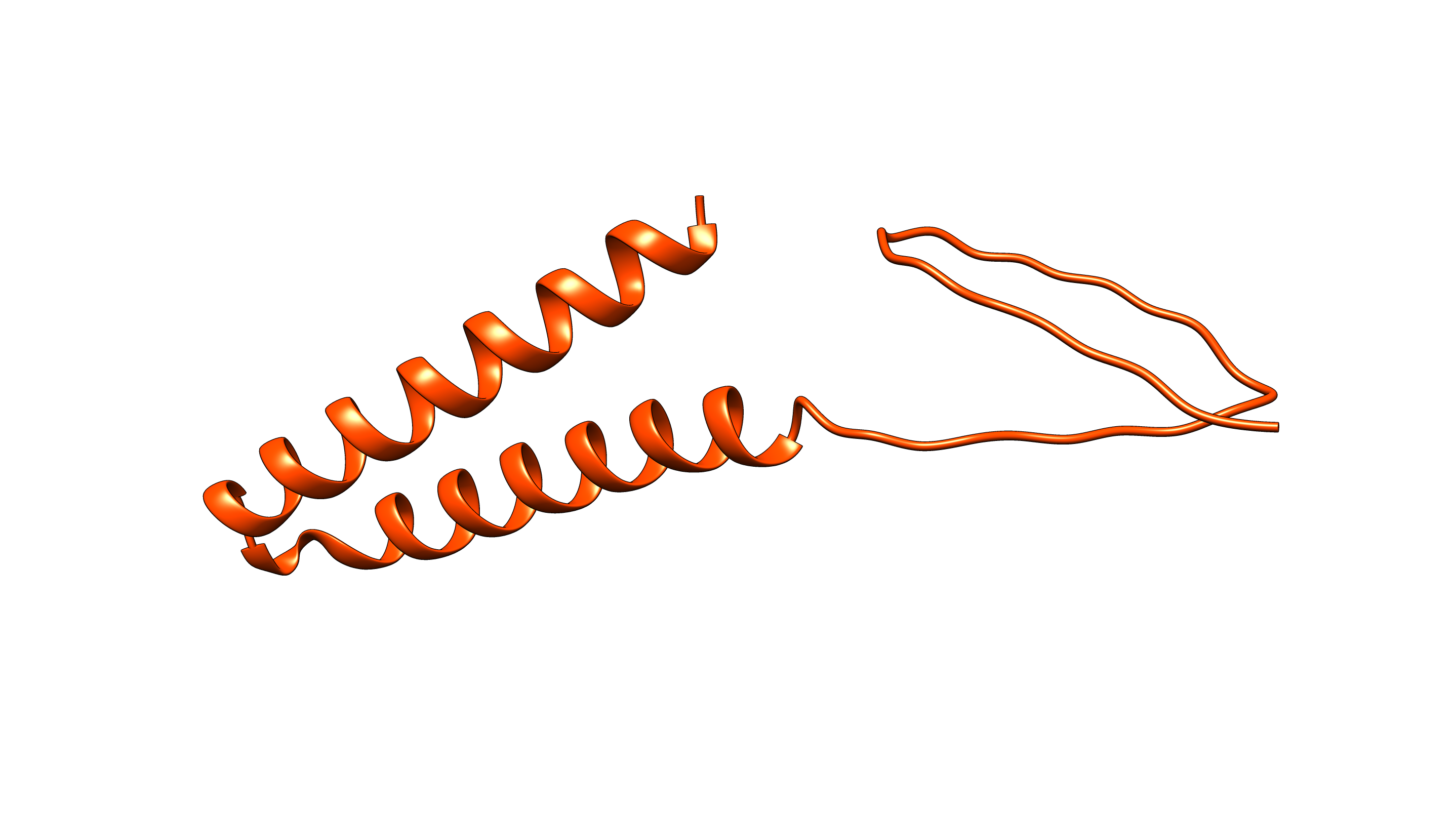

### image.png

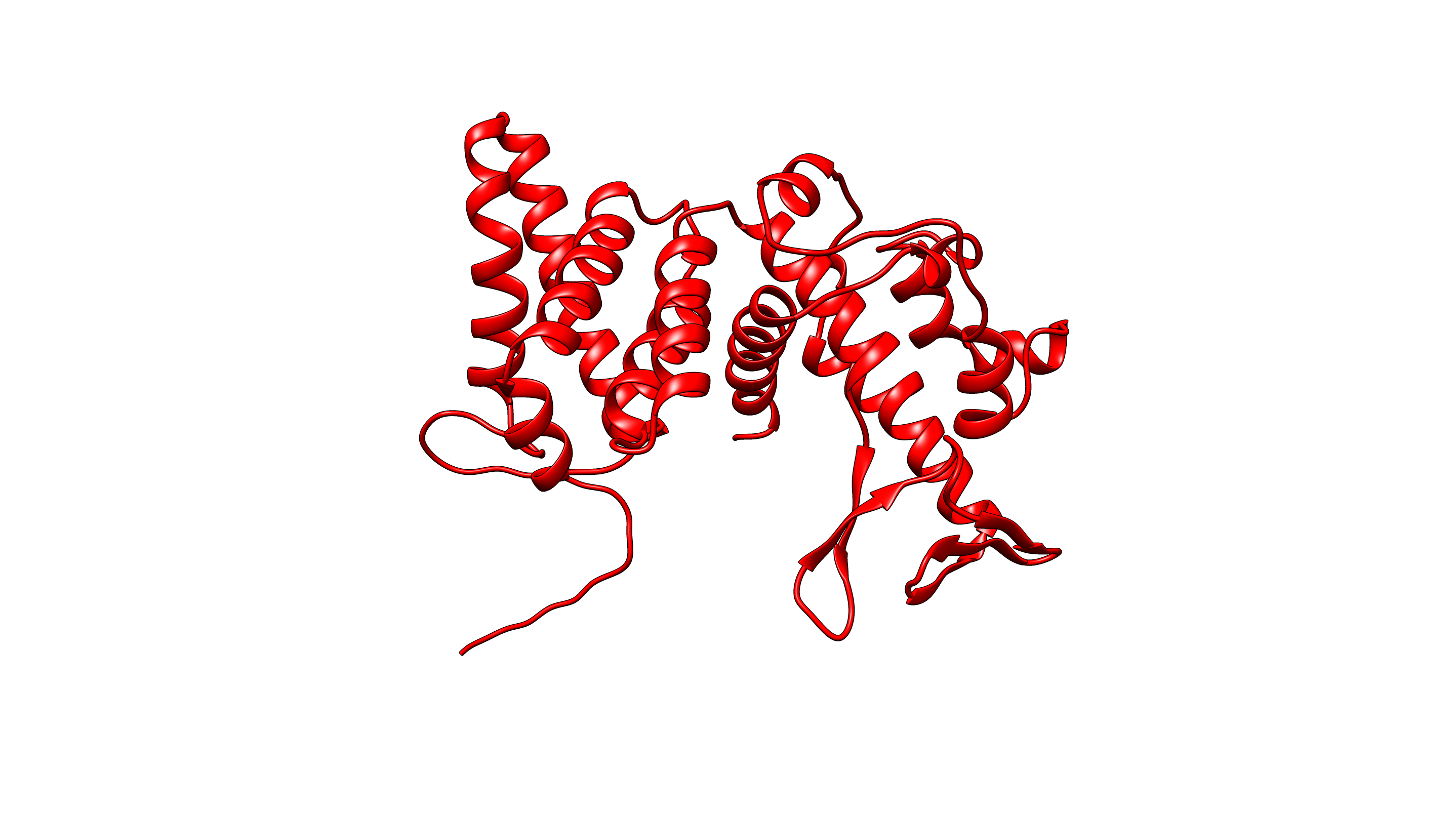

### image.png

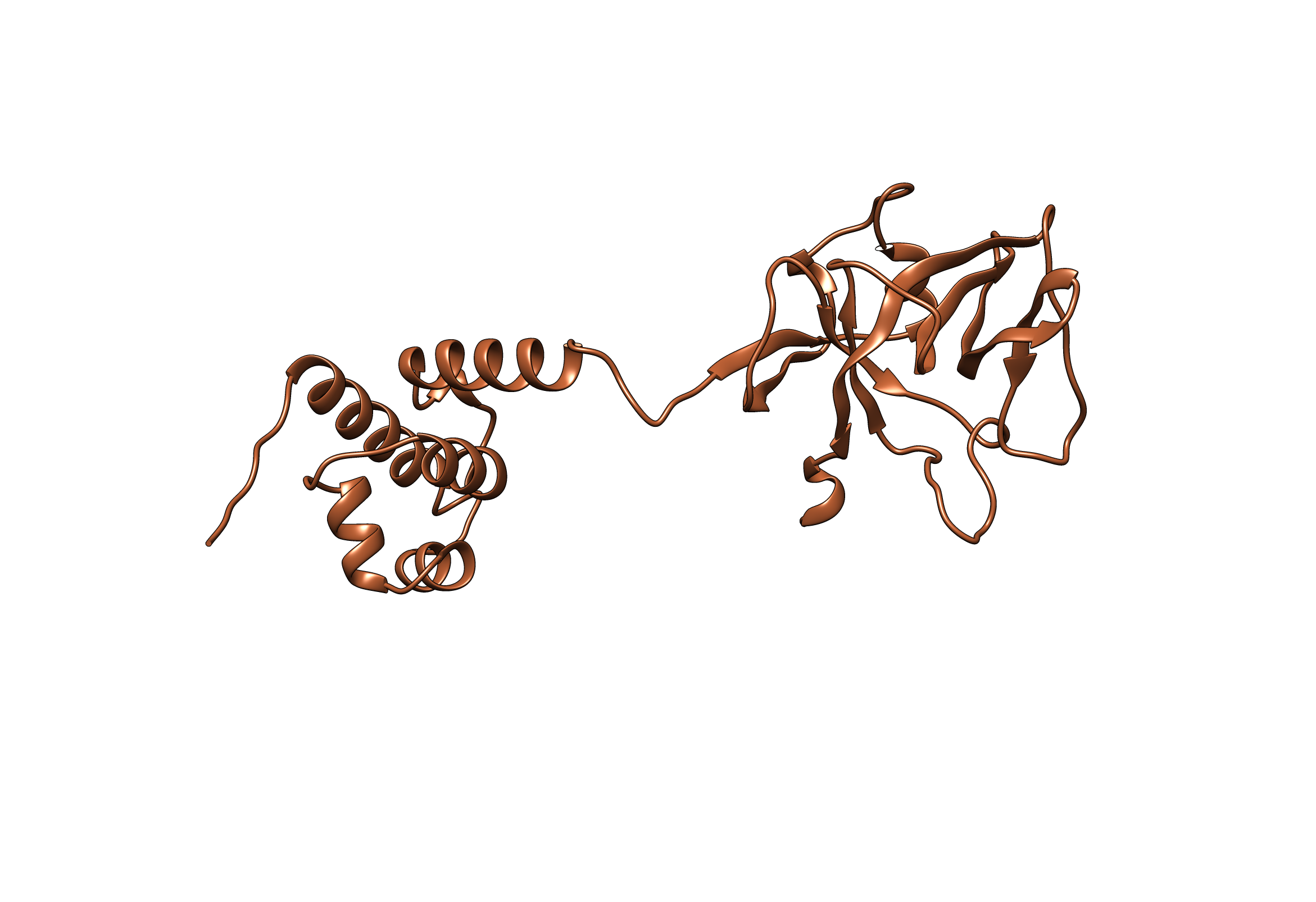

### image.png

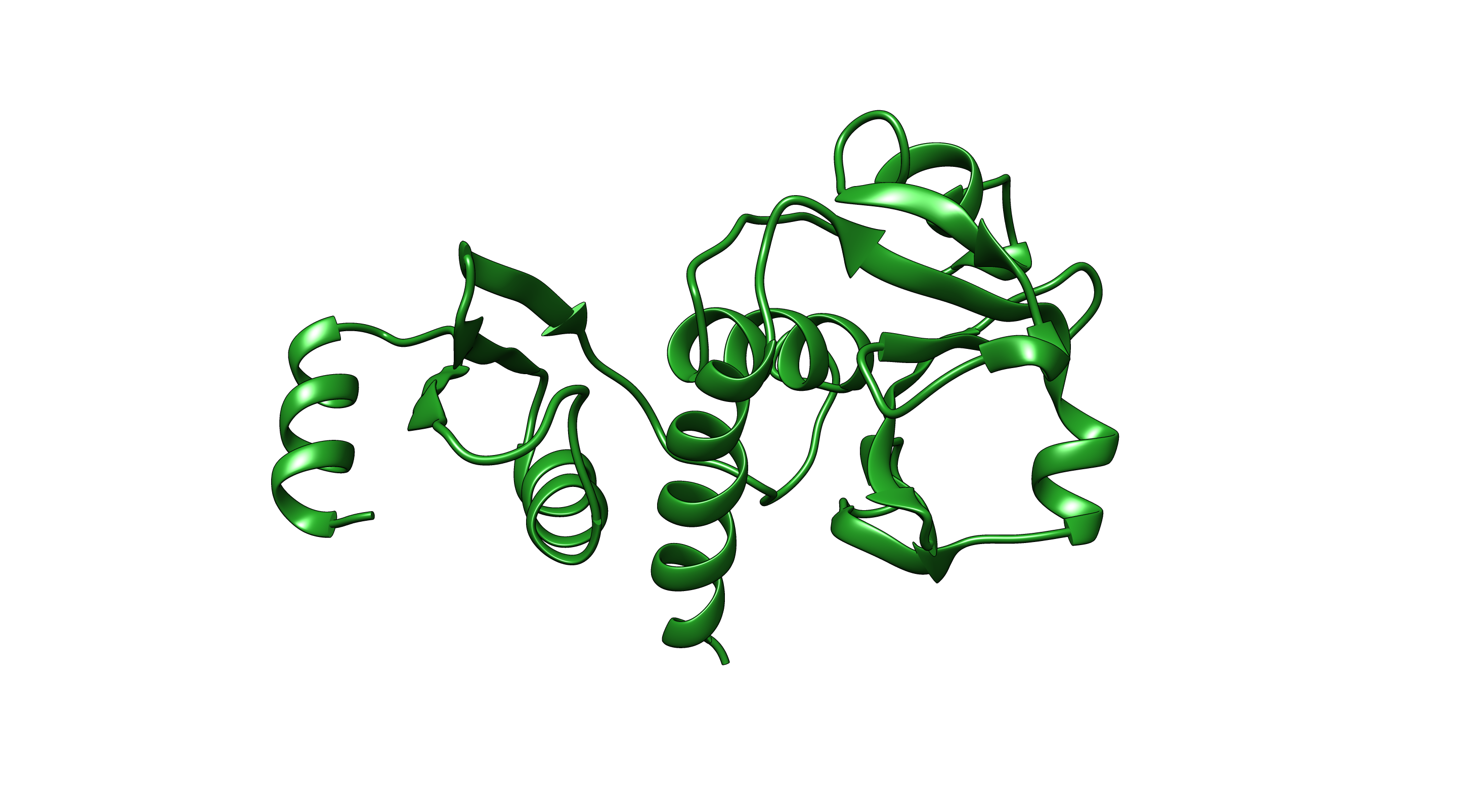

### image.png

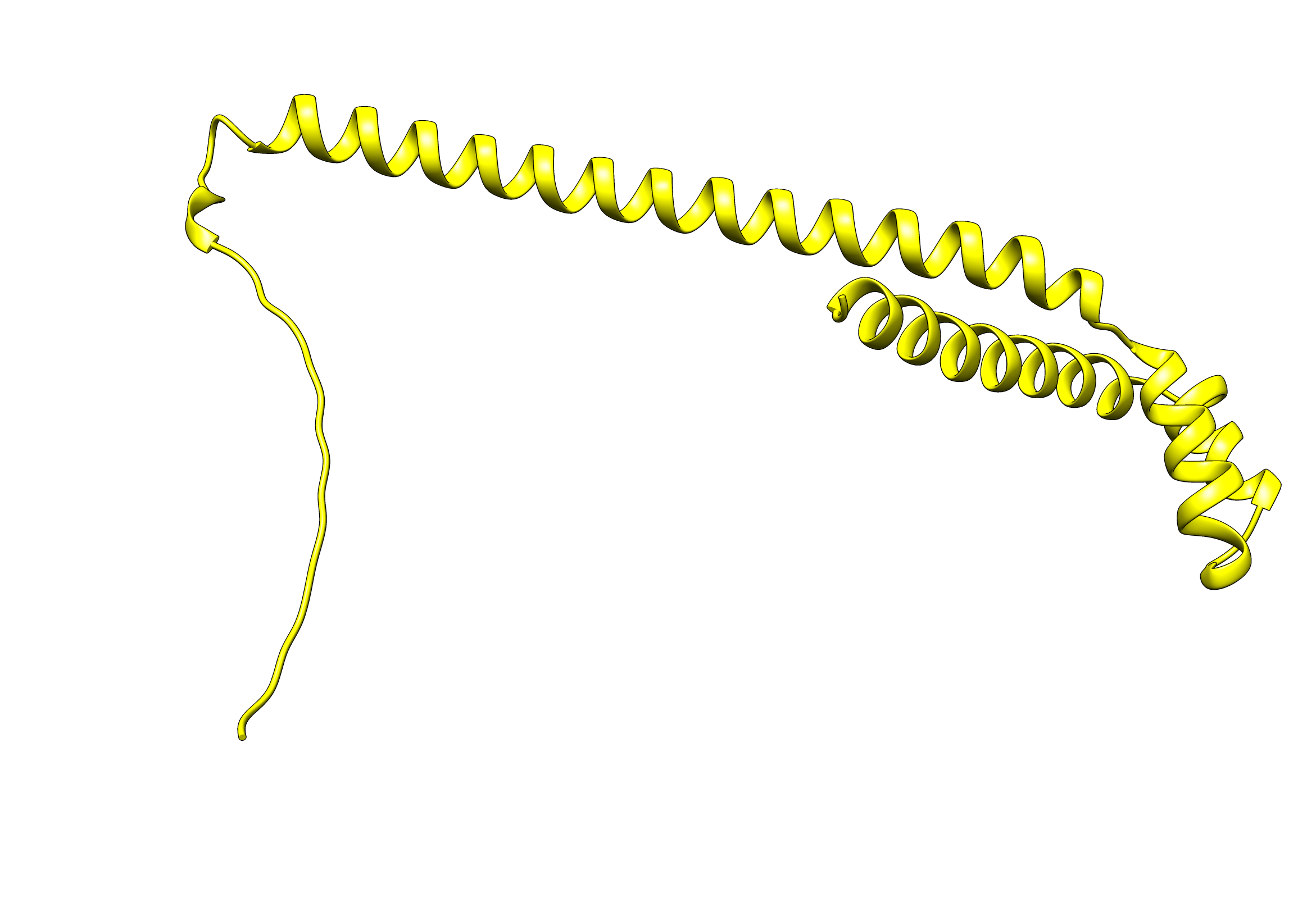

### image.png

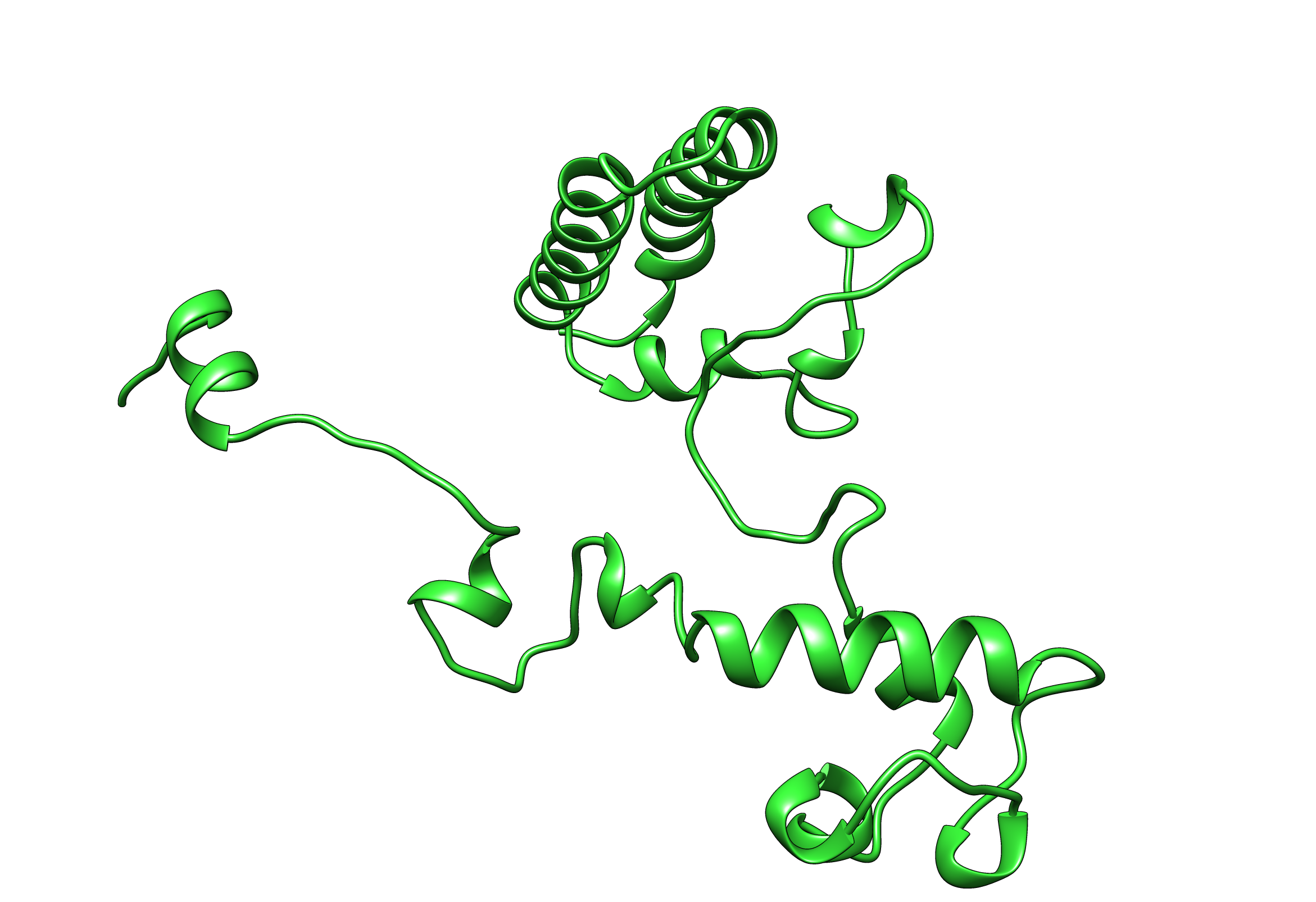

### image.png

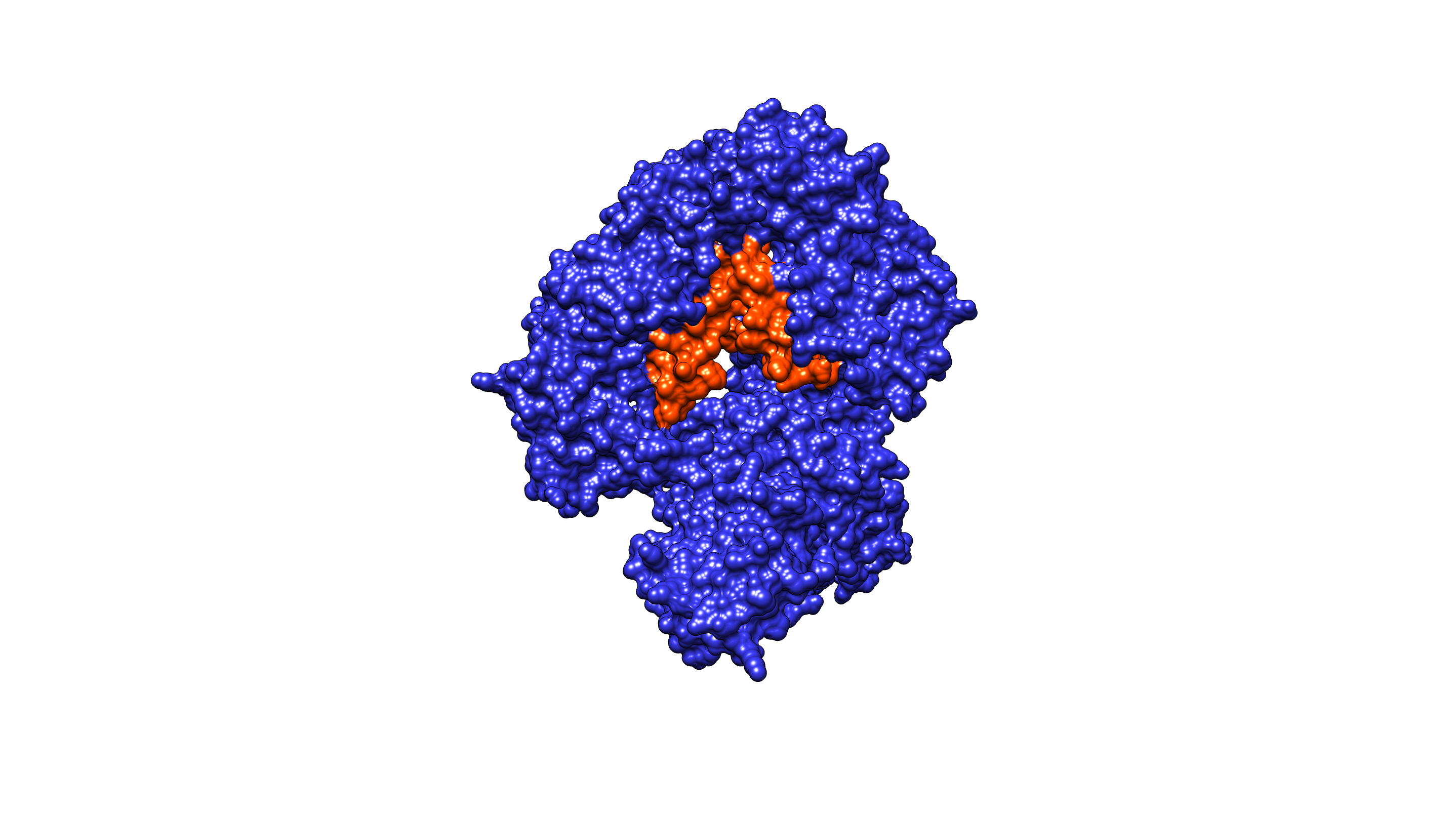

### image.png

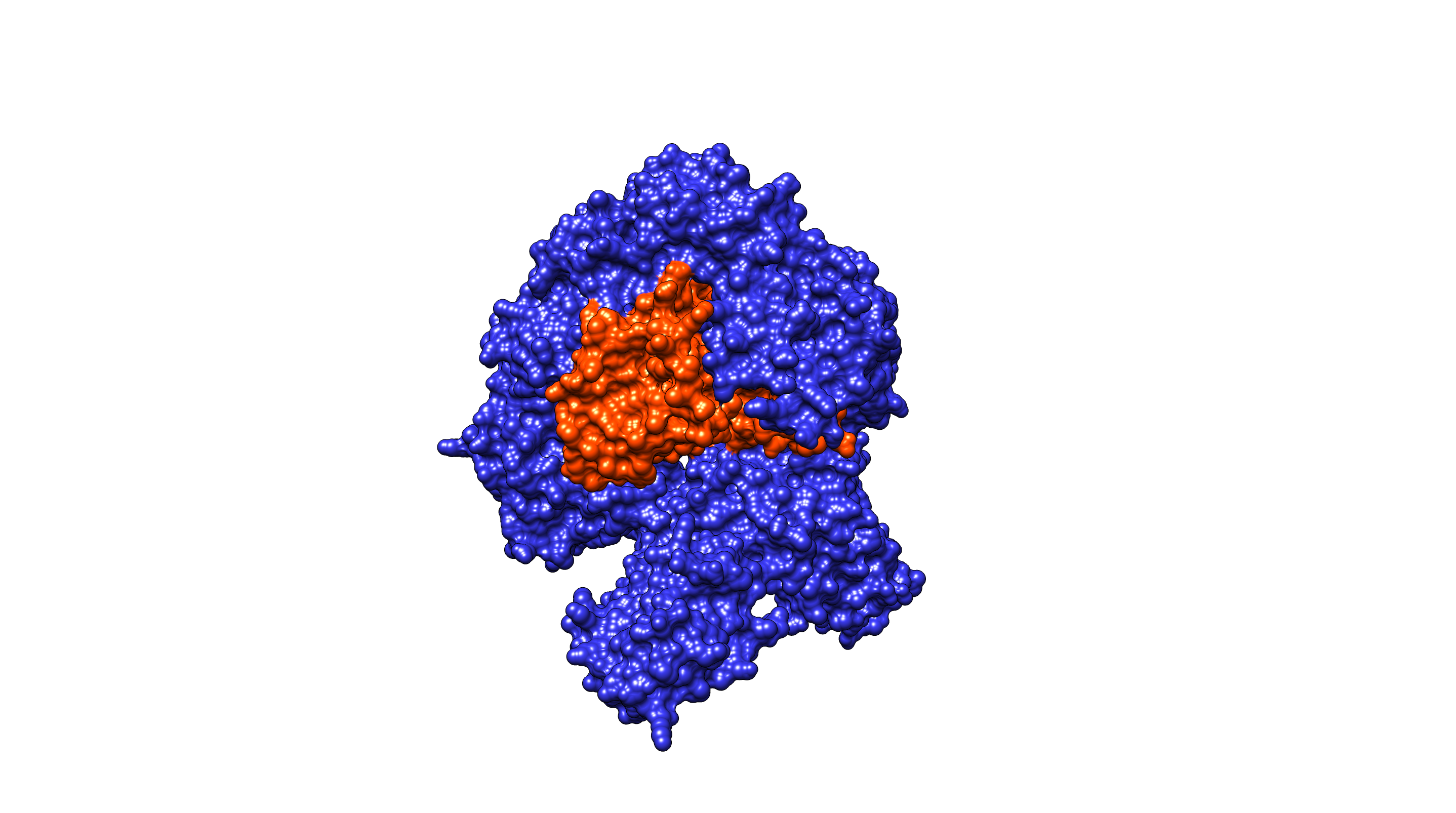

### image.png

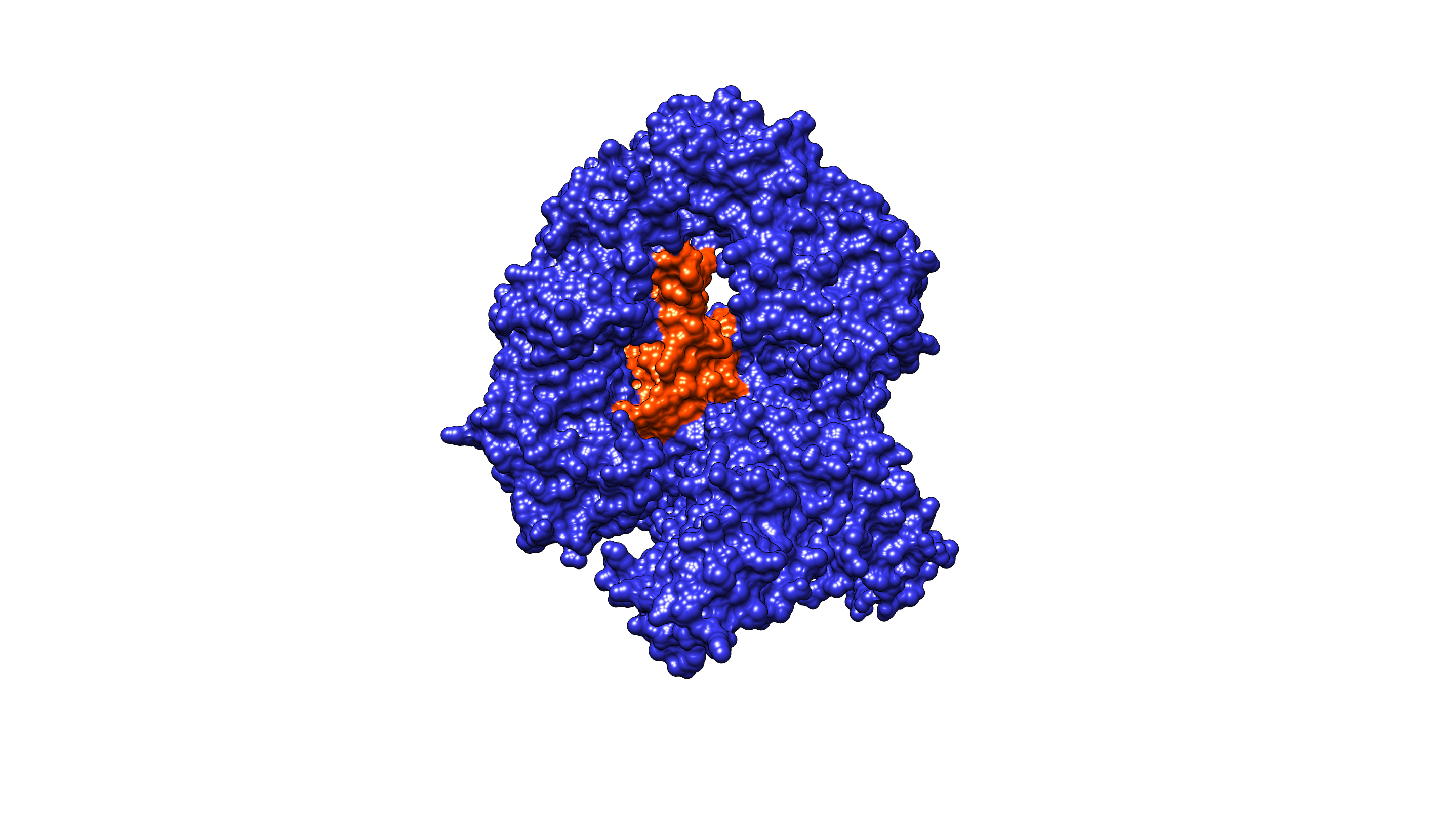

### image.png

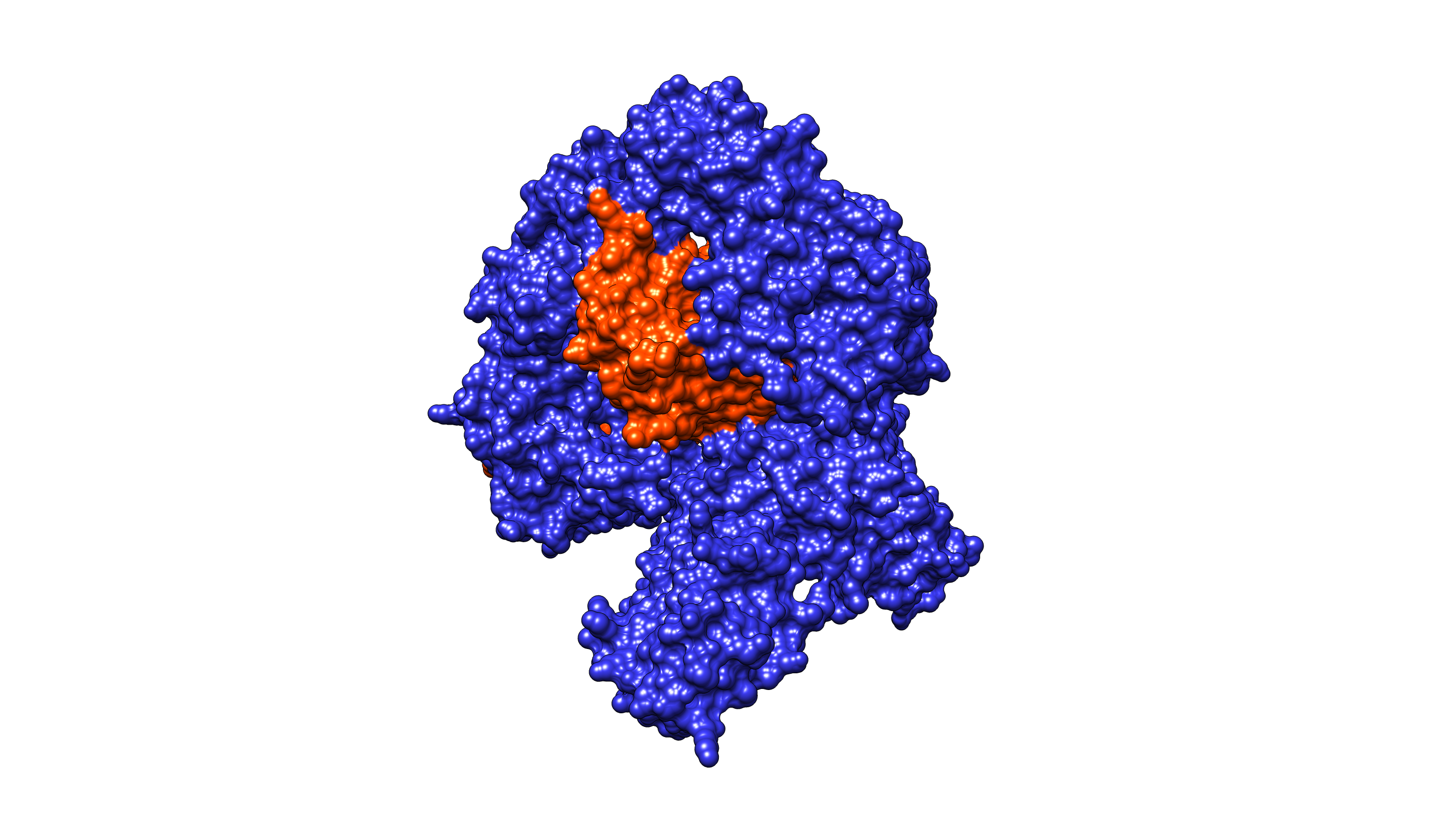

### image.png

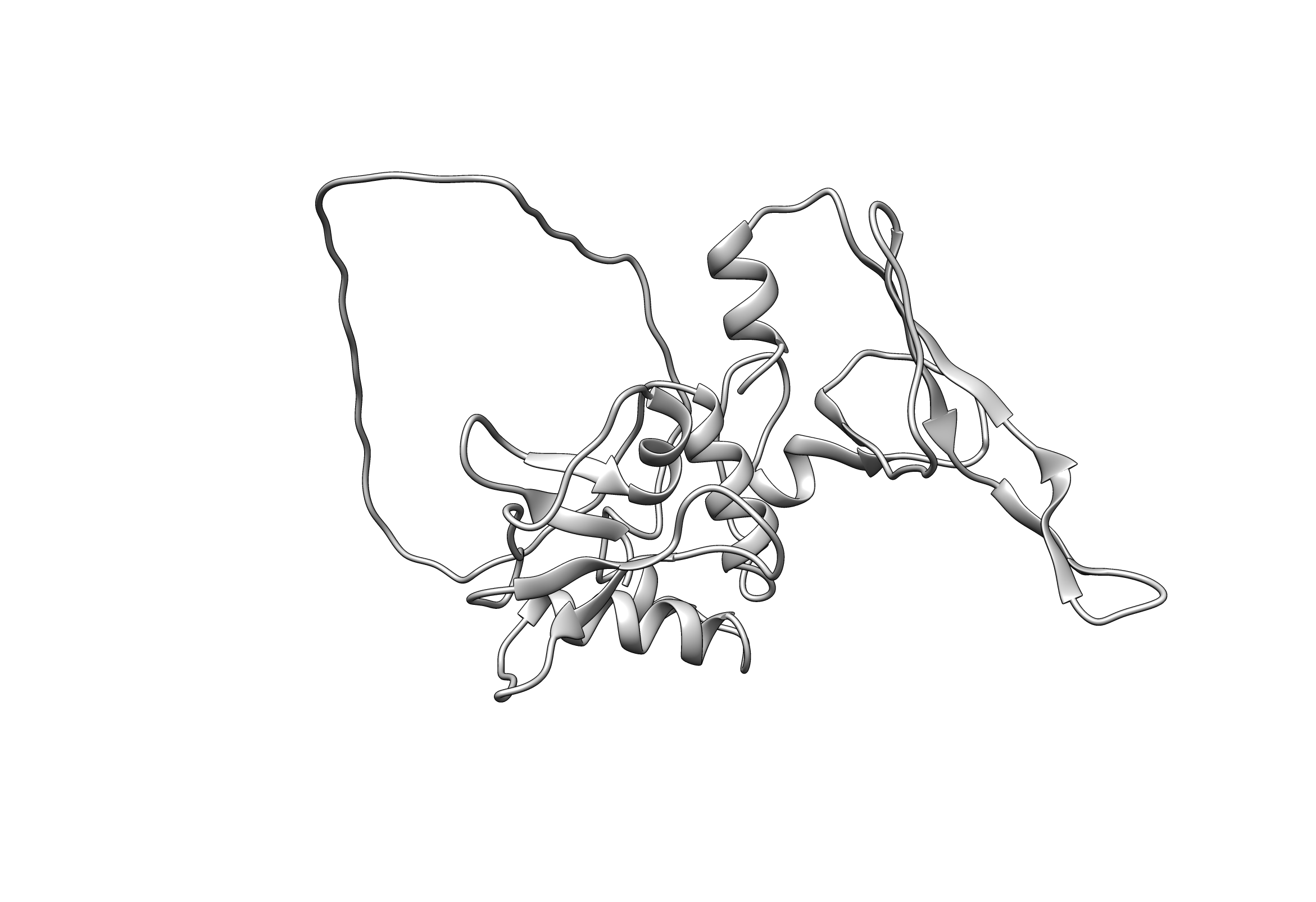

### image.png

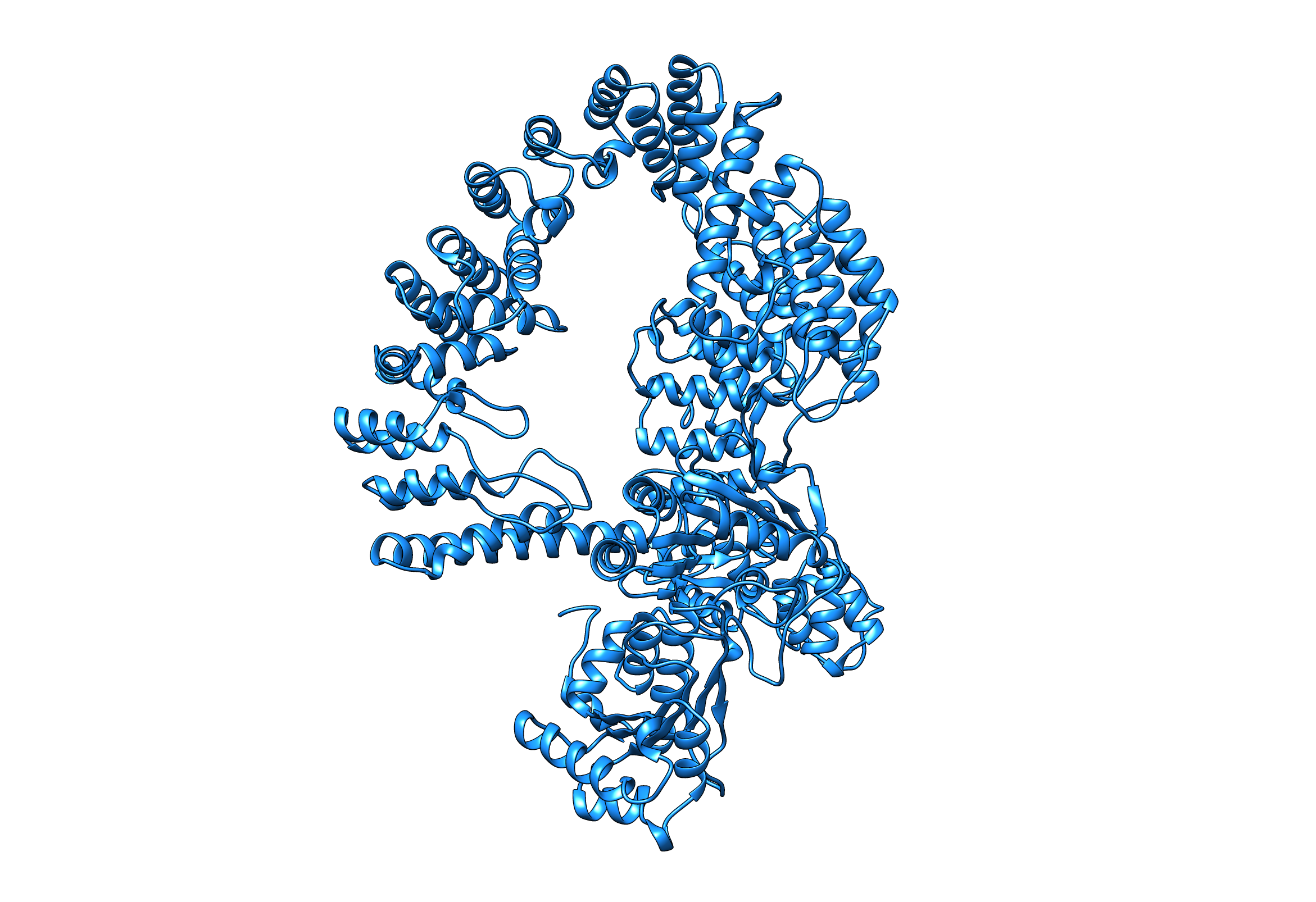

### image.png

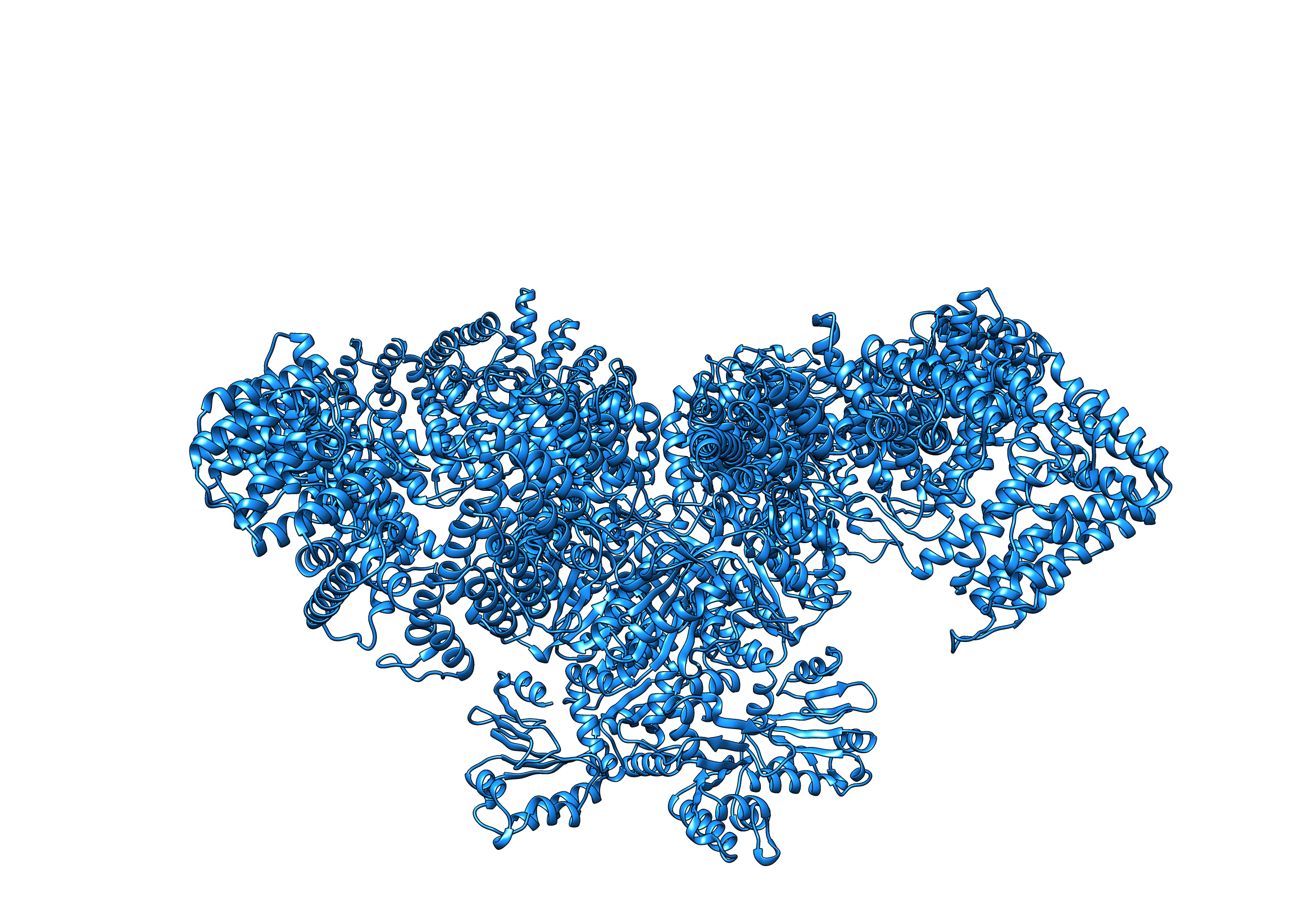

### image.png

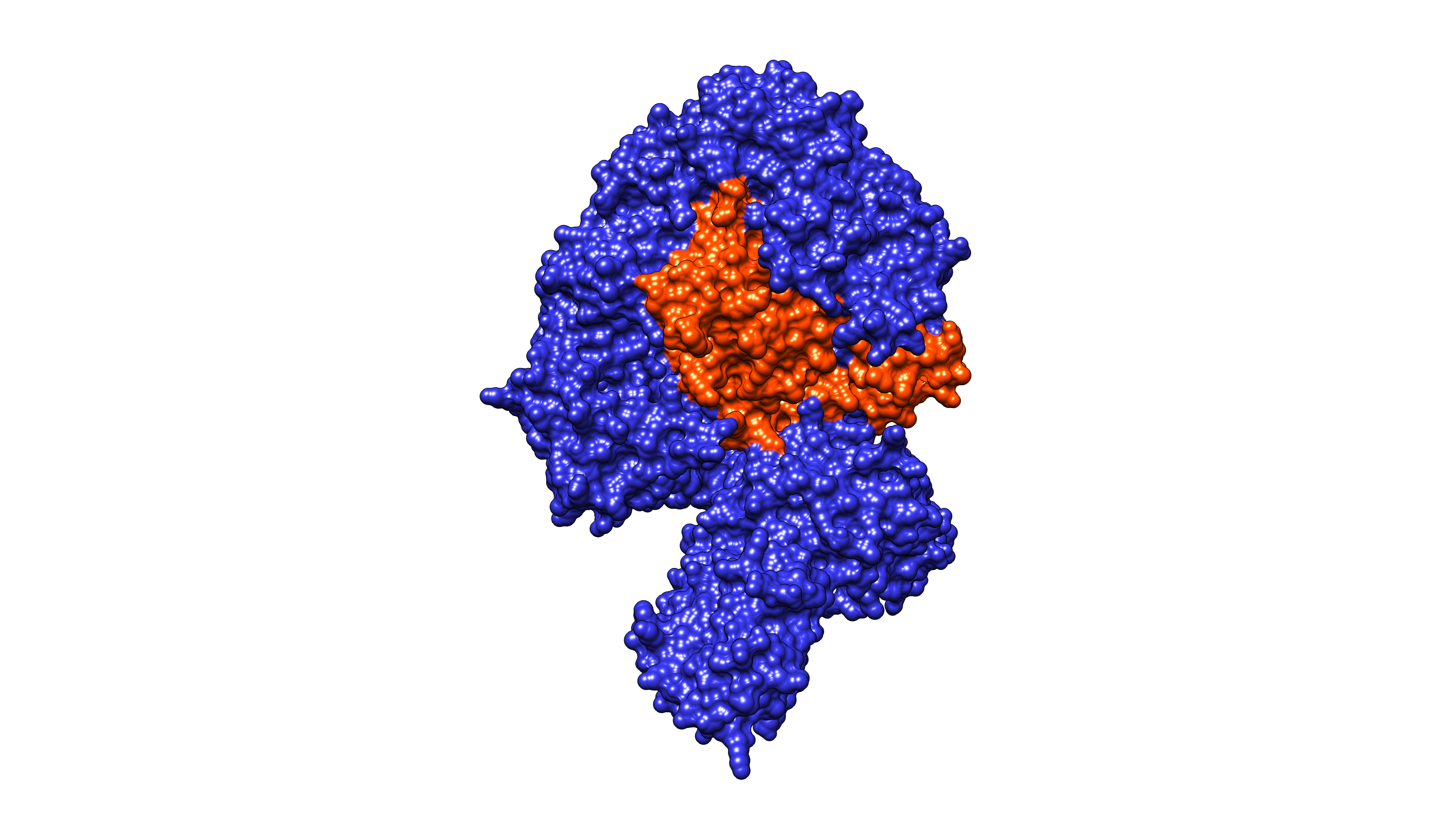

### image.png

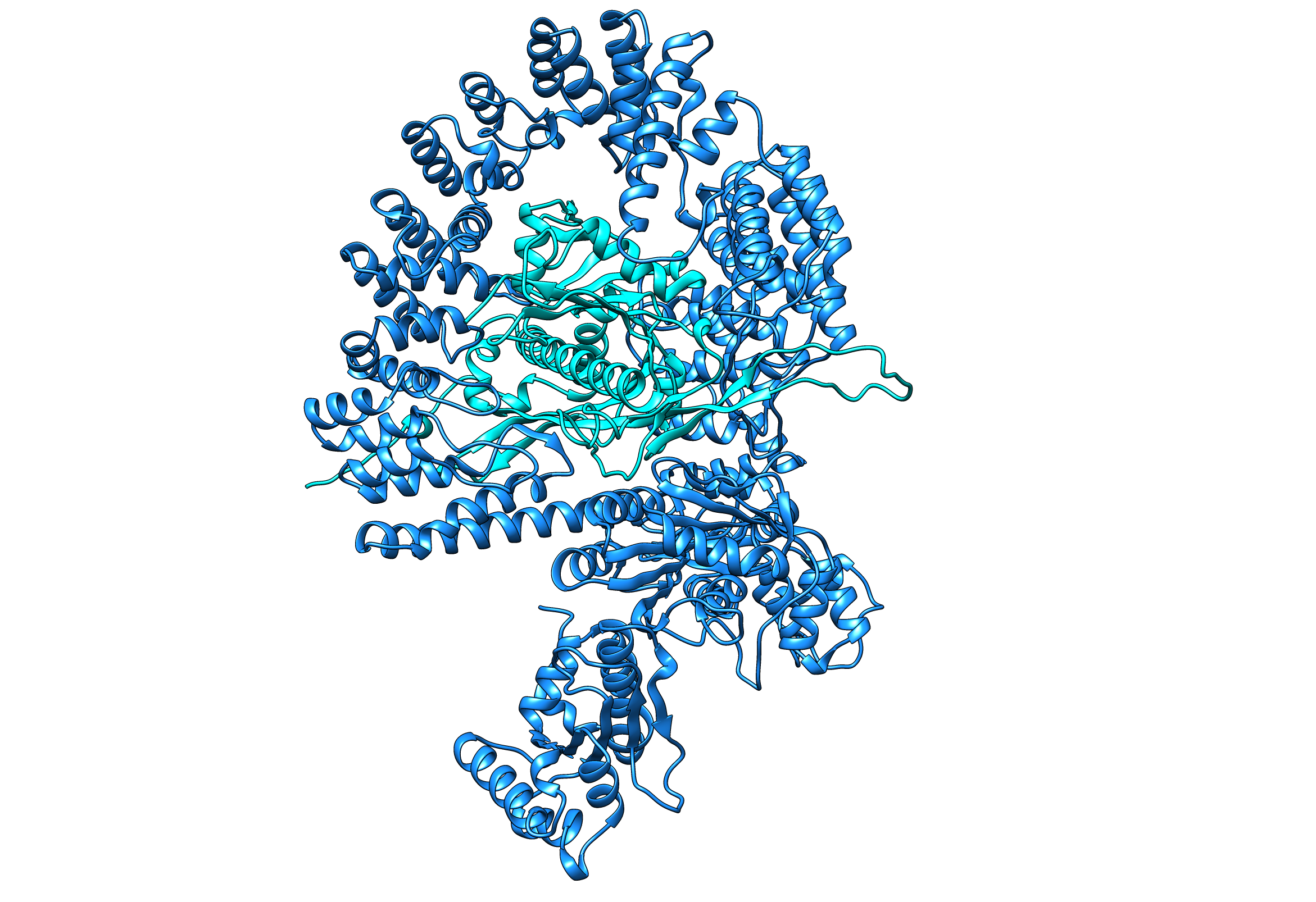

### image.png

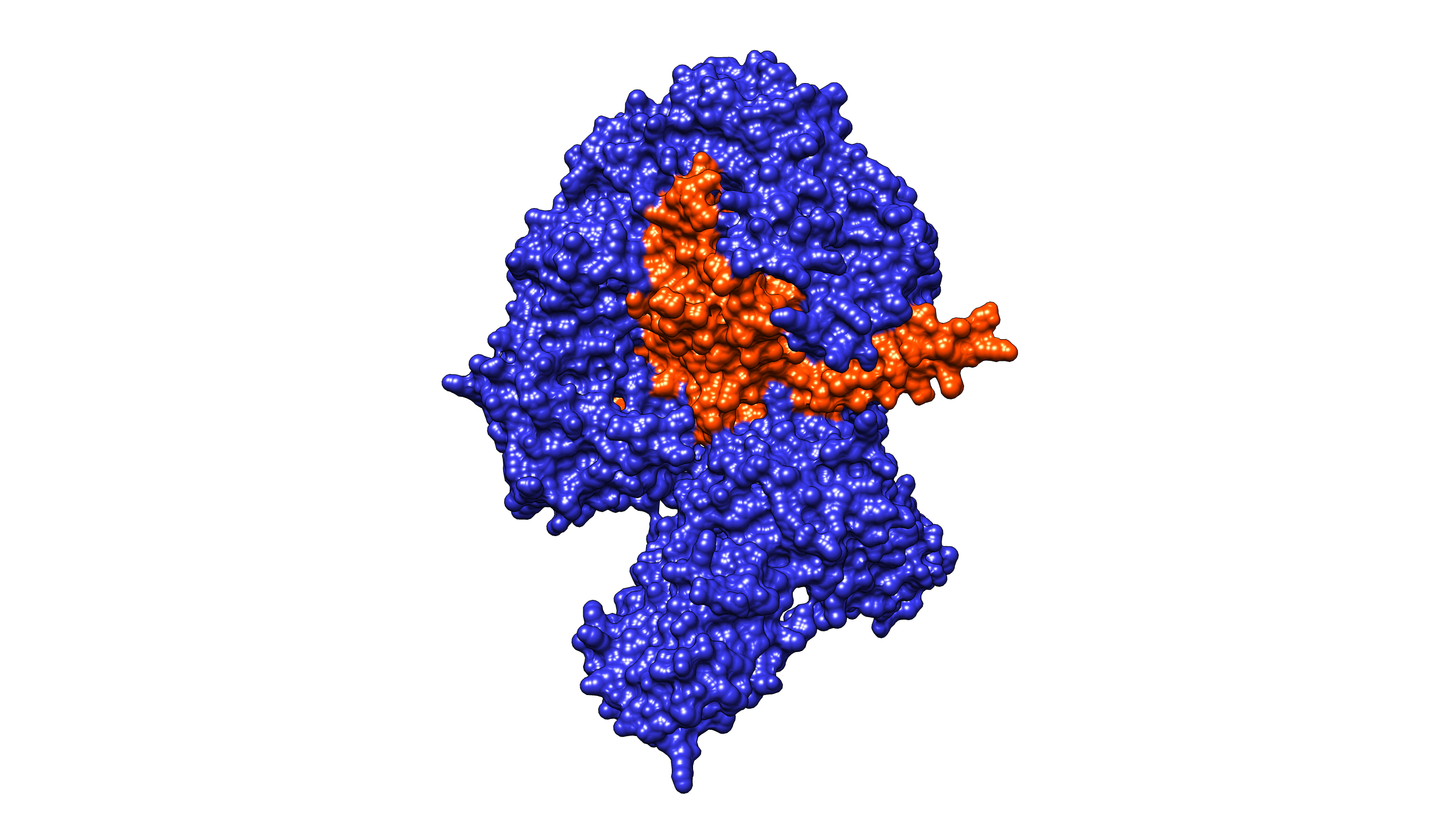

### image.png

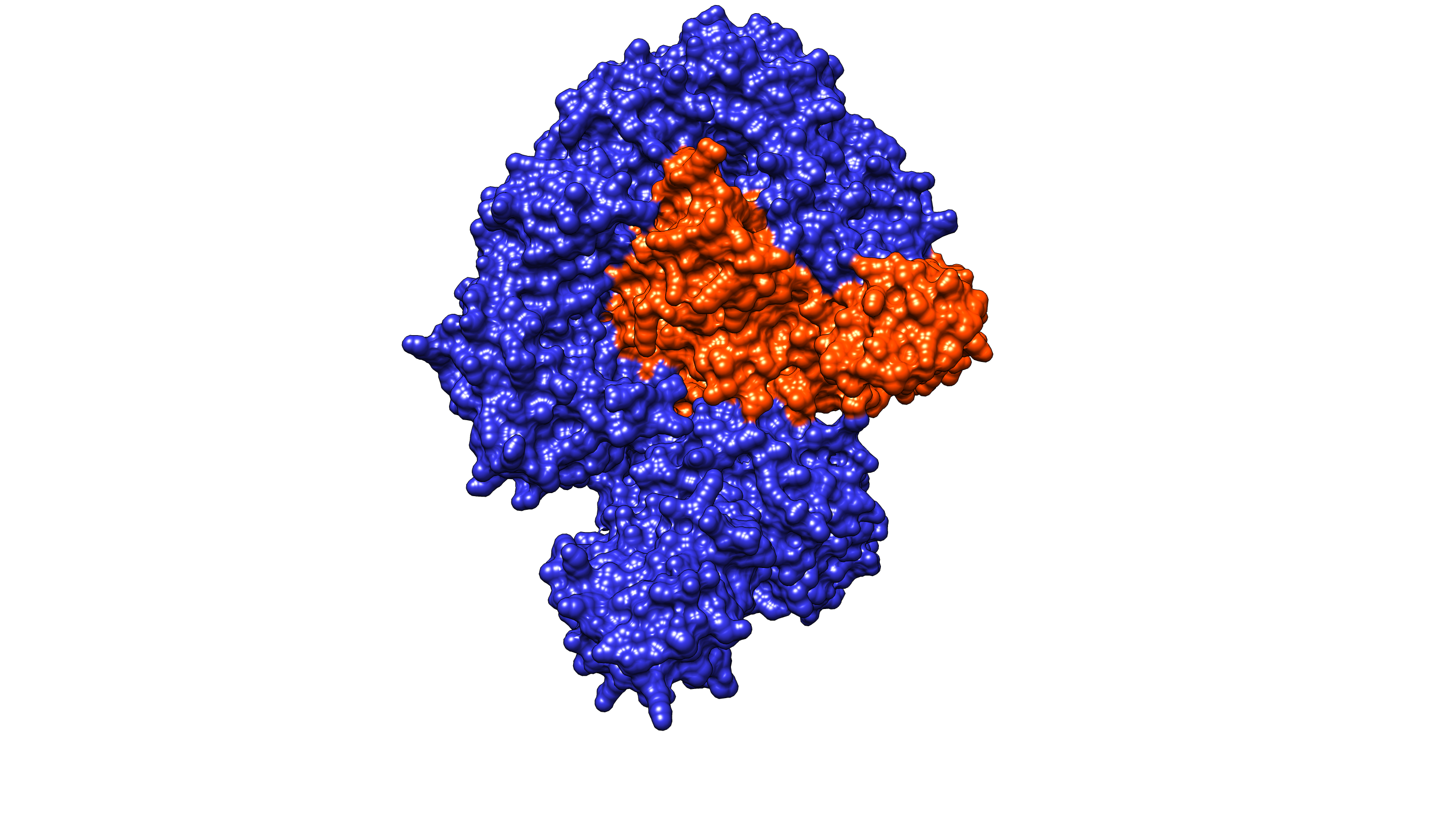

### image.png

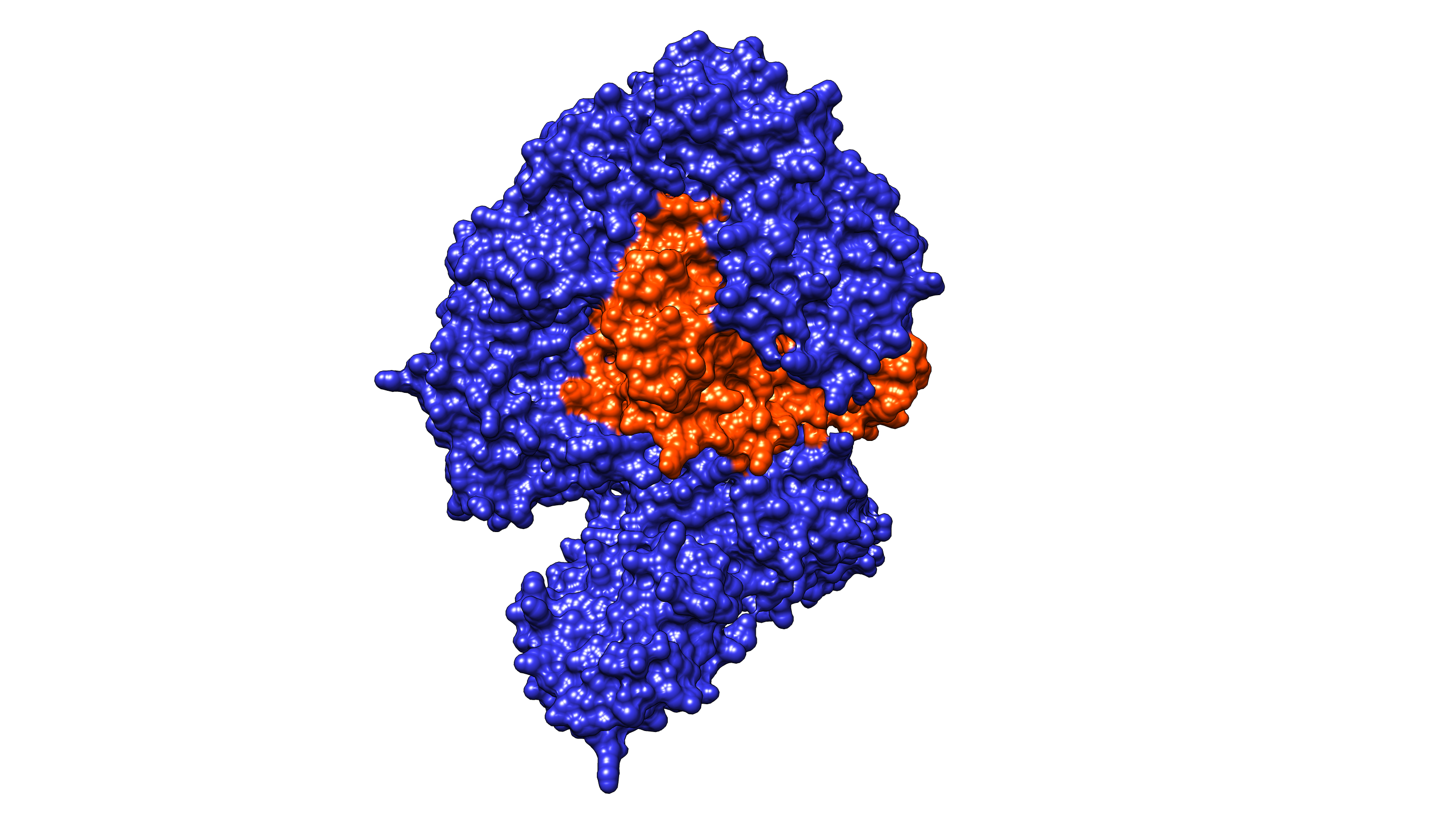

### image.png

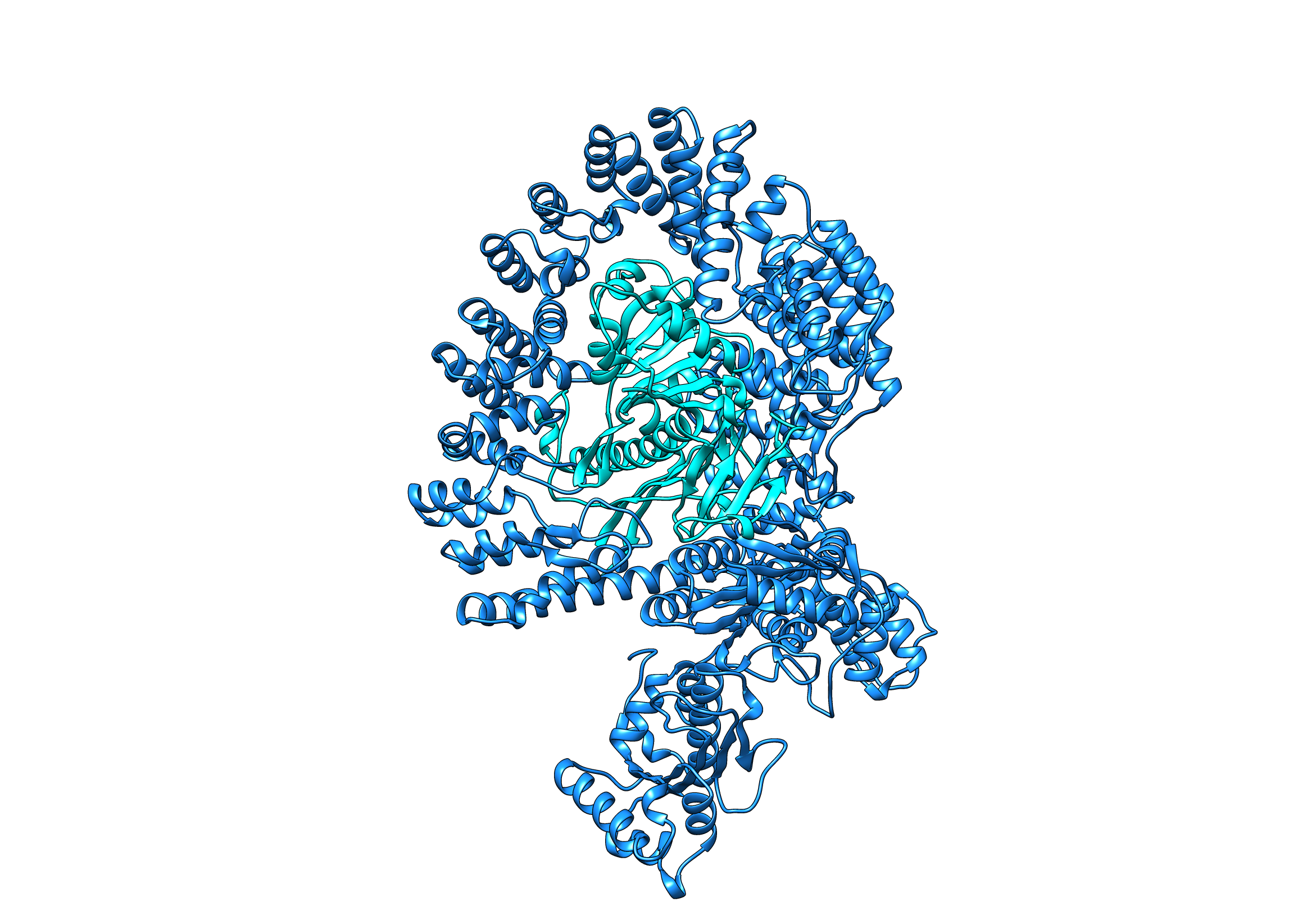

### imagerotate90.png

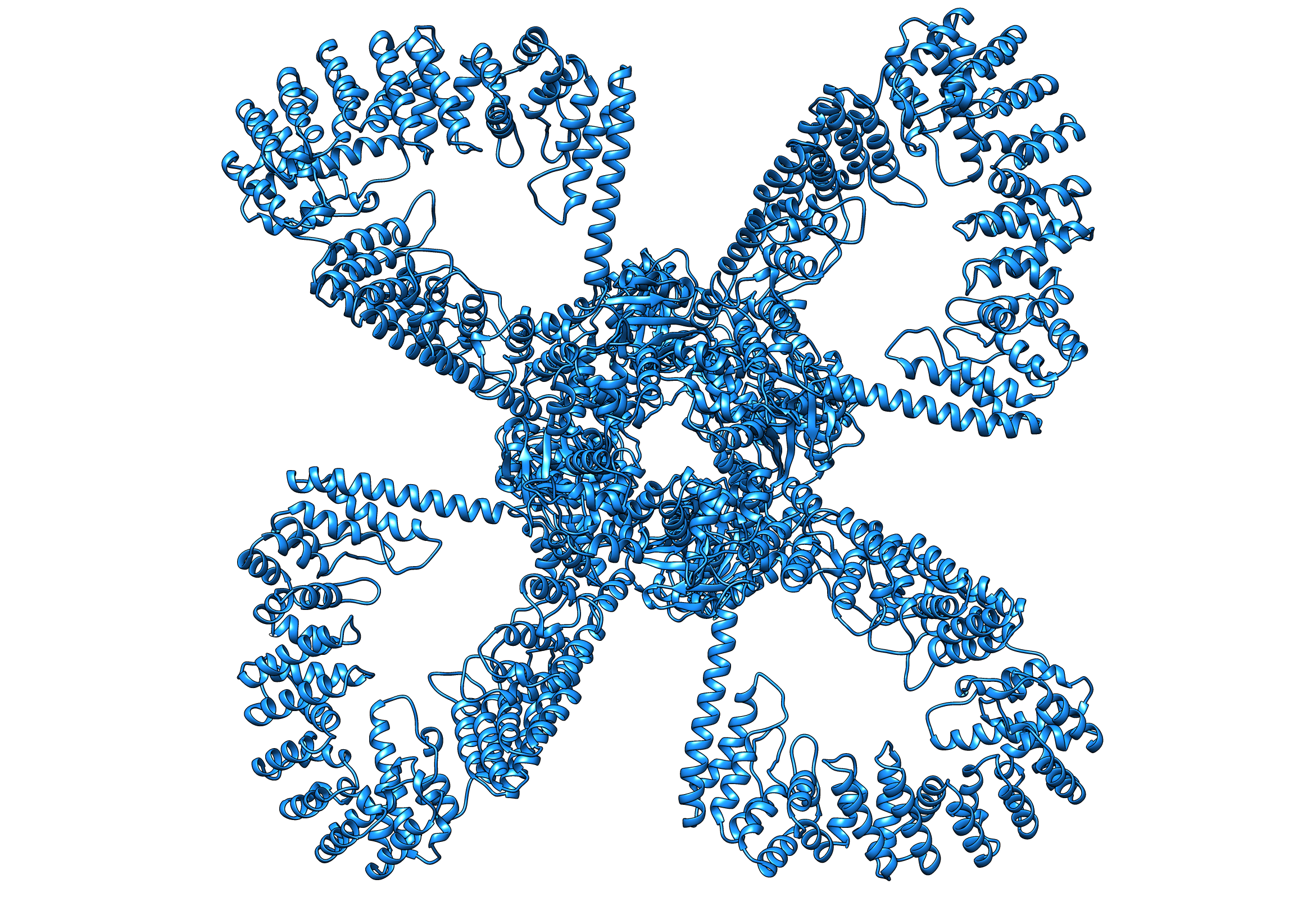

### imagerotate90.png

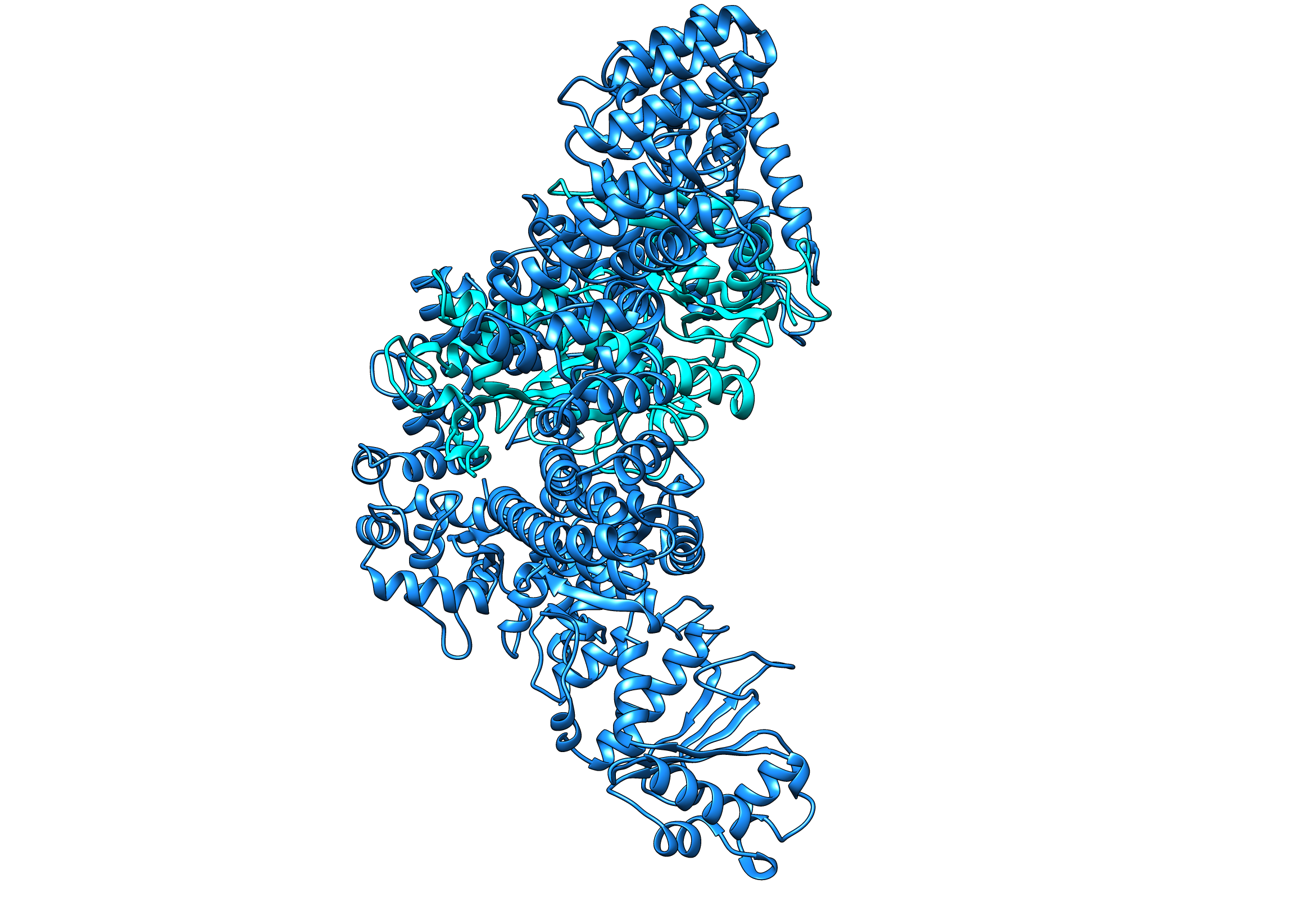

### imagerotate90.png

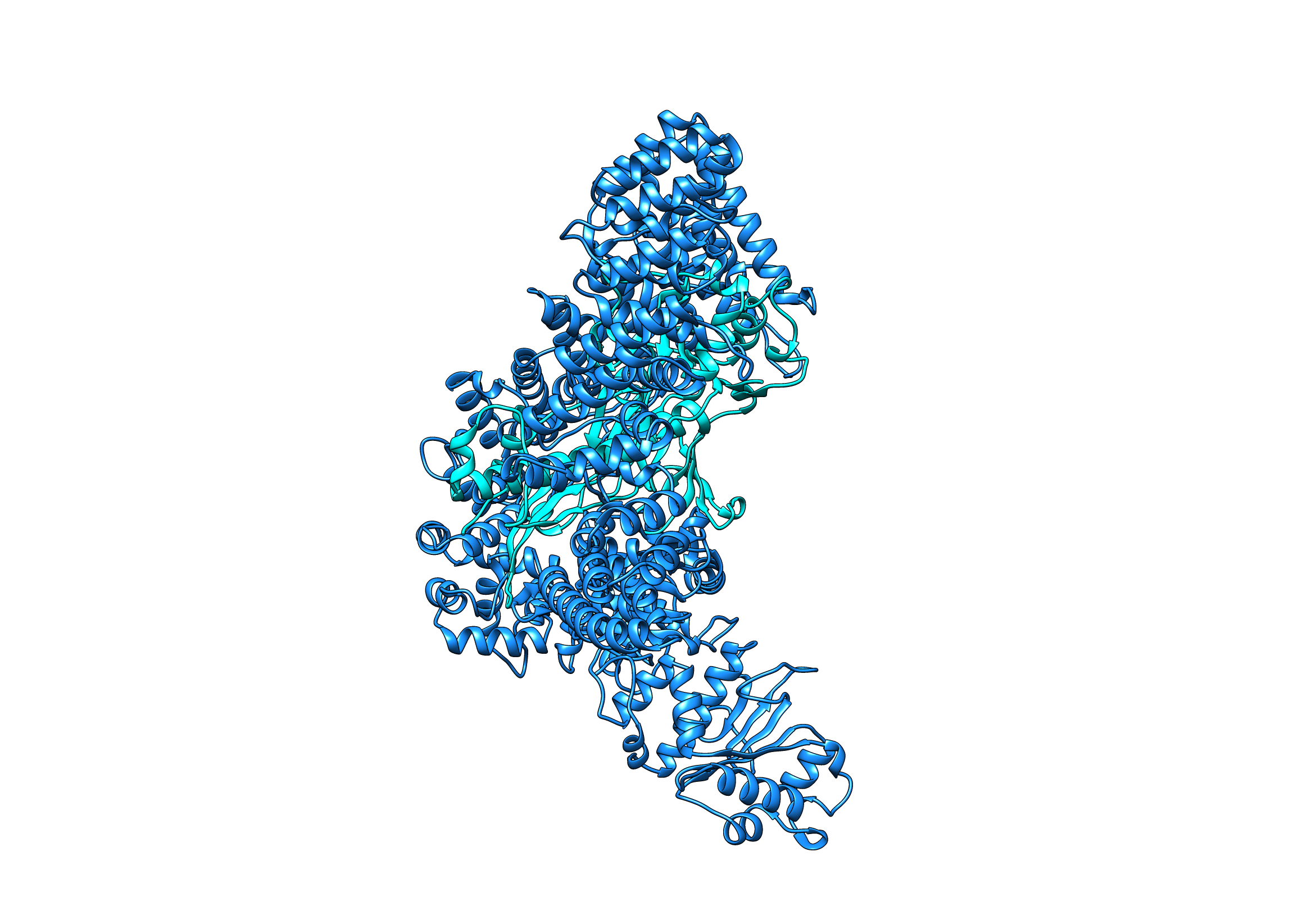

### imagespacefillin180.png

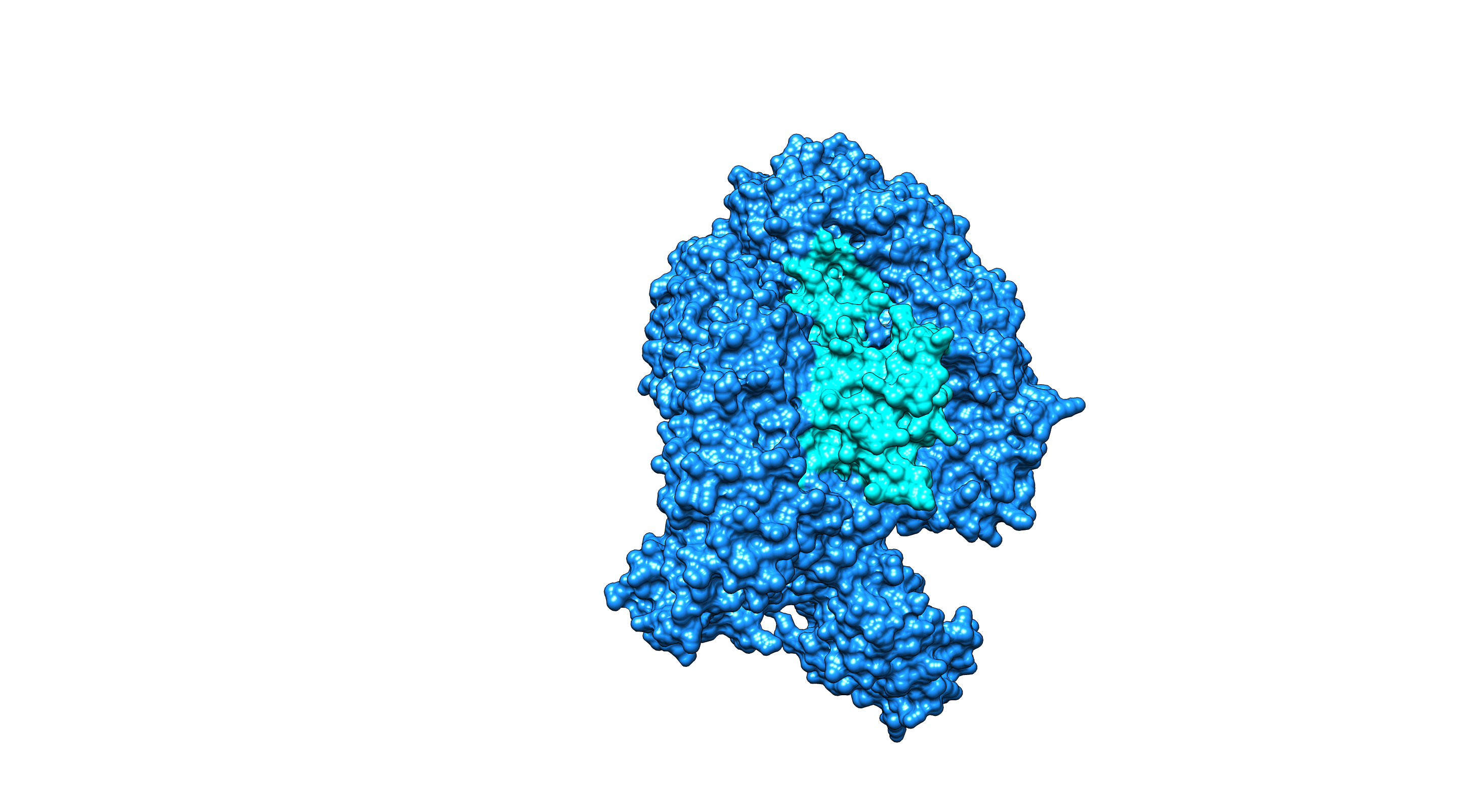

### imagespacefillin9090.png

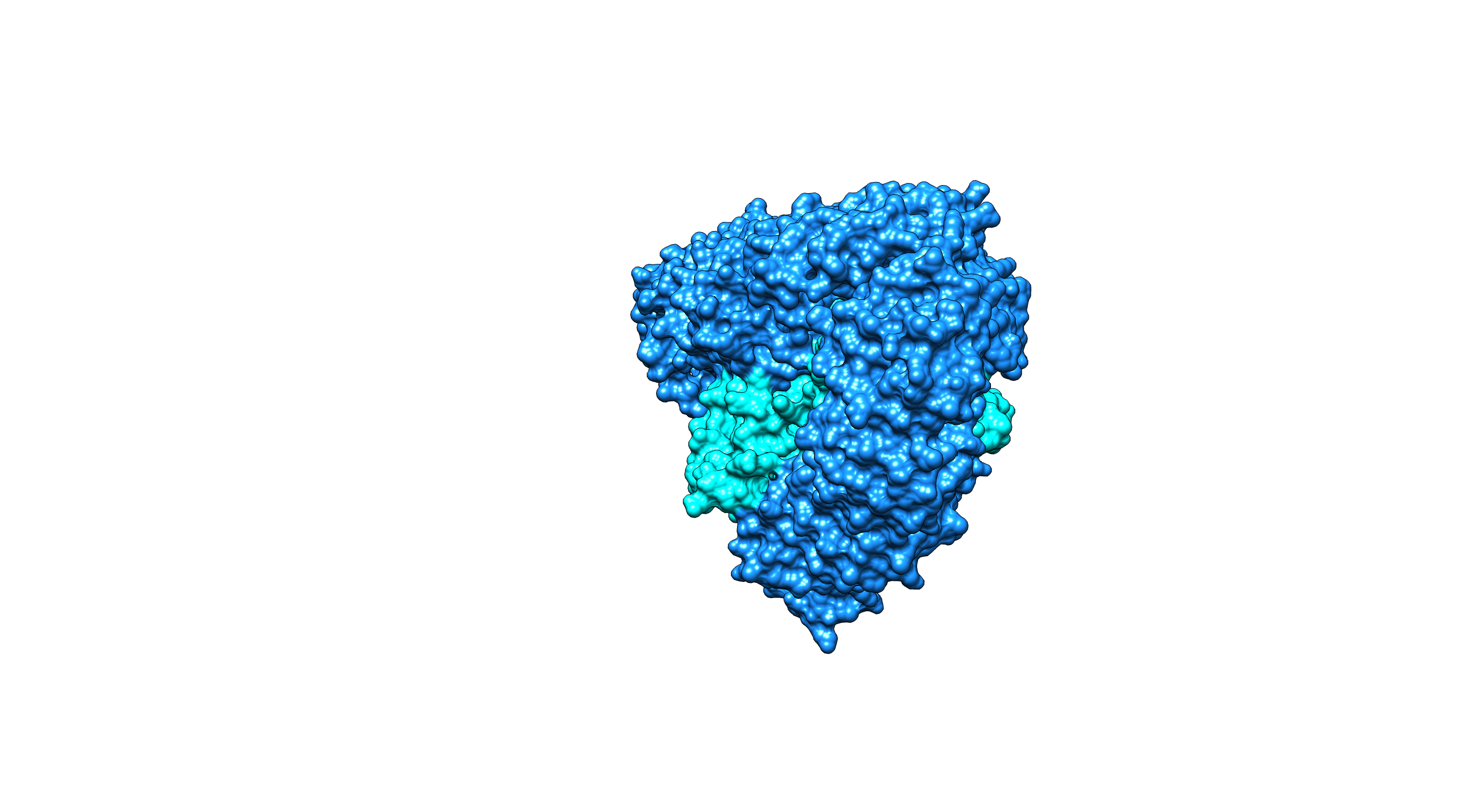

### imagespacefillinh.png

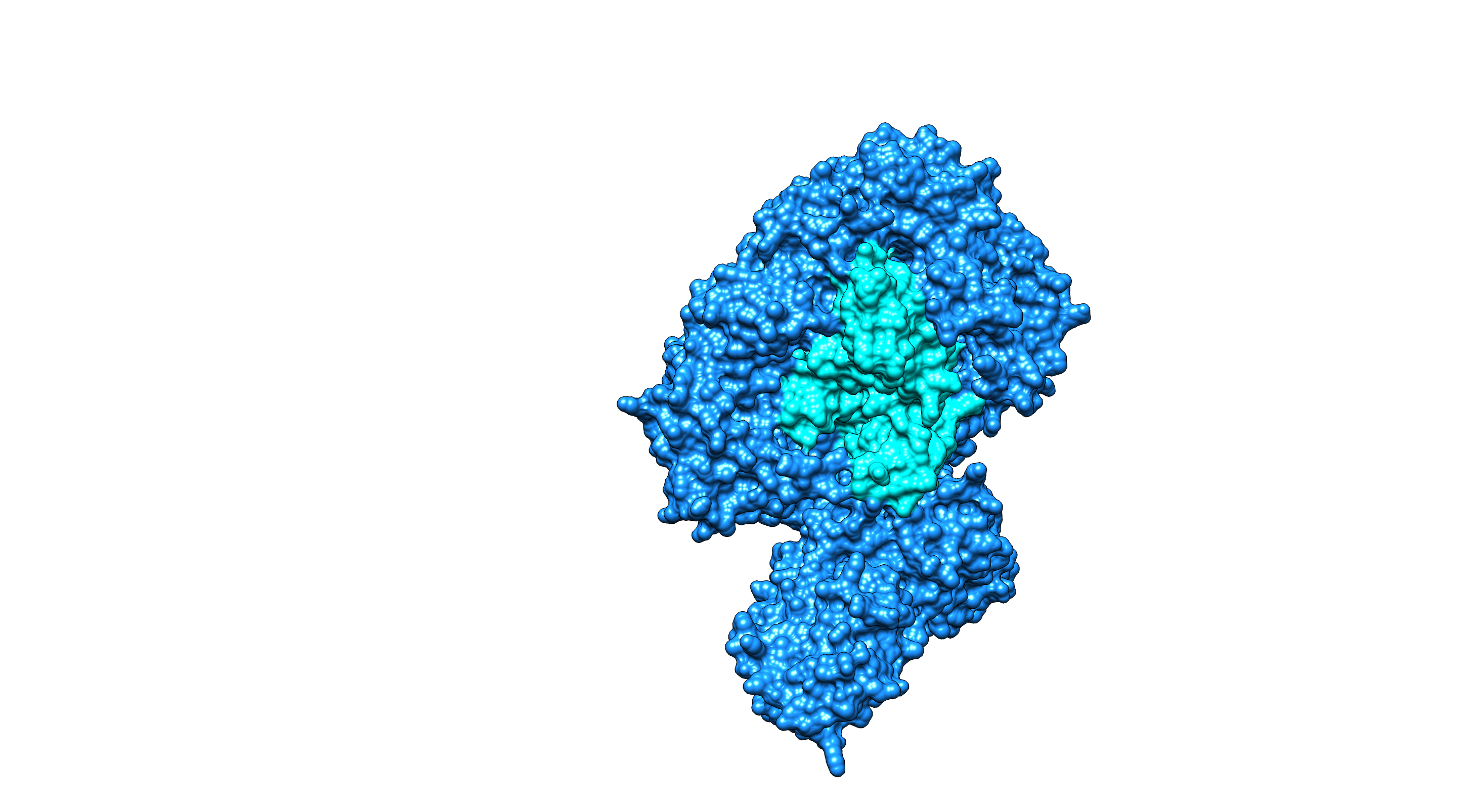

### lambdap42.png

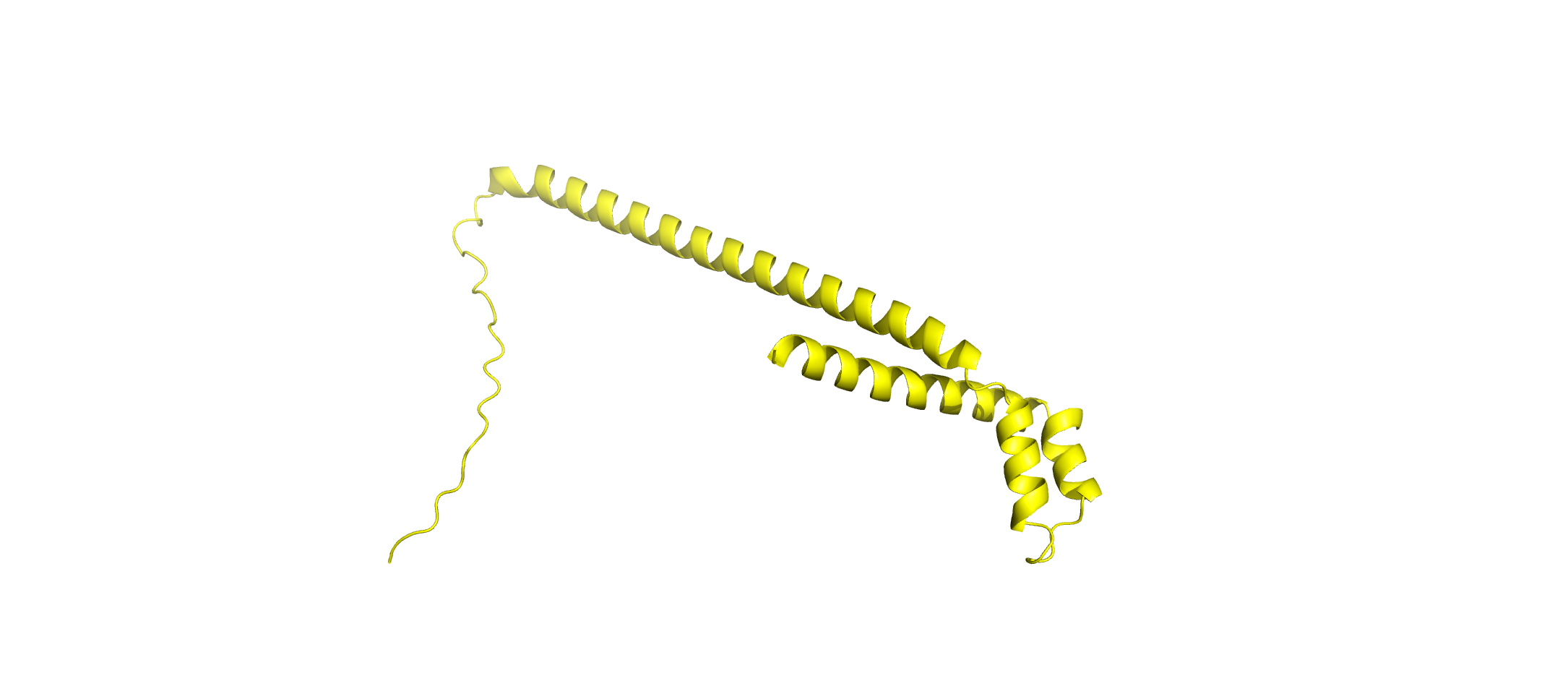

### lambdap88 brown.png

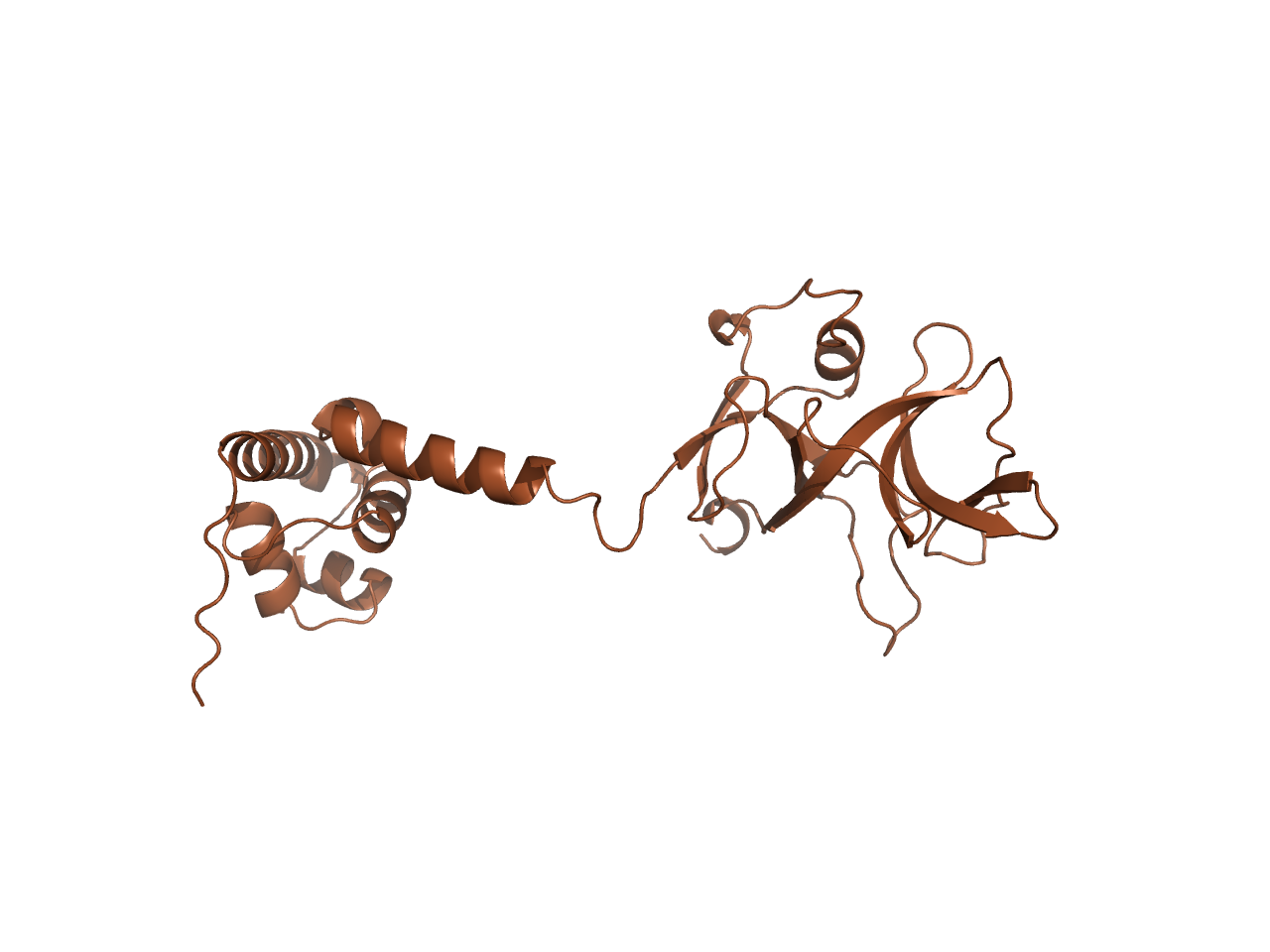

### lambdap88.png

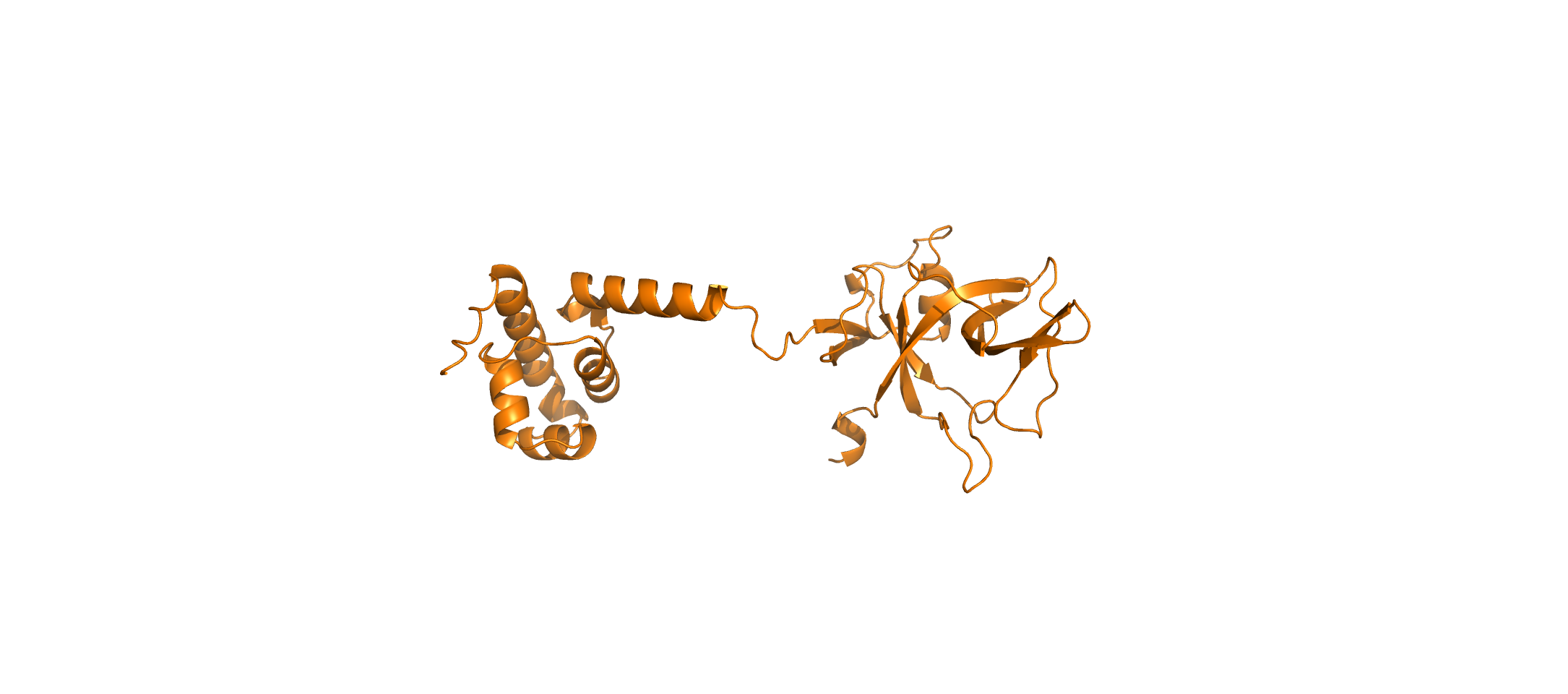

### multicolorimage90degrees.png

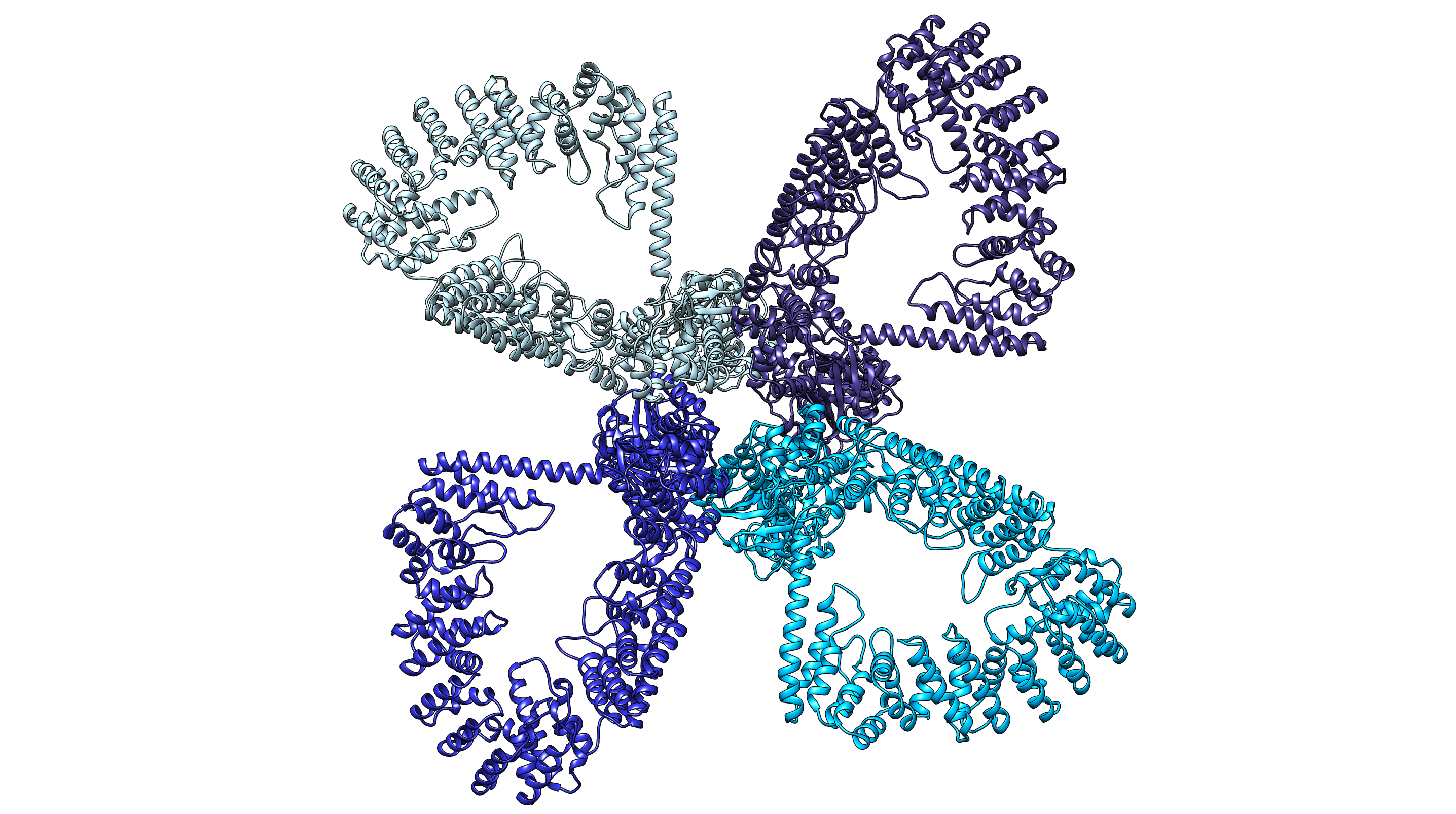

### multicolorimage.png

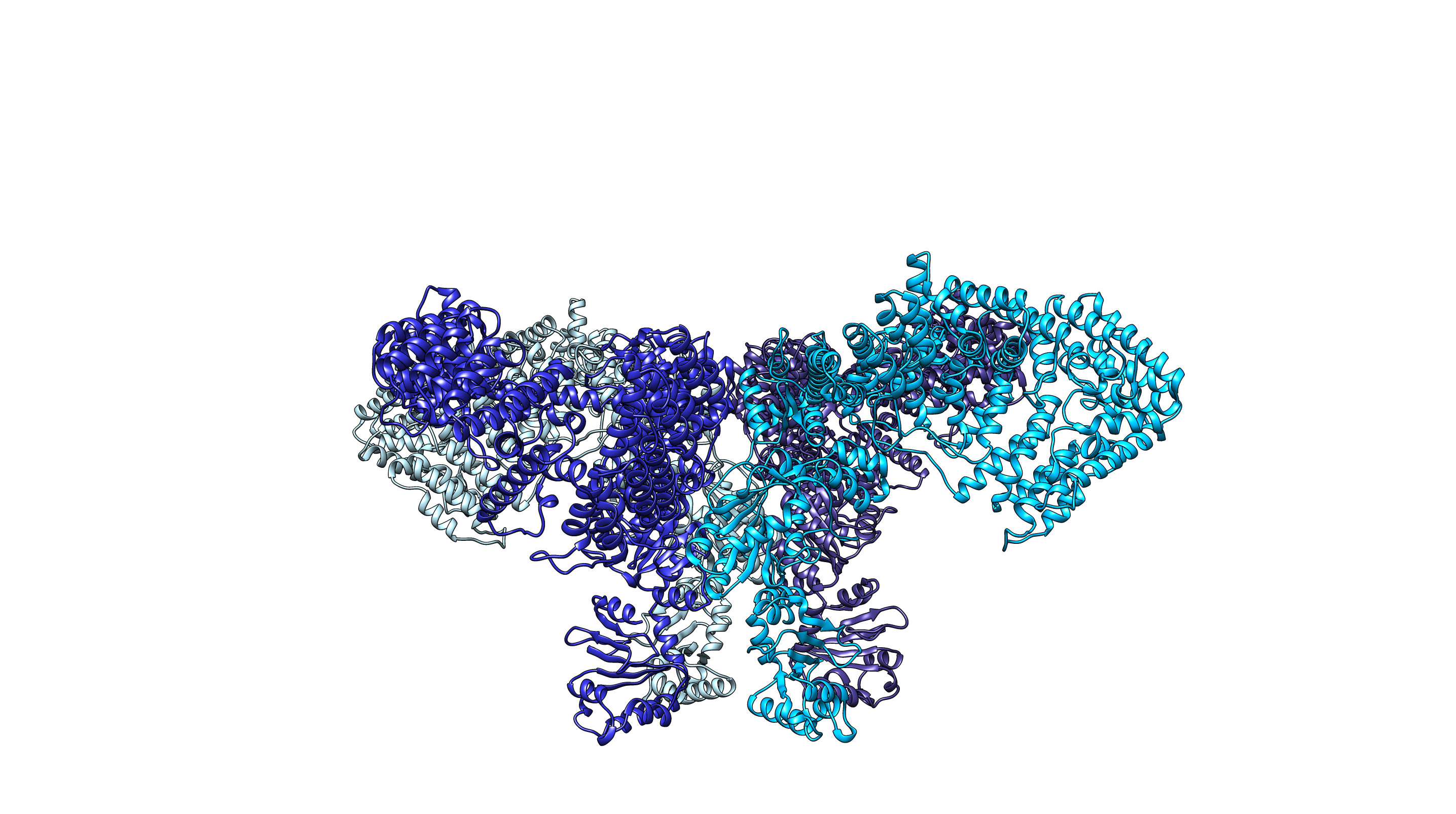

### pdl4b.png

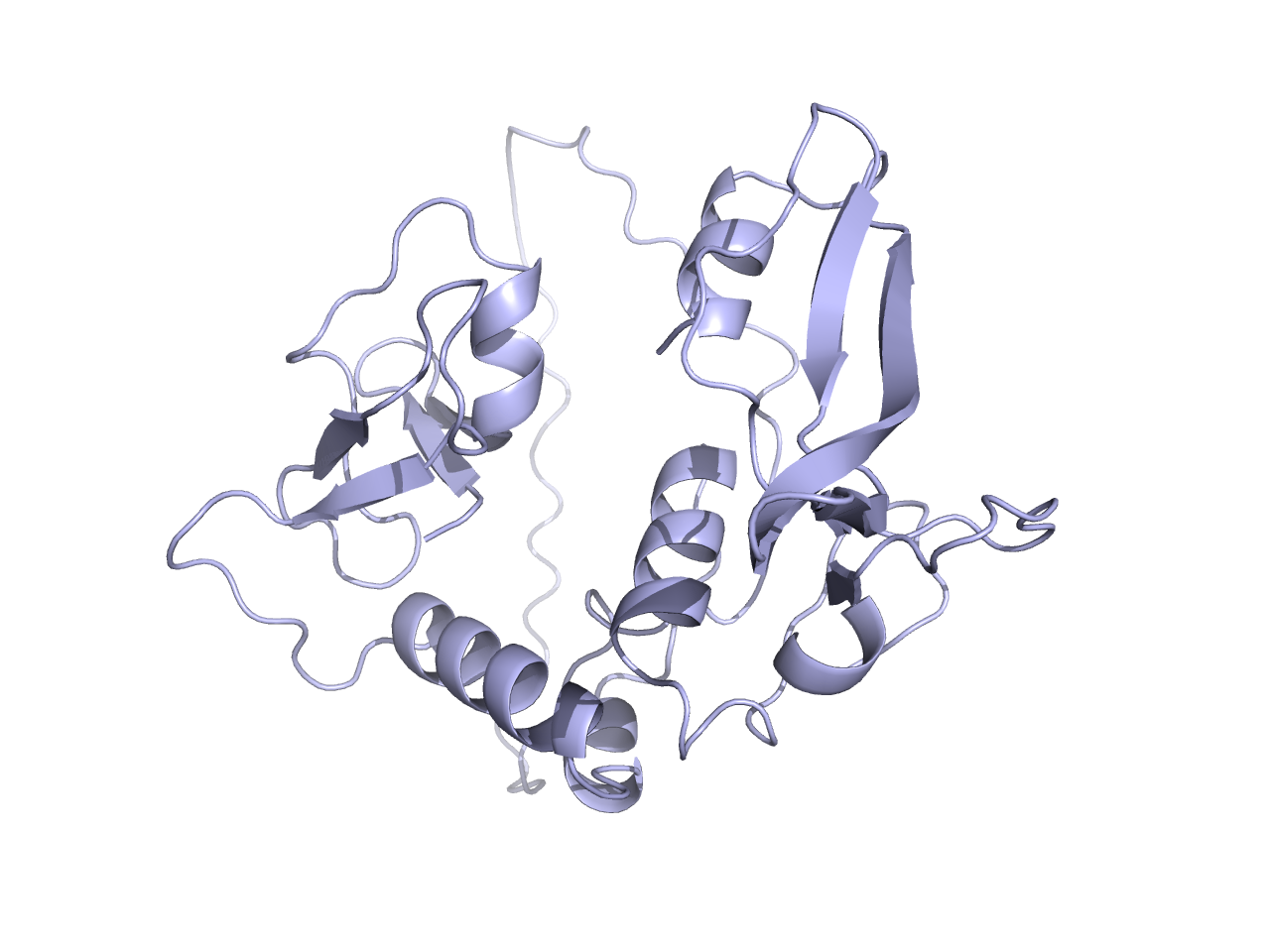

### rainbow90 to left.png

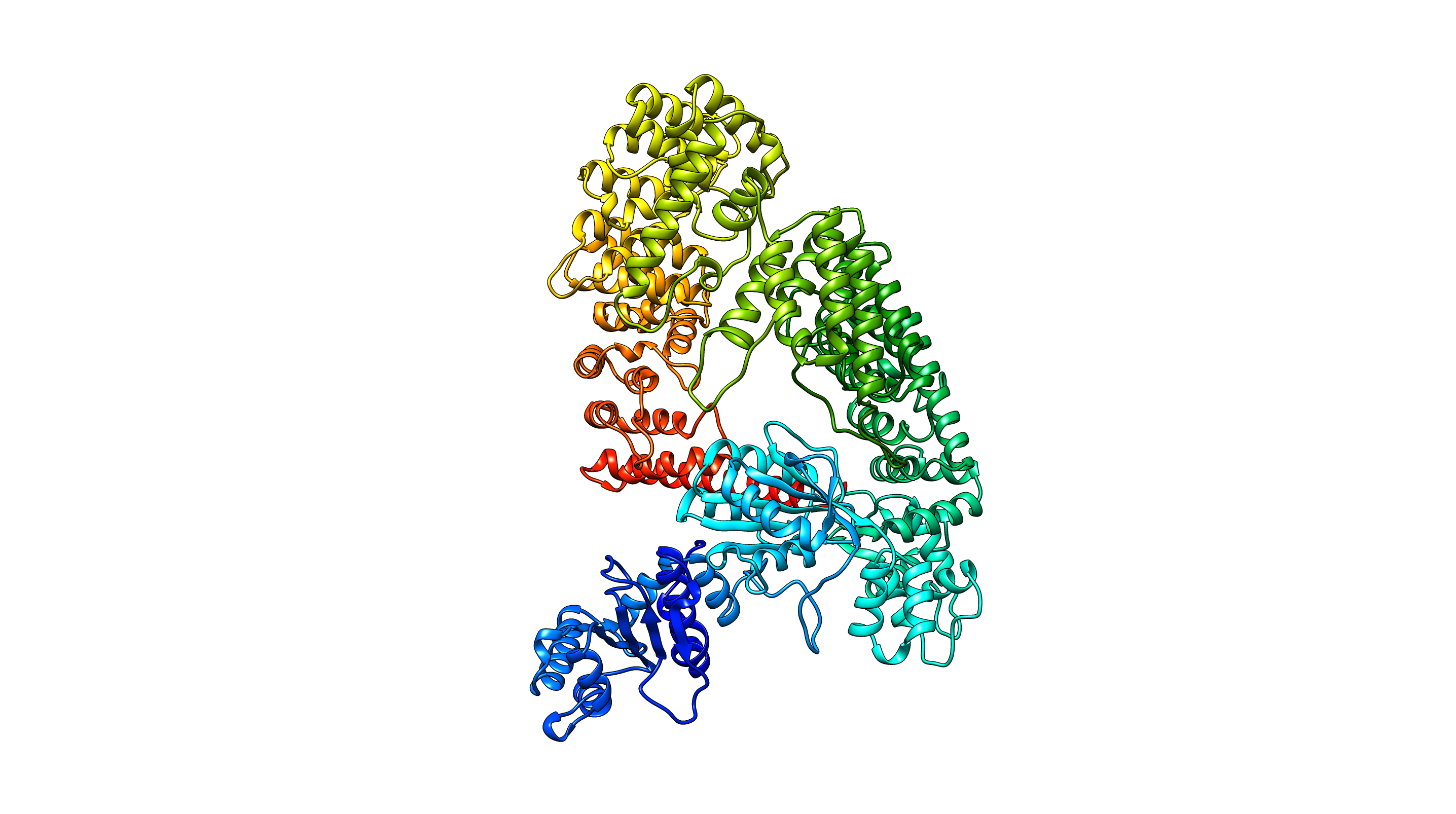

### rainbow tetramer90.png

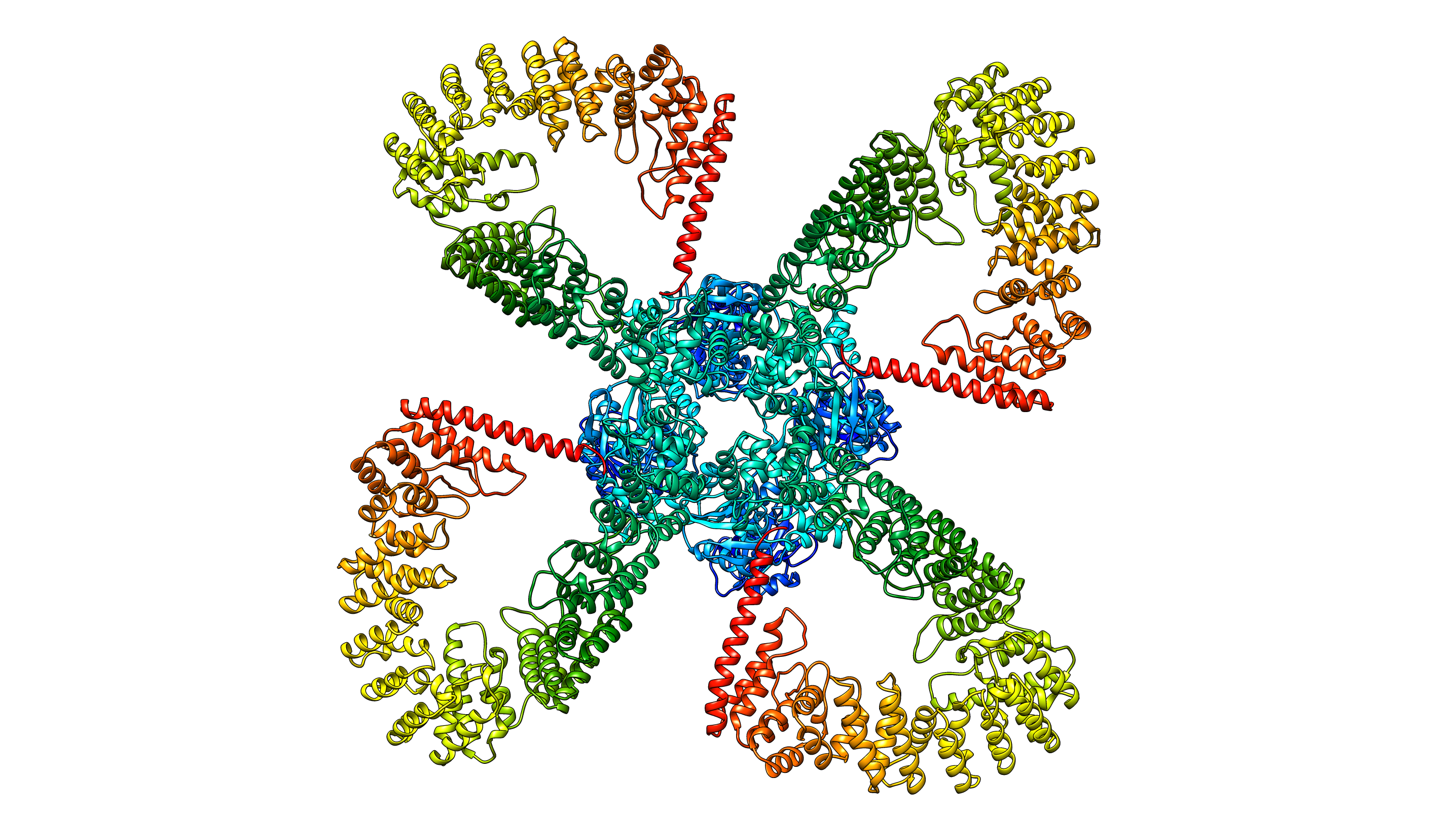

### rainbow tetramer 90 to left.png

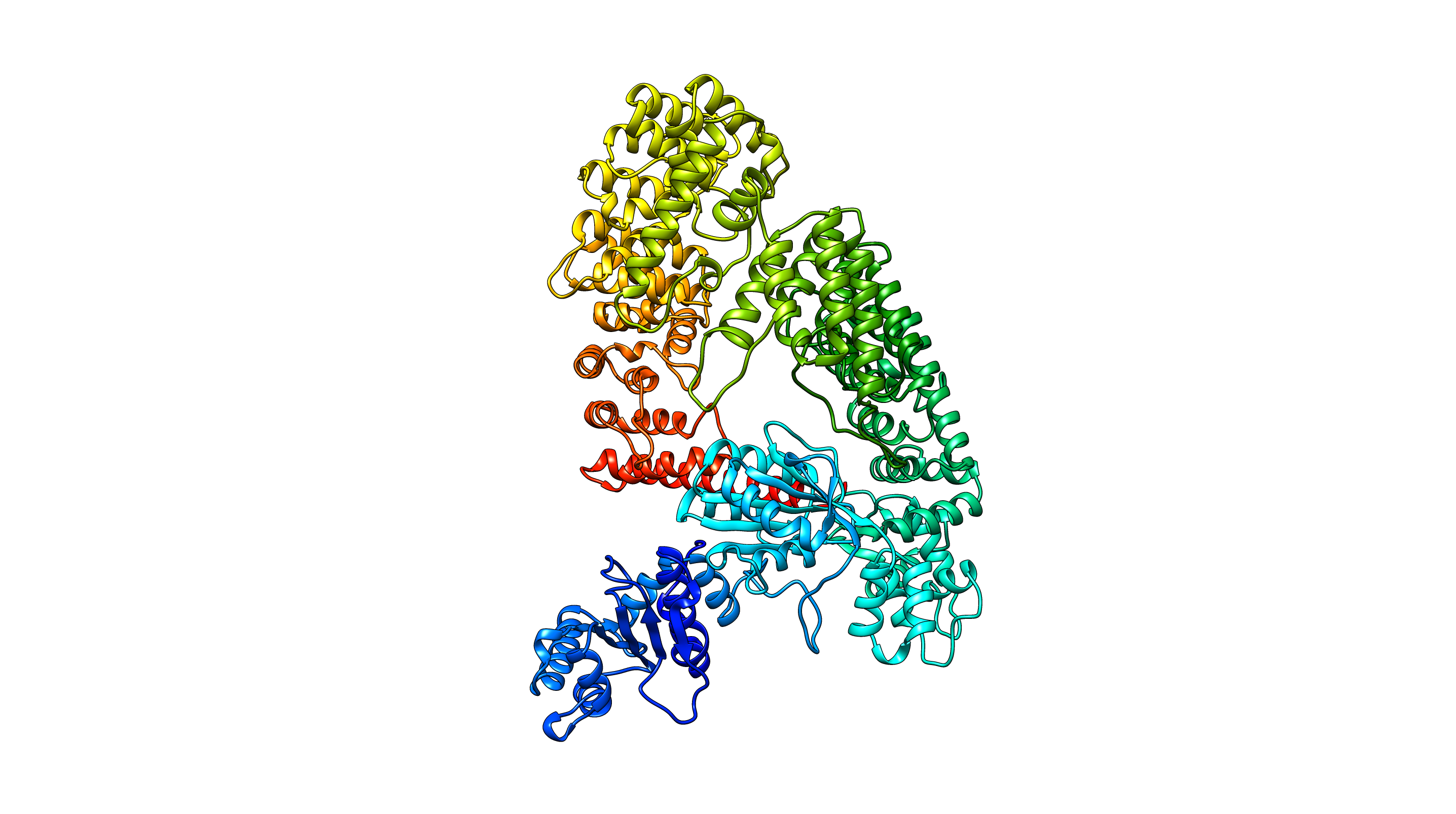

### rainbow tetramer.png

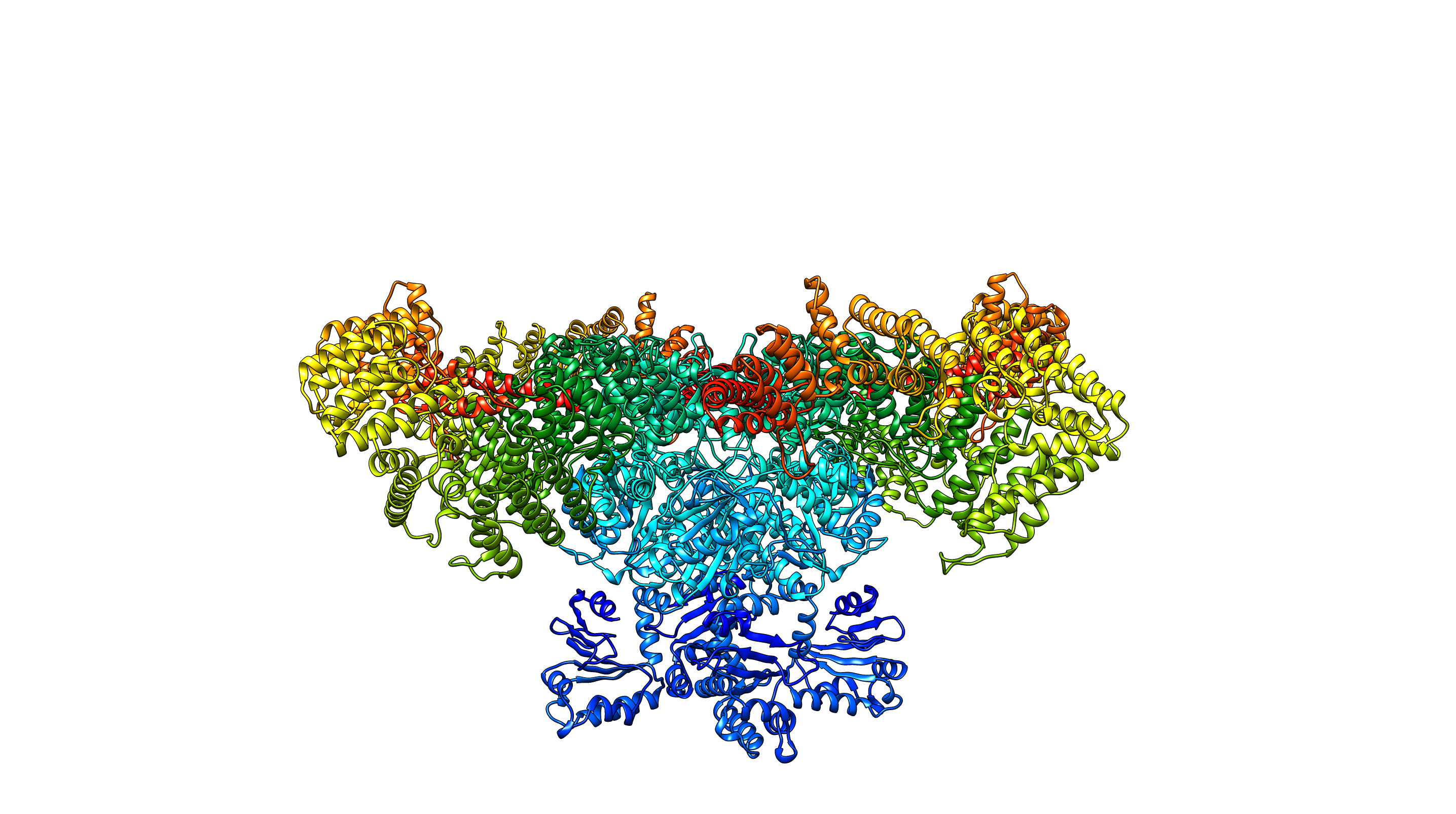

### rainbow.png

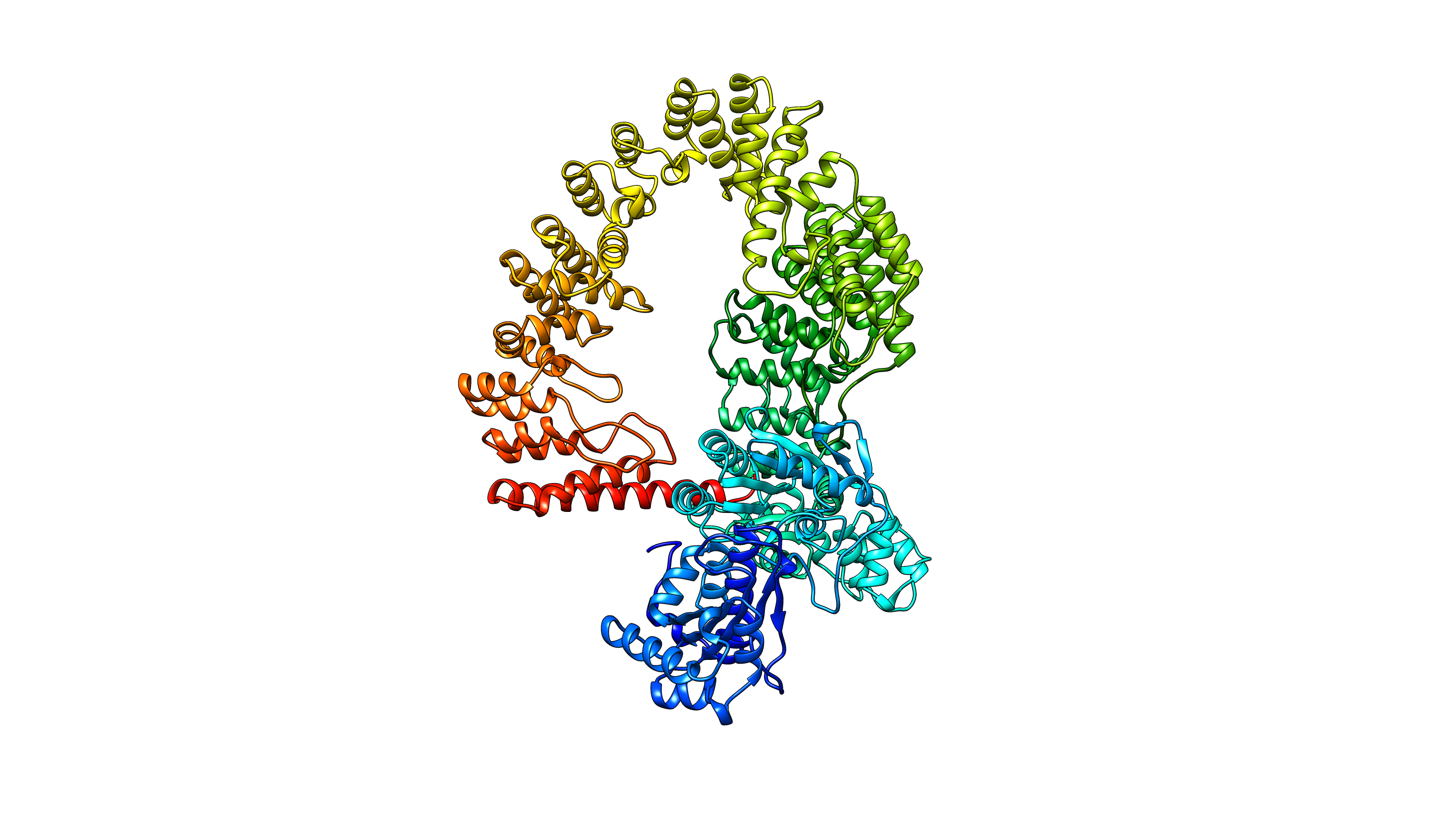

### Screenshot 2025-04-25 at 1.56.40ΓÇ»PM.png

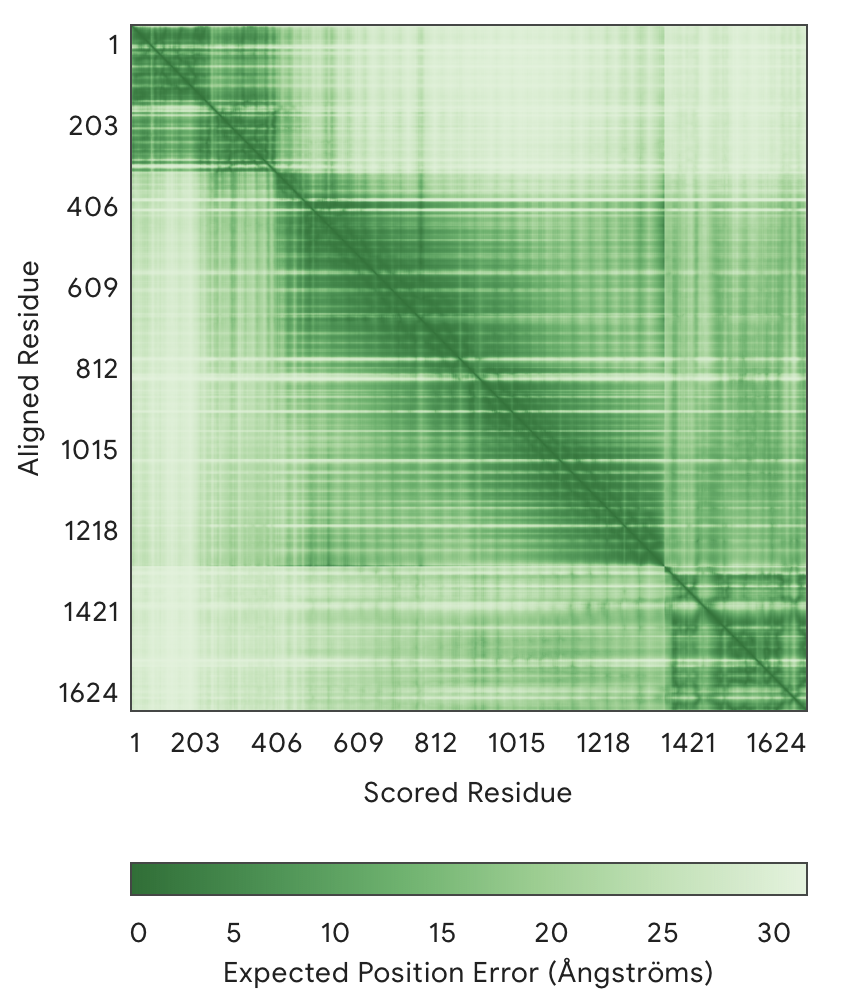

### Screenshot 2025-04-25 at 2.05.07ΓÇ»PM.png

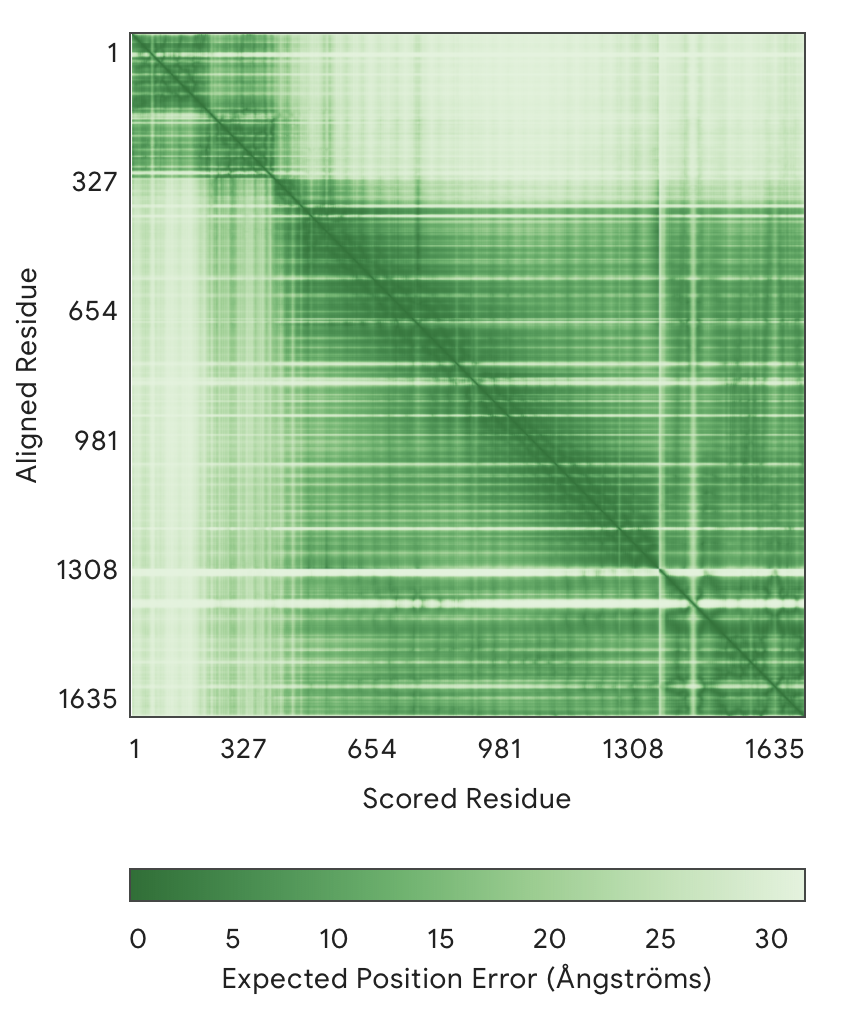

### T7gp14.png

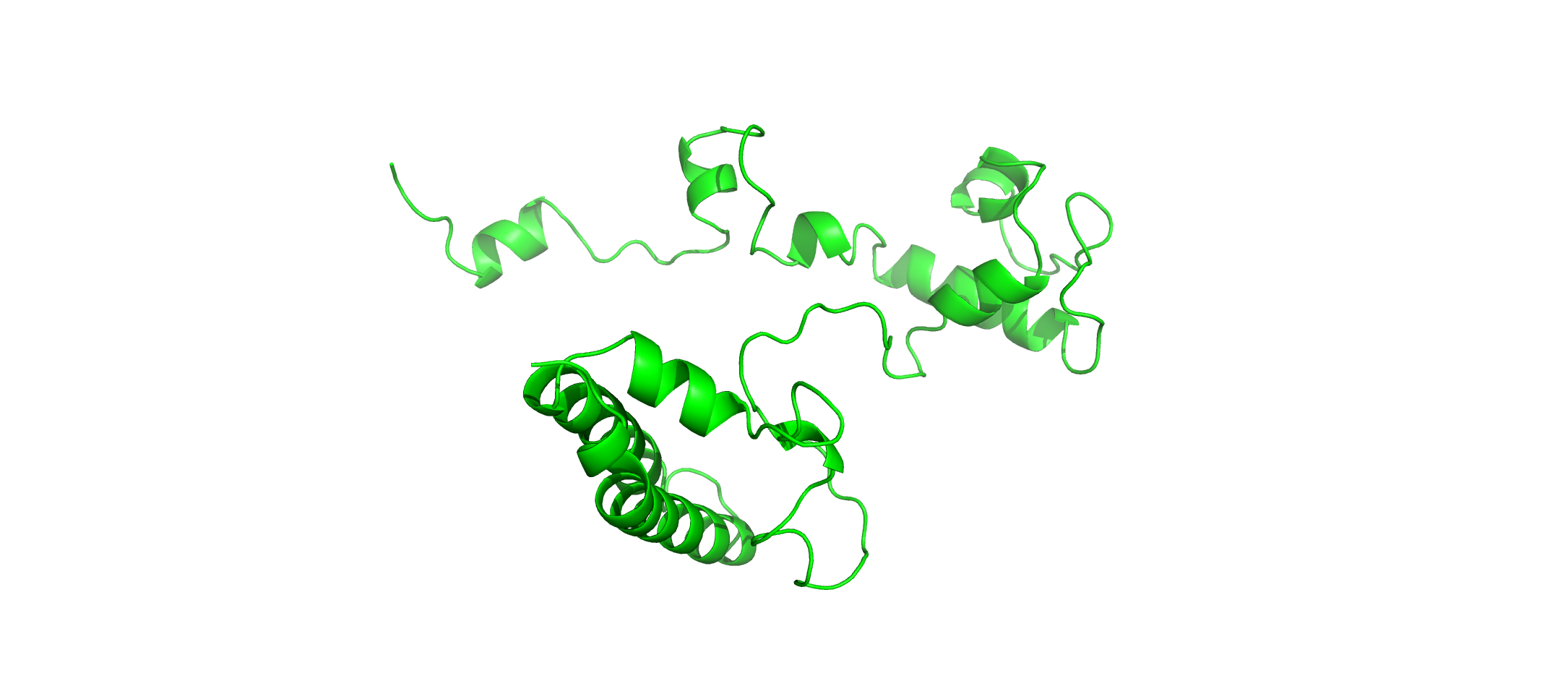

### T7p48.png

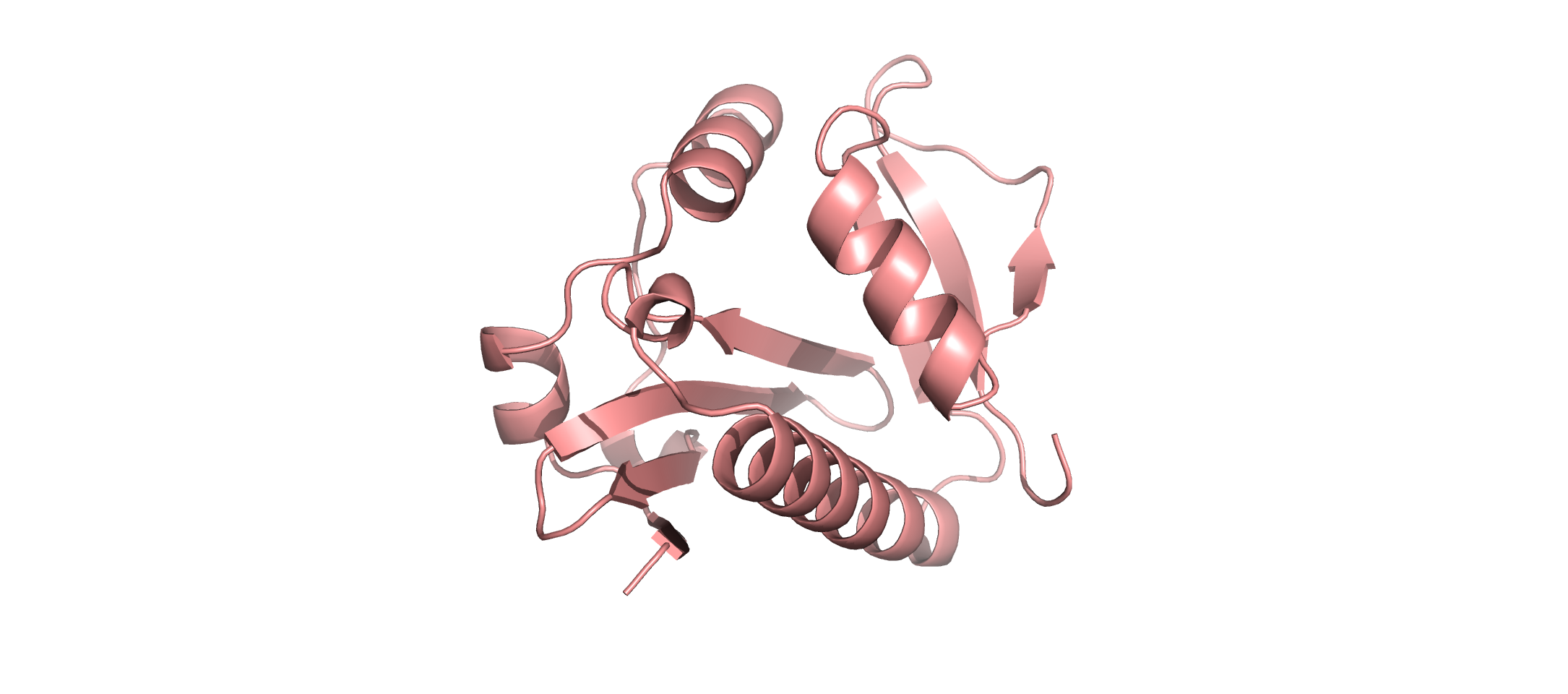

### tetramer25deltete.png

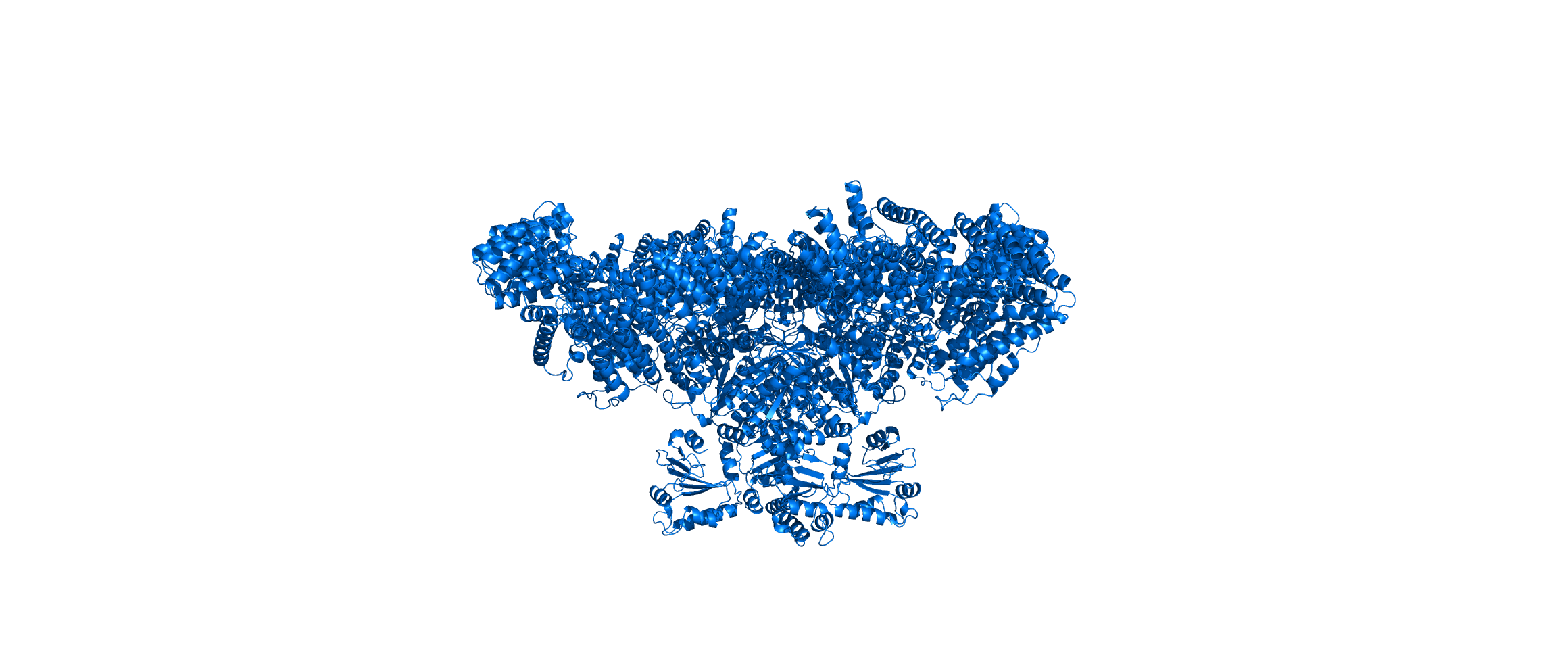
